# Inference and reconstruction of the heimdallarchaeial ancestry of eukaryotes

**DOI:** 10.1101/2023.03.07.531504

**Authors:** Laura Eme, Daniel Tamarit, Eva F. Caceres, Courtney W. Stairs, Valerie De Anda, Max E. Schön, Kiley W. Seitz, Nina Dombrowski, William H. Lewis, Felix Homa, Jimmy H. Saw, Jonathan Lombard, Takuro Nunoura, Wen-Jun Li, Zheng-Shuang Hua, Lin-Xing Chen, Jillian F. Banfield, Emily St John, Anna-Louise Reysenbach, Matthew B. Stott, Andreas Schramm, Kasper U. Kjeldsen, Andreas P. Teske, Brett J. Baker, Thijs J. G. Ettema

**Author notes:** Equal contribution (Laura Eme, Daniel Tamarit, Eva F. Caceres). Theoretical Biology and Bioinformatics, Department of Biology, Faculty of Science, Utrecht University, Padualaan 8, 3584CH Utrecht, The Netherlands (Daniel Tamarit). Department of Biology, Lund University, Sölvegatan 35, 223 62 Lund, Sweden. Structural and Computational Biology, European Molecular Biology Laboratory, Meyerhofstraße 1, 69117 Heidelberg, Germany. NIOZ, Royal Netherlands Institute for Sea Research, Department of Marine Microbiology and Biogeochemistry; AB Den Burg, The Netherlands. Department of Biochemistry, University of Cambridge, Cambridge, CB2 1QW, UK. Department of Biological Sciences, The George Washington University, Washington, DC, USA.

## Abstract

In the ongoing debates about eukaryogenesis, the series of evolutionary events leading to the emergence of the eukaryotic cell from prokaryotic ancestors, members of the Asgard archaea play a key role as the closest archaeal relatives of eukaryotes. However, the nature and phylogenetic identity of the last common ancestor of Asgard archaea and eukaryotes remain unresolved. Here, we analyze distinct phylogenetic marker datasets of an expanded genomic sampling of Asgard archaea and evaluate competing evolutionary scenarios using state-of-the-art phylogenomic approaches. We find that eukaryotes are placed, with high confidence, as a well-nested clade within Asgard archaea, as a sister lineage to Hodarchaeales, a newly proposed order within Heimdallarchaeia. Using sophisticated gene tree/species tree reconciliation approaches, we show that, in analogy to the evolution of eukaryotic genomes, genome evolution in Asgard archaea involved significantly more gene duplication and fewer gene loss events compared to other archaea. Finally, we infer that the last common ancestor of Asgard archaea likely was a thermophilic chemolithotroph, and that the lineage from which eukaryotes evolved adapted to mesophilic conditions and acquired the genetic potential to support a heterotrophic lifestyle. Our work provides key insights into the prokaryote-to-eukaryote transition and the platform for the emergence of cellular complexity in eukaryotic cells.

## Main

Understanding how complex eukaryotic cells emerged from prokaryotic ancestors represents a major challenge in biology^1, 2^. A main point of contention in refining eukaryogenesis scenarios revolves around the exact phylogenetic relationship between Archaea and eukaryotes. The use of phylogenomic approaches with improved models of sequence evolution combined with a much-improved archaeal taxon sampling – progressively unveiled by metagenomics – has recently yielded strong support for the “two-domain” tree of life, in which the eukaryotic clade branches from within Archaea^3–8^. The discovery of the first Lokiarchaeia genome provided additional evidence for the two-domain topology since this lineage was shown to represent, at the time, the closest relative of eukaryotes in phylogenomic analyses^9^. Moreover, Lokiarchaeia genomes were found to uniquely contain many genes encoding eukaryotic signature proteins (ESPs) –proteins involved in hallmark complex processes of the eukaryotic cell–, more so than any other prokaryotic lineage. The subsequent identification and analyses of several diverse relatives of Lokiarchaeia, together forming the Asgard archaea superphylum, confirmed that Asgard archaea represented the closest archaeal relatives of eukaryotes^2, 9, 10^. Their exact evolutionary relationship to eukaryotes, however, remained unresolved: it has been unclear whether eukaryotes evolved from *within* Asgard archaea, or if they represented their sister-lineage^10^. Furthermore, two studies questioned this view of the tree of life altogether, suggesting that Asgard archaea represent a deep-branching Euryarchaeota-related clade^11, 12^, and that, in accordance with the “three-domain” tree, eukaryotes represent a sister group to all Archaea, although this was challenged^13, 14^. A follow-up study that included an expanded taxonomic sampling of Asgard archaeal genome data failed to resolve the phylogenetic position of eukaryotes in the tree of life^15^.

Here, we expand the genomic diversity of Asgard archaea by generating 63 novel Asgard metagenomic-assembled genomes (MAGs) from samples from 11 locations around the world. By analyzing the improved genomic sampling of Asgard archaea using state-of-the-art phylogenomics, including recently developed gene tree/species tree reconciliation approaches for ancestral genome content reconstruction, we firmly place eukaryotes nested within the Asgard archaea. By revealing key features regarding the identity, nature and physiology of the last Asgard archaea and eukaryotes common ancestor (LAECA), our results represent important, thus far missing pieces of the elusive eukaryogenesis puzzle.

### Expanded Asgard archaea genomic diversity

To increase the genomic diversity of Asgard archaea, we sampled aquatic sediments and hydrothermal deposits from eleven geographically distinct sites (Supplementary Table 1, Supplementary Figure 1). After extraction and sequencing of total environmental DNA, we assembled and binned metagenomic reads into MAGs. Of these MAGs, 63 were found to belong to the Asgard archaea superphylum, with estimated median completeness and redundancy of 83% and 4.2%, respectively (Supplementary Table 1). To assess the genomic diversity in this dataset, we reconstructed a phylogeny of ribosomal proteins encoded in a conserved 15-ribosomal protein (RP15) gene cluster^16^ from these MAGs, and all publicly available Asgard archaea assemblies (retrieved June 29^th^, 2021; Figure 1). These analyses expand the genomic sampling across previously described major Asgard archaea clades (i.e., Loki-, Thor-, Heimdall-, Odin-, Hel-, Hermod-, Sif-, Jord- and Baldrarchaeia^9, 10, 15, 17, 18^) and recover a previously undescribed clade of high taxonomic rank (Candidatus Asgardarchaeia; see Ext. Data Fig. 1 and Supplementary Information for proposed uniformization of Asgard archaea taxonomic classification that will be adhered to throughout the present manuscript). We observed that the median estimated Asgard archaeal genome size (3.8 Mega basepairs (Mbp)) is considerably larger than those of representative genomes from TACK archaea and Euryarchaeota (median=1.8 Mbp for both) and DPANN archaea (median=1.2 Mbp) (Supplementary Table 1). Among Asgard archaea, Odinarchaeia display the smallest genomes (median=1.4 Mb), while Loki- and Helarchaeales contain the largest (median=4.3 Mbp for both). Unlike other major Asgard archaeal clades, Heimdallarchaeia possess a wide range of genome sizes, spanning from 1.6 to 7.4 Mbp (median=3.5 Mbp). Indeed, this large class contains five clades with diverse features. These include Njordarchaeales (median genome size=2.4 Mbp) followed by Kariarchaeaceae (median genome size=2.7 Mbp), Gerdarchaeales (median genome size=3.4 Mbp), Heimdallarchaeaceae (median genome size=3.7 Mbp), and finally Hodarchaeales (median genome size=5.1 Mbp). The smallest heimdallarchaeial genome corresponds to the only Asgard archaeal MAG recovered from a marine surface water metagenome (Heimdallarchaeota archaeon RS678)^19^, in agreement with reduced genome sizes typically observed among prokaryotic plankton of the euphotic zone^20^ .

**Figure 1.**
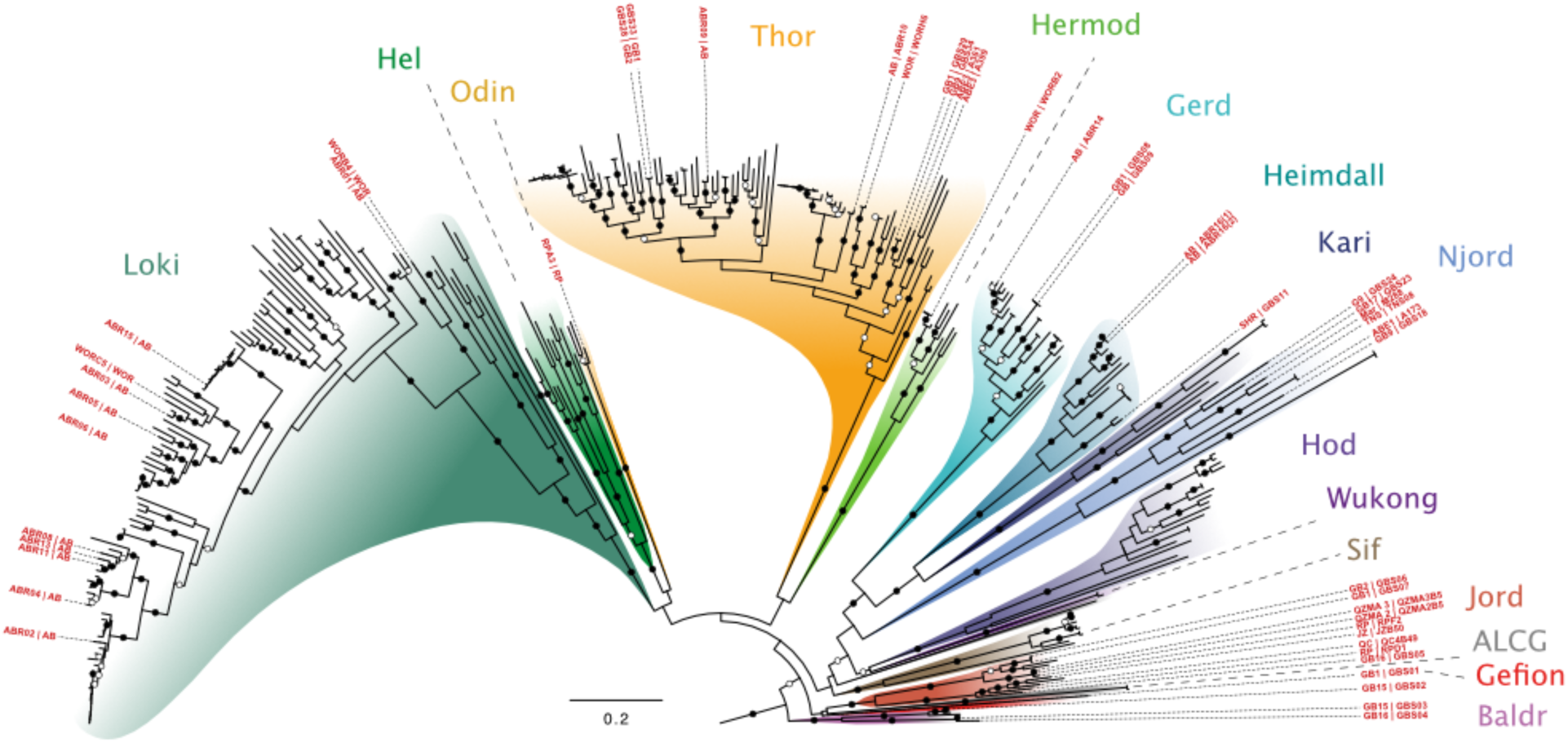
Phylogenomic analysis of 15 concatenated ribosomal proteins expands Asgard archaea diversity. Maximum-likelihood tree (IQ-TREE, WAG+C60+R4+F+PMSF model) of concatenated protein sequences from at least five genes, encoded on a single contig, of a 15 conserved ribosomal protein (RP15) gene cluster retrieved from publicly available and newly reported Asgard archaeal MAGs. Bootstrap support (100 pseudo-replicates) is indicated by circles at branches, with filled and open circles representing 90% and 70% support, respectively. Leaf names indicate geographical source and isolate name (inner and outer label, respectively) for the MAGs reported in this study. Scale bar denotes the average number of substitutions per site. Abbreviations: AB: Aarhus Bay (Denmark); ABE: ABE vent field, Eastern Lau Spreading Center; ALCG: Asgard Lake Cootharaba Group; CR: Colorado River (USA); GB: Guaymas Basin (Mexico); JZ: Jinze (China); LC: Loki’s castle; LCB: Lower Culex Basin (USA); Mar: Mariner vent field, Eastern Lau Spreading Center; NSCS: Northern South China Sea; OWC: Old Woman Creek (USA); QC: QuCai village (China); QZM: QuZhuoMu village (China); RP: Radiata Pool (New Zealand); RS: Red sea; SHR: South Hydrate Ridge; TNS: Taketomi Island (Japan); WOR: White Oak River (USA).

### Identification of phylogenetic conflict

Inferring deep evolutionary relationships in the tree of life is considered one of the hardest problems in phylogenetics. To interrogate the evolutionary relationships within the present set of Asgard archaeal phyla, and between Asgard archaea and eukaryotes, we performed an exhaustive range of sophisticated phylogenomic analyses. We analyzed a preexisting marker dataset comprising 56 concatenated ribosomal protein sequences (RP56)^9, 10^ for a phylogenetically diverse set of 331 archaeal (175 Asgard archaea, 41 DPANN, 43 Euryarchaeota, and 72 TACK archaea representatives), and 14 eukaryotic taxa (see Supplementary Table 2). Of note, the inclusion of an expanded diversity of 12 new Korarchaeota MAGs among these TACK archaea considerably affected phylogenomic analyses (see below). Initial maximum-likelihood (ML) phylogenetic inference based on this RP56 dataset confirmed the existence of 12 major Asgard archaeal clades of high taxonomic rank (Supplementary Figure 2). These include the previously described Loki-, Odin-, Heimdall-, Thor-^9, 10^, Helarchaeia^10^, for which we here present 36 new genomes, and the recently proposed Sif-^18^, Hermod-^17^, Jord-^21^, Wukong-^15^ and Baldrarchaeia^15^, for most of which we also identified new near-complete MAGs. Finally, we identified 15 MAGs representing the recently described Njordarchaeales^22^ (which we show below is a divergent candidate order within Heimdallarchaeia), and a single MAG representing a new candidate class, Asgardarchaeia (which will be discussed in a separate manuscript; Tamarit et al, in prep) (Figure 1). Importantly, careful inspection of the obtained RP56 tree uncovered a potential artefact: Njordarchaeales, considered *bona fide* Asgard archaea based on the presence of many encoded ‘typical’ Asgard-like ESPs^10^, were found to branch outside of the Asgard archaea, at the base of the TACK superphylum and as a sister lineage to Korarchaeota in the RP56 tree. In addition, eukaryotes were found to branch at the base of the clade formed by Korarchaeota and Njordarchaeales, although with weak support. Hereafter, we focused on disentangling the historically correct phylogenetic signal from noise and artefacts.

### Alternative phylogenomic markers

Despite often being used in phylogenomic analyses, ribosomal proteins have been suggested to contribute to phylogenetic artefacts due to inherent compositional sequence biases^23, 24^. Additionally, considering the inconsistency of the obtained placement of eukaryotes compared to previous analyses, the incoherent placement of Njordarchaeales, and the presence of long branches at the base of both of these clades in the RP56 tree, we sought to use an alternative phylogenetic marker set to obtain a stable Asgard archaeal species tree, and to further investigate the phylogenetic position of eukaryotes. We constructed an independent ‘new marker’ dataset comprising 57 proteins of archaeal origin in eukaryotes (NM57 dataset; see Methods). The NM57 proteins are mostly involved in diverse informational, metabolic, and cellular processes, but do not include ribosomal proteins (Supplementary Table 2). Besides being longer, and hence putatively more phylogenetically informative compared to the RP56 markers, the broader functional distribution of NM57 markers is less likely to cause phylogenetic reconstruction artefacts induced by strong co-evolution between proteins – something that is to be expected for functionally and structurally cohesive ribosomal proteins^25^. Indeed, in case co-evolving protein sequences are compositionally biased, and hence violate evolutionary model assumptions of fixed composition over species, their concatenation is expected to strengthen the artefactual, non-phylogenetic signal and the statistical support for incorrect relationships^26^. We thus decided to independently evaluate the concatenated NM57 and RP56 marker datasets for downstream phylogenomic analyses. We observed that ML phylogenomic analyses of the NM57 dataset did not only recover Njordarchaeales as *bona fide* Asgard archaea, they were also placed as the closest relatives of eukaryotes (BS=98%; Supplementary Figure 3), as was proposed in a recent analysis^22^. To investigate the underlying causes for the contradicting results between the NM57 and RP56 datasets, we first assessed the effect of taxon sampling on phylogenetic reconstructions by removing eukaryotic and/or DPANN and/or Korarchaeota sequences from the alignments, for two main reasons: (1) eukaryotes and DPANN archaea represent long-branching clades potentially inducing long branch attraction (LBA) artefacts; and (2) we wanted to investigate the effects of removing eukaryotes and Korarchaeota, which were the sister lineages of Njordarchaeales in the NM57 and RP56 phylogenetic analyses, respectively. Following this, we recoded the alignments into 4 states (using SR4-recoding^27^) to ameliorate potential phylogenetic artefacts arising from model misspecification at mutationally saturated or compositionally biased sites^14, 28–30^. Further, with a similar goal, we applied a fast-evolving site removal (FSR) procedure to the concatenated datasets, since fast-evolving sites are often mutationally saturated. We performed phylogenetic analyses of the above-mentioned datasets in both ML and Bayesian Inference (BI) frameworks, under sophisticated evolutionary models that account for sequence heterogeneity in the substitution process across sites (mixture models; Supplementary Table 2).

Phylogenomic analyses of the above-mentioned combinations of taxon sampling, data treatments and phylogenetic frameworks revealed that Njordarchaeales are artefactually attracted to Korarchaeota in RP56 datasets (Supplementary Information). This attraction is likely caused by the high compositional similarity of njord- and korarchaeal RP56 ribosomal protein sequences, which is probably linked to their shared hyperthermophilic lifestyle (Supplementary Figures 4-6). Analyses of RP56 datasets from which Korarchaeota were removed, recovered Njordarchaeales as an order at the base of or within Heimdallarchaeia (Supplementary Figure 7), consistent with phylogenomic analyses of the NM57 dataset that included Korarchaeota (Supplementary Figure 3). Next, in our efforts to resolve the phylogenetic placement of eukaryotes, we initially performed phylogenomic analyses on variations of the RP56 and NM57 datasets (Supplementary Table 2 and Discussion). However, since compared to the RP56 dataset, the NM57 dataset is larger and less compositionally biased, and is thus expected to have retained a stronger and more congruent phylogenetic signal, we focused the rest of our study on this more reliable dataset.

### Eukarya represent a well-nested clade within Heimdallarchaeia

Subsequent phylogenetic analyses of untreated NM57 datasets with various taxon sampling variations recovered eukaryotes as sister-clade to Njordarchaeales in ML analyses (e.g., Supplementary Figure 3, Supplementary Table 2 and Supplementary information). However, ML analyses of the SR4-recoded datasets retrieved a complex phylogenetic signal, as in some cases eukaryotes were placed at the base of all Heimdallarchaeia (including Njordarchaeales) and Wukongarchaeia. This strongly suggests that the previously observed phylogenetic affiliation between Njordarchaeales and eukaryotes could represent an artefact. Furthermore, when both SR4-recoding and FSR treatments were combined, eukaryotes were nested within Heimdallarchaeia, as sister-group to the order Hodarchaeales (Figure 2; Supplementary Figure 8), and this position was supported by ML analyses of NM57 datasets across all taxon selection variations (removing DPANN archaea, and/or Korarchaeota and/or Njordarchaeales). Congruently, the monophyly of eukaryotes and Hodarchaeales was systematically recovered by BI of recoded datasets (both with and without FSR; Figure 2, Supplementary Table 2). In addition, the position of Njordarchaeales shifted during these analyses, moving from a deep position at the base of Heimdallarchaeia and Wukongarchaeia, to a more nested position forming a clade with Gerdarchaeales and Kari- and Heimdallarchaeaceae (Supplementary Discussion). This shift is observed in both the NM57 and the RP56 datasets analyses when SR4-recoding and FSR was combined (Supplementary Figures 9-10), supporting that Njordarchaeales represent a divergent order-level lineage of Heimdallarchaeia.

**Figure 2.**
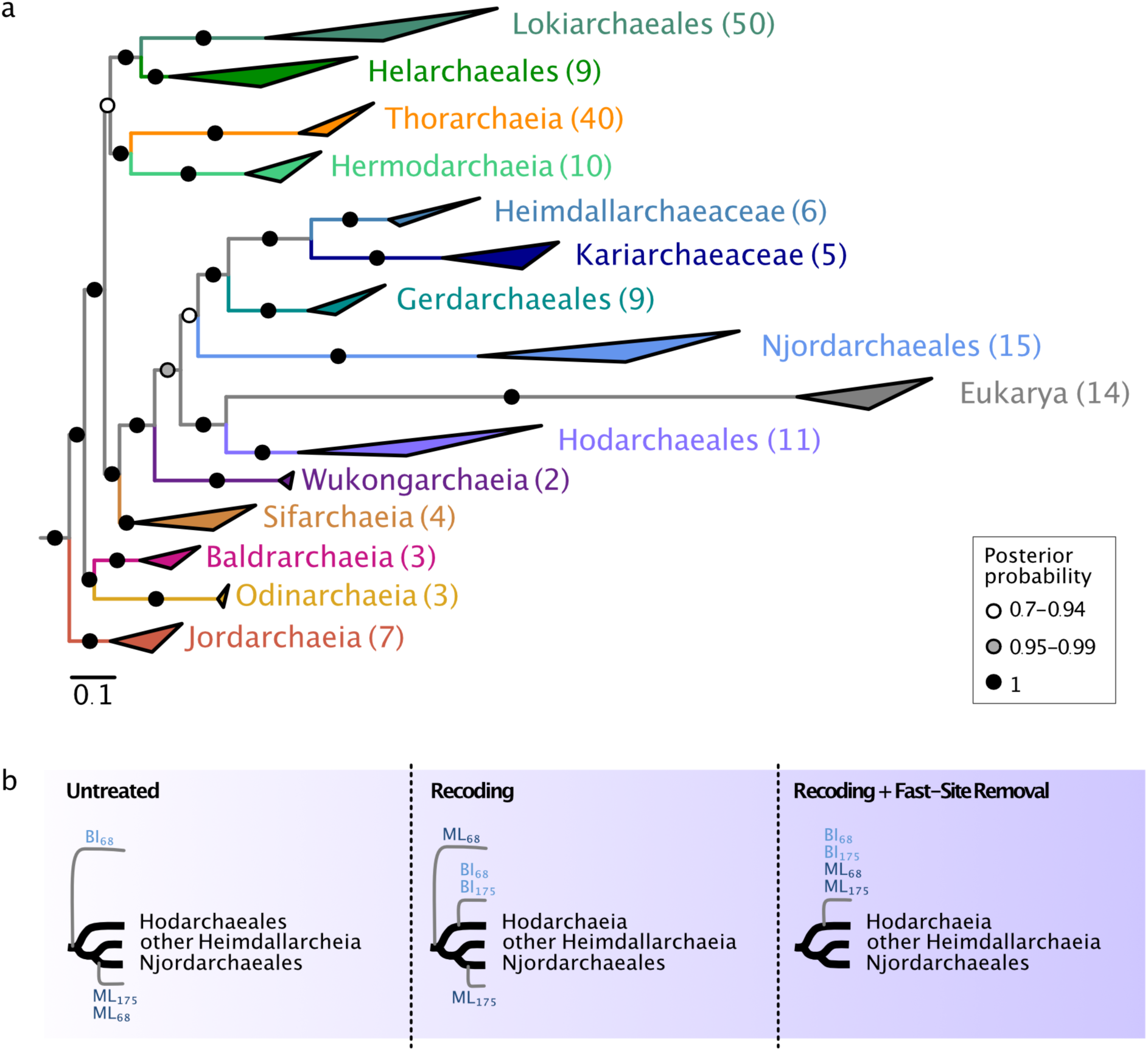
Phylogenomic analyses based on 57 concatenated non-ribosomal proteins support the emergence of eukaryotes as sister to Hodarchaeales. a. Bayesian inference (BI) based on 278 archaeal taxa, using Euryarchaeota and TACK archaea as outgroup (not shown) (NM57-nDK_sr4 alignment, 15,733 amino acid positions). The concatenation was SR4-recoded and analyzed using the CAT+GTR model (4 chains, ∼25,000 generations). b. Schematic representation of the shift in the position of eukaryotes (grey branches) in ML and BI analyses of this dataset under different treatments. Untreated: unprocessed dataset; Recoding: SR4-recoded dataset; Recoding+Fast-Site Removal: Fast-site removal combined with SR4-recoding (the topology most often recovered after removing 10% to 50% fastest-evolving sites, in steps of 10%, is shown). 175 and 68 refer to phylogenomic datasets containing 175 and 68 Asgard archaea, respectively. Note that BI was not performed for the 175 untreated dataset due to computational limitations (for detailed overview of phylogenomic analyses, see Supplementary Table 3). Scale bar denotes the average expected number of substitutions per site.

In summary, resolving the position of eukaryotes relative to Asgard archaea is anything but trivial (see Supplementary Discussion). In our efforts to extract the true phylogenetic signal, we provide confident support for eukaryotes forming a well-nested clade within the Asgard archaea phylum, consistent with the 2D tree of life scenario. More specifically, we observe that eukaryotes affiliate with the Heimdallarchaeia in analyses in which we systematically reduce phylogenetic artefacts, predominantly converging on a position of eukaryotes as sister to Hodarchaeales, which is also in line with the observed ESP content and genome evolution dynamics (see below).

### Informational ESPs in Hodarchaeales

We found that most of the ESPs previously identified in a limited sampling of Asgard archaea^9, 10^ are widespread across all phyla included in the present study (Figure 3, Supplementary Table 3). Notably, we observed some exceptions in support of the phylogenetic affiliation between Hodarchaeales and eukaryotes, particularly among ESPs involved in information processing: (1) the ε DNA polymerase subunit is only found in Hodarchaeales; (2) ribosomal protein L28e (Rpl28e/Mak16) homologs are unique to Njord- and Hodarchaeales members; (3) many archaea that lack genes coding for the synthesis of diphthamide, a modified histidine residue which is uniquely present in archaeal and eukaryotic elongation factor 2 (EF-2), instead encode a second EF-2 paralog that misses key-residues required for diphthamide modification^31^. Interestingly, we found that among all Asgard archaea, only MAGs of all sampled Hodarchaeales members encode *dph* genes in addition to a single gene encoding canonical EF-2, which branches at the base of their eukaryotic counterparts in phylogenetic analyses (Supplementary Figure 11; Supplementary Information); (4) While RPL22e and RNA polymerase subunit RPB8 are found in several Asgard archaeal phyla, the only Heimdallarchaeia genomes encoding these genes are members of the Hodarchaeales. Finally, (5) we identified N-terminal histone tails characteristic of eukaryotic histones in all three Hodarchaeales MAGs, as well as in three Njordarchaeales genomes (see Supplementary Information). Altogether, the identification of these key-informational ESPs, in agreement with phylogenomic analyses described above, supports that Hodarchaeales represent the closest archaeal relatives of eukaryotes.

**Figure 3.**
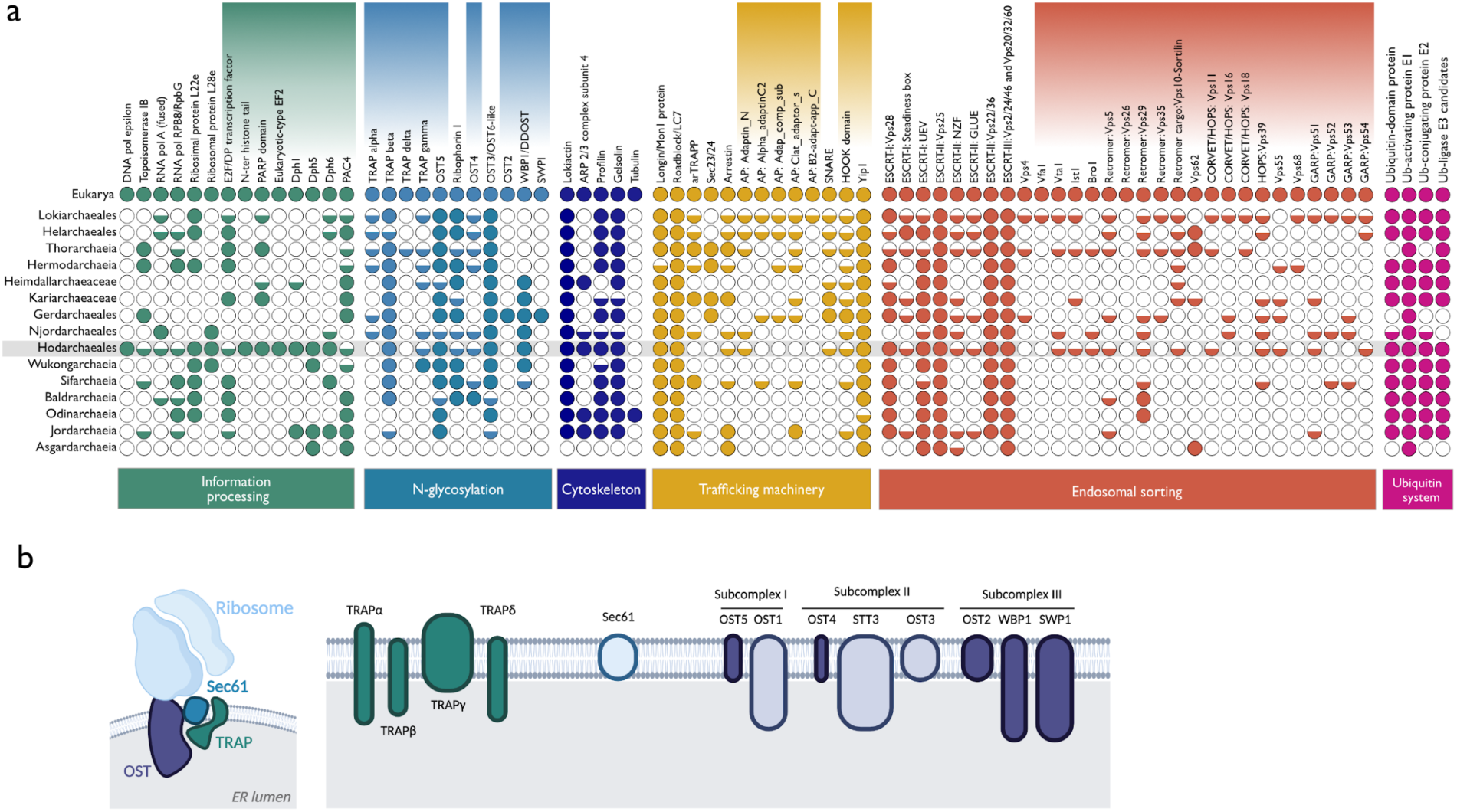
Eukaryotic signature proteins in Asgard archaea. a. Distribution of ESP homologs in Asgard archaea grouped by function. Shaded rectangles above protein names indicate ESPs newly identified as part of this study. Predicted homologs are depicted by colored circles: fully filled circles indicate that we detected homologs in at least half of the representative genomes of the clade; half-filled circles indicate that we detected homologs in fewer than half of the representative genomes of the clade. Hodarchaeales homologs are highlighted with a grey background. Accession numbers are available in **Supplementary Table 3**. b. Asgard archaea encode homologs of eukaryotic protein complexes involved in N-glycosylation. The Sec61, the OST and TRAP complexes are depicted according to their eukaryotic composition and localization. On the right-hand side of the panel, dark-colored subunits represent eukaryotic proteins which have prokaryotic homologs in Asgard archaea newly identified as part of this work; Light-colored subunit homologs have been described previously^10^. Figure generated with Biorender.com.

### Expanded eukaryotic-like protein translocation repertoire

In our search for putative new ESPs in the expanded Asgard archaeal genomic diversity, we uncovered several additional homologs of proteins associated with the eukaryotic translocon, a protein complex primarily responsible for the post-translational modification of proteins, and subsequent insertion into, or transport across the membrane of the endoplasmic reticulum (ER)^32^. The eukaryotic translocon is comprised of the core Sec61 protein-conducting channel, and several accessory components, including the oligosaccharyltransferase (OST) and translocon-associated protein (TRAP) complexes (Figure 3b), both of which are involved in the biogenesis of N-glycosylated proteins^33^. The eukaryotic TRAP complex is composed of two to four subunits in eukaryotes. Using distant-homology detection methods, we identified homologs from three of these subunits to be broadly distributed across Asgard archaeal genomes, while the fourth one was detected only in a few thorarchaeial MAGs (Figure 3b). The eukaryotic OST complex generally comprises 6-8 subunits organized into three subcomplexes that are collectively embedded in the ER membrane^34^ (Figure 3b). Apart from STT3/AglB (OST subcomplex-II), which represents the catalytic subunit and is universally found across all three domains of life, other OST subcomplexes generally do not possess prokaryotic homologs beyond the Ost1/Ribophorin I (OST subcomplex-I) and Ost3/Tusc3 (OST subcomplex-II) subunits previously reported in Asgard archaea^10^. Here, we report the identification of Asgard archaeal homologs of all five additional subunits, Ost2/Dad1, Ost4, Ost5/TMEM258, SWP1/Ribophorin II and WBP1/Ost48. While we identified homologs of Ost4 and Ost5 (OST subcomplex-I) in most Asgard archaeal classes, the distribution of Ost2, WBP1, and Swp1, the first subcomplex-III subunits described in prokaryotes to date, was restricted to Heimdallarchaeia, including Njordarchaeales for WBP1, further supporting their monophyly. Our findings indicate that Asgard archaea and, by inference, LAECA, potentially encode relatively complex machineries for the N-linked glycosylation and translocation of proteins (Figure 3b).

### Vesicular biogenesis and trafficking proteins

Intracellular vesicular transport represents a key process that emerged during eukaryogenesis. Previous studies have reported that Asgard archaeal genomes encode homologs of eukaryotic proteins comprising various intracellular vesicular trafficking and secretion machineries, including the ESCRT (endosomal sorting complexes required for transport), TRAPP (transport protein particle) and COPII (coat protein complex II) vesicle coatomer protein complexes^9, 10^. Furthermore, as much as 2% of the genes of Asgard archaeal genomes were found to encode small GTPase homologs – a broad family of eukaryotic proteins, encompassing the Ras, Rab, Arf, Rho and Ran subfamilies, that are broadly implicated in budding, transport, docking and fusion of vesicles in eukaryotic cells^9, 10^. Here, we report the identification of Asgard archaeal homologs of subunits of additional vesicular trafficking complexes (Figure 3, Ext. Data Fig. 2, Supplementary Table 3). Noticeably, we found putative homologs of all four subunits composing eukaryotic adaptor proteins (AP) and coatomer protein (COPI) complexes, which, in eukaryotic cells, are involved in the formation of clathrin-coated pits and vesicles responsible for packaging and sorting cargo for transport through the secretory and endocytic pathways^35^. Those complexes are composed of two large subunits, belonging to the β- and γ-families, a medium μ-subunit, and a small σ-subunit. We found homologs of all functional domains composing those subunits, albeit sparsely distributed (Ext. Data Fig. 2, Supplementary Information). Additionally, we found homologs of several protein complexes involved in eukaryotic endosomal sorting such as the retromer, the HOPS/CORVET and the GARP complexes (Figure 3, red shading). Retromer is a coat-like complex associated with endosome-to-Golgi retrograde traffic^36^ and we detected four of its five subunits in Asgard MAGs. One of these subunits is Vps5-BAR, which in Thorarchaeia is often fused to Vps28, a subunit of the ESCRT-I subcomplex, suggesting a functional link between BAR domain proteins and the thorarchaeial ESCRT complex. The GARP (Golgi-associated retrograde protein) complex is a multisubunit tethering complex located at the trans-Golgi network in eukaryotic cells, where it also functions to tether retrograde transport vesicles derived from endosomes^37, 38^, similarly to the retromer. GARP comprises four subunits, three of which we could detect in Asgard archaeal genomes, with a sparse and punctuated distribution. Functioning in opposite direction from the retromer and GARP complexes are the CORVET (Class C core vacuole/endosome tethering) and HOPS (Homotypic fusion and protein sorting) complexes^39^. Endosomal fusion and autophagy in eukaryotic cells depend on them and they share four core subunits^40^, three of which can be found in Asgard archaea, in addition to one of the HOPS specific subunits^41^.

Finally, while numerous components of the ESCRT-I, II and III systems have been previously detected in Asgard archaea^9, 10, 42^, we report here the identification of Asgard homologs for the ESCRT-III regulators Vfa1, Vta1, Ist1, and Bro1.

### Ancestral Asgard archaea genome reconstruction

The analysis of Asgard archaeal genome data obtained through metagenomics, combined with the insights derived from cytological observations of the first two cultured Asgard archaea ‘*Candidatus* Prometheoarchaeum syntrophicum’^43^ and ‘*Candidatus* Lokiarchaeum ossiferum’^44^, have generated new hypotheses about the nature of the archaeal ancestor of eukaryotes^43, 45–48^. However, these theories are mostly based on a limited number of features displayed by a single, or a few Asgard archaeal lineages. While informative, features of present-day Asgard archaea do not necessarily resemble those of LAECA, as these are potentially separated by over 2 Gya of evolution^49^. Furthermore, Asgard archaeal phyla display a highly variable genome content with respect to ESPs and predicted metabolic features^43, 46, 48, 50, 51^, suggesting a complex evolutionary history of those traits. In light of these considerations, we inferred ancestral features of LAECA by using an ML evolutionary framework. We employed a recently developed probabilistic gene-tree species-tree reconciliation approach^52, 53^ in combination with the extended taxonomic sampling of Asgard archaeal genomes to reconstruct the evolutionary history of homologous gene families and ancestral gene content across the Asgard archaeal species tree. For this, we inferred ML phylogenetic trees of all 17,200 protein families encoded across 181 archaeal genomes, including representatives from Asgard and TACK archaea, and Euryarchaeota clades. Importantly, as missing genes and potential contaminations in MAGs will be regarded as recent gene loss and gain events in our ancestral reconstruction analyses, the use of incomplete MAGs with low contamination levels is unlikely to have a major impact on the inferred gene content of the deep archaeal ancestors that were reconstructed in the present study (also see Supplementary Information).

We first compared the distributions of estimated ancestral proteome sizes, and numbers of inferred gene duplications, losses and gains (i.e., horizontal gene transfers and originations) in all archaeal ancestral nodes (Supplementary Figure 12). Intriguingly, we observed that Heimdall- (and particularly the ancestor of Hodarchaeales) and Lokiarchaeia ancestors display significantly higher gene duplication rates compared to TACK and Euryarchaeota ancestors (Figure 4a). In addition, we found that most Asgard archaeal ancestors displayed gene loss rates comparable to other archaea, with the exception of Thorarchaeia, Lokiarchaeales and Jordarchaeia, which showed significantly lower loss rates. In agreement with the observed evolutionary genome dynamics, we found that predicted proteome sizes of most Asgard archaea ancestors are significantly larger than other archaeal ancestors (P<0.001), with Lokiarchaeia ancestors displaying the largest estimated proteome size (Supplementary Figure 13). Similarly, the Hodarchaeales ancestor had an estimated proteome size of 4,053 proteins, versus 3,134 for the last Asgard archaea common ancestor (LAsCA), reflecting the high duplication and low loss rates in that clade. The streamlined genome content of the Odinarchaeia ancestor represents an exception to the general trend of genome expansion across Asgard archaea, and possibly reflects an adaptation to high temperatures^54^.

**Figure 4.**
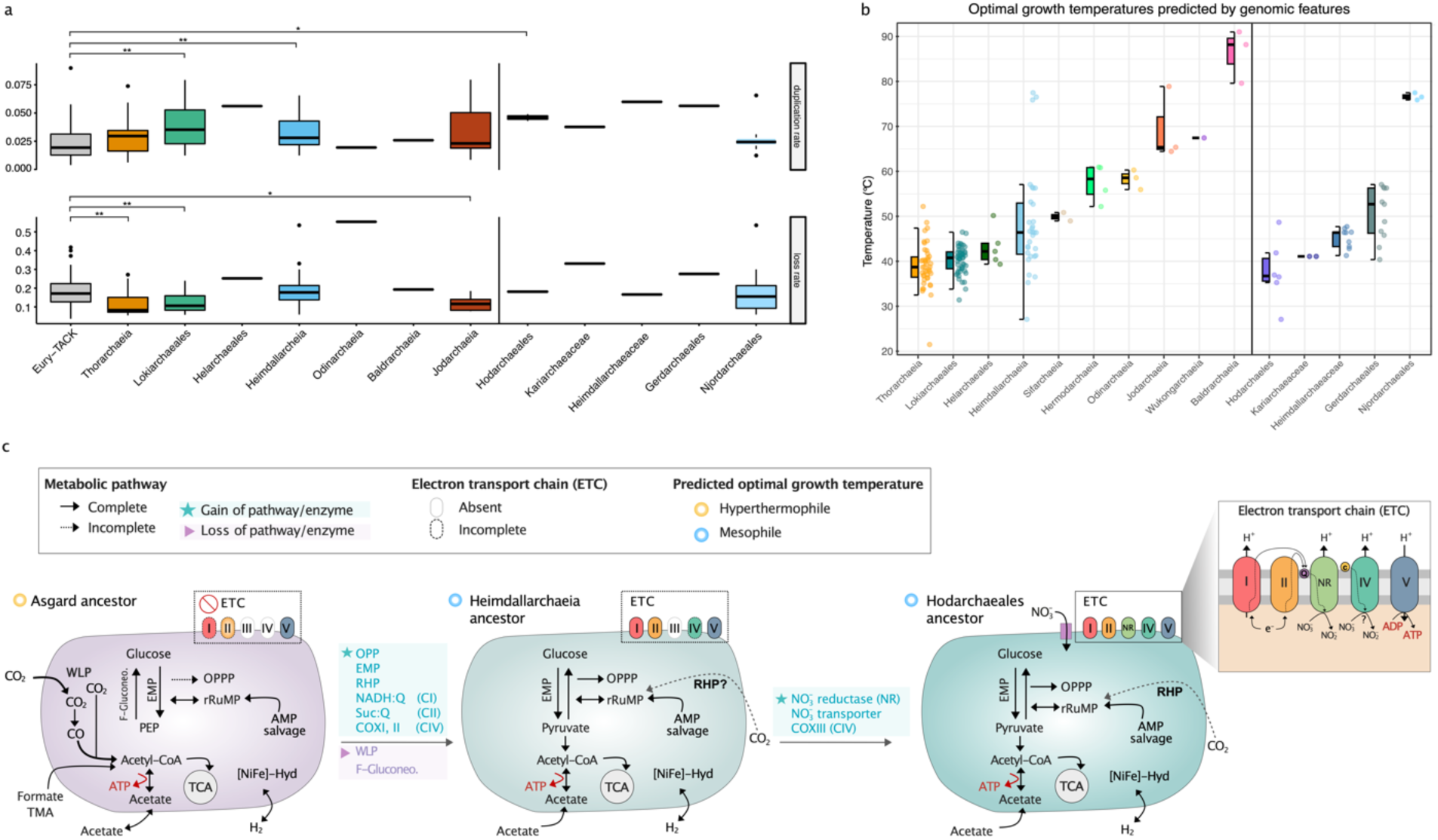
Genome dynamics, Optimal Growth Temperature predictions, and metabolic reconstruction of Asgard ancestors. a. Duplication (upper panel) and loss rates (lower panel) inferred for Asgard archaeal ancestors, normalized by proteome size and plotted by phylum. P-values for each Wilcoxon test against the median values of internal nodes belonging to TACK and Euryarchaeota are shown above each category, where *: p-value <= 0.05, **: p-value <=0.01, ***: p-value <=0.001. b. Optimal Growth Temperature (OGT) predictions, in degrees Celsius. OGT were predicted for the genomes presented here based on genomic and proteomic features^56^ (Supplementary Table 5). Since nucleotide fractions of the ribosomal RNAs are used in this method, only those genomes with predicted rRNA genes could be analyzed. The right-hand panel depicts OGT within Heimdallarchaeia. Note that Njord- and Gerdarchaeales are predicted to be thermophiles (most genomes encode a reverse gyrase). In contrast, Hodarchaeales display the lowest OGT among Heimdallarchaeia. c. Based on the presence/absence of the thermophily-diagnostic enzyme reverse gyrase and the following metabolic signatures in each of the ancestors, we predict that the last Asgard common ancestor probably transitioned from a hyperthermophilic fermentative lifestyle to a mesophilic mixotroph lifestyle. The LAsCA likely encoded gluconeogenic pathways *via* the reverse EMP gluconeogenic pathway and *via* FBP aldolase/phosphatase (FBP A/P). The major energy-conserving step in the early Asgard ancestors could have been the ATP synthesis by fermentation of small organic molecules (i.e., acetate, formate, formaldehyde). The reverse ribulose monophosphate pathway (rRuMP) was a key pathway in the LAsCA for the generation of reducing power. The Wood-Ljungdahl pathway (WLP) appeared only to be present in the LAsCA and was lost in the more recent ancestors (of Heimdallarchaeia and Hodarchaeales) indicated here. The tricarboxylic acid (TCA) cycle is predicted to be complete in all five investigated Asgard ancestors. The inferred presence of the electron transport chain (ETC) components is shown for selected ancestors of major Asgard archaea groups, with the Hodarchaeales common ancestor encoding the most complete set of ETC subunits, and likely using nitrate as a terminal electron acceptor. Therefore, membrane-associated ATP biosynthesis coupled to the oxidation of NADH and succinate and reduction of nitrate to nitrite within the respiratory chain could have been present in the LAECA. Abbreviations: Q: quinone; c: cupredoxin; FBP A/P: Fructose 1,6-bisphosphate aldolase/phosphatase; EMP: Embden-Meyerhof-Parnas; OPPP: Oxidative pentose phosphate pathway; rRuMP: Reversed ribulose monophosphate pathway; RHP: reductive hexulose-phosphate; RuBisCO: Ribulose-1,5-bisphosphate carboxylase/oxygenase; PRK: phosphoribulokinase; AMP: adenosine monophosphate salvage pathway. Details of copy numbers of key enzymes involved in central carbon metabolism are found in Supplementary Table 4.

### Ancestral features of LAECA

Using the approach described above, we also reconstructed the ancestral metabolic and physiological properties across the Asgard archaeal species tree, including the proposed closest archaeal relatives of eukaryotes, the Hodarchaeales. We infer that the LAsCA was a chemolithotroph that required the synthesis of organic building blocks via the Wood-Ljungdahl pathway (WLP) (Figure 4b and Supplementary information), for which we inferred the presence of key enzymes, including carbon monoxide dehydrogenase/acetyl-CoA synthase (CODH/ACS) and the formylmethanofuran dehydrogenase (FmdABCDE). In addition, our analyses revealed that the last common ancestors of individual Asgard archaeal phyla either had the genetic potential to switch between autotrophy and heterotrophy (Loki-, Thor-, Jord- and Baldrarchaeia) or a predominantly heterotrophic fermentative (Odin-and Heimdallarchaeia) lifestyle (Figure 4b, Supplementary Information). Specifically, we observed that the WLP was lost prior to the split between Njordarchaeales and the other Heimdallarchaeia (and therefore prior to the emergence of LAECA), indicating that LAECA was a heterotrophic fermenter (Supplementary Table 4).

Furthermore, we infer that the central carbon metabolism of Heimdallarchaeia (including Hodarchaeales) included the Embden-Meyerhof-Parnas (EMP) pathway and a partial oxidative pentose phosphate (OPP) pathway - both considered core modules of present-day eukaryotic central carbon metabolism. While the enzymes of these pathways in Asgard archaea do not share a common evolutionary origin with those of eukaryotes, this indicates that LAECA had a similar central carbon metabolism compared to modern eukaryotes (Supplementary Figure 14-15).

In addition, our analyses support the idea that the last common ancestor of Heimdallarchaeia contained several components of the electron transport chain (ETC)^48^. We inferred that the last common ancestor of Hodarchaeales likely contained CI, CII, CIV and a nitrate reductase complex (NarGHIJ), indicating that nitrate might have been used as a terminal electron acceptor to perform anaerobic respiration. As such, the last Hodarchaeales common ancestor likely generated ATP using an electron transport chain where electrons from NADH and succinate were transferred through a series of membrane-associated complexes with quinones and cupredoxins as electron carriers to ultimately reduce nitrate^55^.

As indicated above, a significant fraction of the currently sampled Asgard archaea diversity originates from geothermal or hydrothermal environments. Indeed, using an algorithm based on genome-derived features^56^, we confirmed that (most) Njordarchaeales, Baldr- and Jordarchaeia are hyperthermophiles, Odinarchaeia are thermophiles, and Loki- and Thorarchaeia are mesophiles (Figure 4c, Supplementary Table 5). While Heimdallarchaeia seem to contain both meso- and thermophiles, we infer a mesophilic physiology for Hodarchaeales, obtaining the lowest predicted optimal growth temperatures among all Asgard archaea (median=36.7 °C). Asgard archaeal hyperthermophiles contain reverse gyrase, a topoisomerase that is typically encoded by hyperthermophilic prokaryotes^57^. We infer that a reverse gyrase was possibly present in LAsCA and that it was subsequently lost in all heimdallarchaeial orders except for Njordarchaeales. This observation would be compatible with a scenario in which Asgard archaea have a hyperthermophilic ancestry, but in which eukaryotes evolved from an Asgard archaea lineage that had adapted to mesophilic growth temperatures.

## Discussion

Beyond genomic exploration, several studies have started to unveil important physiological, cytological and ecological aspects of Asgard archaea^43, 58–60^. Yet, while such insights are certainly relevant, the cellular and physiological characteristics of present-day Asgard archaea will almost certainly not resemble those of LAECA. Therefore, inferences about the identity and nature of LAECA and the process of eukaryogenesis should be made within an evolutionary context. We used an evolutionary framework to analyze an expanded Asgard archaeal genomic diversity, comprising 11 clades of high taxonomic rank. Using comprehensive phylogenomic analyses involving the evaluation of distinct marker protein datasets and systematic assessment of suspected phylogenetic artefacts and state-of-the-art models of evolution, we identified Hodarchaeales, a class-level clade within the Heimdallarchaeia, as the closest relatives of eukaryotes. Evidently, phylogenomic analyses aiming to pinpoint the phylogenetic position of eukaryotes in the tree of life are extremely challenging, and our results stress the importance of testing for possible sources of bias affecting phylogenomic reconstructions, as was recently reviewed^61^. The implementation of a probabilistic gene tree/species tree reconciliation approach allowed us to infer the evolutionary dynamics and ancestral content across the archaeal species tree providing several new insights into the Asgard archaeal roots of eukaryotes. Altogether, our results reveal a picture in which the Asgard archaeal ancestor of eukaryotes had, compared to other archaea, a relatively large genome, resulting mainly from more numerous gene duplication and fewer gene loss events. It is tempting to speculate that the elevated gene duplication rates observed in our analyses represent an ancestral feature of LAECA, and that it remained the predominant modus of genome evolution during the early stages of eukaryogenesis. We also inferred that the duplicated gene content of LAECA included several protein families involved in cytoskeletal and membrane-trafficking functions, including among others actin homologs, ESCRT complex subunits and small GTPase homologs. Our findings complement those of another study^62^ reporting that eukaryotic proteins with an Asgard archaeal provenance, as opposed to those inherited from the mitochondrial symbiont, duplicated the most during eukaryogenesis, particularly proteins of cytoskeletal and membrane-trafficking families.

Beyond genome dynamics, our analyses of inferred ancestral genome content across the Asgard archaeal species tree indicates that, while Asgard archaea likely had a thermophilic ancestry, the lineage from which eukaryotes evolved was adapted to mesophilic conditions, which is compatible with a generally assumed mesophilic ancestry of eukaryotes. Furthermore, we infer that LAECA had the genetic potential to support a heterotrophic lifestyle, and may have been able to conserve energy via nitrate respiration. In addition, based on taxonomic distribution and evolutionary history of ESPs we show that complex pathways involved in protein targeting and membrane trafficking, and in genome maintenance and expression in eukaryotes were inherited from their Asgard archaeal ancestor. Of note, we identified additional Asgard archaeal homologs of eukaryotic vesicular trafficking complex components. Of these, some Asgard archaeal proteins display sequence similarity to proteins which, in eukaryotes, are part of the clathrin adaptor protein complexes and of the COPI complex. These complexes are particularly interesting since they are involved in the biogenesis of vesicles responsible for sorting cargo and subsequent transport through the secretory and endocytic pathways^35^. Altogether, these results further suggest a potential for membrane deformation, and possibly trafficking, in Asgard archaea. The ability to deform membranes was recently shown in two papers reporting the first cultivated Lokiarchaeia lineages, ‘*Ca.* P. syntrophicum strain MK-D1’^43^ and ‘*Ca.* Lokiarchaeum ossiferum’^44^, whose cells both displayed unique morphological complexity including long and often branching protrusions facilitated by a dynamic actin cytoskeleton. Thus far no^43^, or only limited^44^ visible endomembrane structures were observed in these first cultured representatives of Asgard archaea. However, it is important to restate here that, being separated by some 2 Gya of evolution, the cellular features of present-day Asgard archaeal lineages do not necessarily resemble those of LAECA. Furthermore, given the disparity of the distribution patterns of membrane trafficking homologs in Asgard archaea, it will be crucial to isolate representatives of phyla other than Lokiarchaeia and study their cell biological features and potential for endomembrane biogenesis. Of particular interest would be members of the Heimdallarchaeia, and specifically Hodarchaeales, as the currently identified closest relatives of eukaryotes, as well as Thorarchaeia lineages, which seem to generally contain a particularly rich suite of homologs of eukaryotic membrane trafficking proteins.

By phylogenetically placing eukaryotes as a firmly-nested clade within the presently identified Asgard archaeal diversity, and by inferring ancestral genomic content across the Asgard archaea, our work provides insights into the identity and nature of the Asgard archaeal ancestor of eukaryotes, guiding future studies aiming to uncover new pieces of the elusive eukaryogenesis puzzle.

## Methods

### Sample collection, sequencing, assembly and binning

We sampled aquatic sediments from eleven geographically distant sites: Guaymas Basin (Mexico), Lau Basin (Eastern Lau Spreading Center and Valu Fa Ridge, south-west Pacific Ocean), Hydrate Ridge (offshore of Oregon, USA), Aarhus Bay (Denmark), Radiata Pool (New Zealand), Taketomi Island Vent (Japan), the White Oak River estuary (USA), and Tibet Plateau and Tengchong (China) (Supplementary Table 1).

*a. Jordarchaeote JZB50, QC4B49, QZMA23B3, QZMA2B5, QZMA3B5*

Hot spring sediment samples were collected from Tibet Plateau and Yunnan Province (China) in 2016. The microbial community compositions have been described and reported previously^63, 64^. Samples were collected from the hot spring pools using a sterile iron spoon into 50 ml sterile plastic tubes, then transported to the lab on dry ice, and stored at -80°C for DNA extraction. The genomic DNA of the sediment samples was extracted using FastDNA Spin Kit for Soil (MP Biomedicals, Irvine, CA) according to the manufacturer’s instructions. The obtained genomic DNA was purified for library construction, and sequenced on an Illumina HiSeq2500 platform (2 × 150 bp). The raw reads were filtered to remove Illumina adapters, PhiX and other Illumina trace contaminants with BBTools v38.79, and low-quality bases and reads using Sickle (v1.33; https://github.com/najoshi/sickle). The filtered reads were assembled using metaSPAdes (v3.10.1) with a kmer set of “21, 33, 55, 77, 99, 127”. The filtered reads were mapped to the corresponding assembled scaffolds using bowtie2 v2.3.5.1^65^. The coverage of a given scaffold was calculated using the command of “jgi_summarize_bam_contig_depths” in MetaBAT v2.12.1^66^. For each sample, scaffolds with a minimum length of 2.5 kbp were binned into genome bins using MetaBAT v2.12.1, with both tetranucleotide frequencies (TNF) and scaffold coverage information considered. The clustering of scaffolds from the bins and the unbinned scaffolds was visualized using ESOM with a minimum window length of 2.5 kbp and max window length of 5 kbp, as previously described^67^. Misplaced scaffolds were removed from bins and unbinned scaffolds whose segments were placed within the bin areas of ESOMs were added to the corresponding bins. Scaffolds with a minimum length of 1 kbp were uploaded to ggKbase (http://ggkbase.berkeley.edu/). The ESOM-curated bins were further evaluated based on consistency of GC content, coverage and taxonomic information, and scaffolds identified with abnormal information were removed. The ggKbase genome bins were curated individually to fix local assembly errors using ra2.py ^68^.

*b. Njordarchaeote A173, A3132, M288 and Thorarchaeote A361, A381, A399*

Hydrothermal vent deposits were collected from the ABE (ABE 1, 176° 15.48’W, 21° 26.68’S, 2142 m; ABE 3, 176° 15.59’W, 21° 26.95’S, 2131 m) and Mariner (176° 36.07’W, 22° 10.81’S, 1914 m) vent fields along the Eastern Lau Spreading Center in April/May of 2015 during the RR1507 Expedition on the RV Roger Revelle. Sample collection and processing were done as previously described^69^. DNA was extracted from homogenized rock slurries using the DNeasy PowerSoil kit (Qiagen) as per the manufacturer’s instructions. Samples were prepared for sequencing on the Illumina HiSeq 3000 using Nextera DNA Library Prep kits (Illumina), and metagenomes (2×150 bp) were sequenced at the Oregon State University Center for Genome Research and Computing. Trimmomatic^70^ v.0.36 was used to trim low-quality regions and adapter sequences from raw reads (parameters: ILLUMINACLIP:TruSeq3-PE-2.fa:2:30:10, LEADING:20, SLIDINGWINDOW:4:20, MINLEN:50). Clean paired reads were then interleaved using the khmer software package^71^. Interleaved and unpaired reads were assembled with MEGAHIT v.1.1.1-2-g02102e1 (--k-min 31, --k-max 151, --k-step 20, --min-contig-len 1000)^72, 73^. Trimmed reads were mapped back to the contigs to determine read coverage using Bowtie 2 v.2.2.9 ^65, 74^ and SAMtools v.1.3.1^75^. Binning was performed with MetaBAT v.0.32.4^66^ using tetranucleotide frequency and read coverage. Bin completion and contamination were estimated with CheckM v.1.0.7^76^.

*c. Lokiarchaeote ABR01, ABR02, ABR03, ABR04, ABR05, ABR06, ABR08, ABR11, ABR13, ABR15, Thorarchaeote ABR09, ABR10 and Heimdallarchaeote ABR14, ABR16* MAGs were obtained as previously described^31^.

*d. Archaeon WORA1, WORB2, Heimdallarchaeote WORE3, Lokiarchaeote WORB4, WORC5 and Thorarchaeote WORH6*

Sampling, DNA extraction, sequencing library preparation and sequencing methods were previously described^77^. Published assemblies and raw reads for the samples WOR-1-36_30 (SAMN06268458; Gp0056175), WOR-1-52-54 (SAMN06268416; Gp0059784), WOR-3-24_28 (SAMN06268417; Gp0059785) were downloaded from JGI. Short reads were trimmed using Trimmomatic^70^ v0.33 (PE ILLUMINACLIP:2:30:10 SLIDINGWINDOW:4:15 MILEN:100). Contigs shorter than 1000 bp were excluded from the assembly using SeqTK v1.0r75 (https://github.com/lh3/seqtk). Each assembly was binned using CONCOCT v0.4.1^78^ and coverage information from the three datasets, and Asgard bins were subsequently identified based on phylogenies of concatenated ribosomal proteins^10^. Identified Asgard MAGs were used together with publicly available Asgard genomes to recruit trimmed-reads originated from Asgard genomes using CLARK v1.2.3 with the -m 0 option^79^. For each dataset, recruited Asgard reads were independently assembled using SPAdes^80^ and IDBA-UD^81^ and further binned using CONCOCT, using a minimum contig length of 1000 bp. Bins with higher completeness and lower contamination values as predicted by miComplete v1.00^82^ were selected and manually curated using mmgenome v0.7.1^83, 84^ using the coverage information, paired-reads linkage, composition and marker genes information. The samples and assembly method used for each final MAG were: Archaeon WORA1 (WOR-1-52-54; spades), Archaeon WORB2 (WOR-1-52-54; IDBA-UD), Heimdallarchaeote WORE3 (WOR-3-24_28; spades), Lokiarchaeote WORB4 and WORC5 (WOR-1-36_30; IDBA-UD), and Thorarchaeote WORH6 (WOR-1-36_30; spades).

*e. Jordarchaeote RPD1, RPF2 and Odinarchaeote RPA3*

Information about the location of the hot spring sediments from Radiata Pool, sampling and DNA extraction procedures was previously reported^10^. Short paired-end Illumina reads were generated and preprocessed using Scythe (https://github.com/vsbuffalo/scythe) and Sickle (https://github.com/najoshi/sickle) to remove adapters and low-quality reads. Reads were subsequently assembled of IDBA-UD 1.1.3 (--maxk 124). The Jordarchaeote RPF2 MAG was generated by binning contigs according to their tetranucleotide frequencies using esomWrapper.pl (https://github.com/tetramerFreqs/Binning) with a minimum contig length 5000 bp and a window size of 10 Kbp. ESOM maps were manually delineated using the Databionic ESOM viewer (http://databionic-esom.sourceforge.net/). Jordarchaeote RPD1 and Odinarchaeote RPA3 were binned following the methodology described in above (section d), but re-assembling the recruited reads only with IDBA-UD (--maxk 124)^81^.

*f. Jordarchaeote GBS01, GBS02, GBS03, GBS04, GBS05, GBS06, GBS07, Heimdallarchaeote GBS08, GBS09, GBS10, GBS11, Lokiarchaeote GBS14, Njordarchaeote GBS15, GBS16, GBS17, GBS18, GBS19, GBS20, GBS21, GBS22, GBS23, GBS24, GBS25, GBS26, TNS08 and Thorarchaeote GBS28, GBS29, GBS33, GBS34* MAGs were obtained as described in ^85^.

*g. Heimdallarchaeote B3_JM_08* MAG was obtained as described in ^86^.

*h. Thorarchaeote OWC_bin2, OWC_bin3 and OWC_bin4* MAGs were obtained as described in ^31^.

*i. Heimdallarchaeote GBS11*

Samples were made available by the Gulf Coast Repository (GCR) and were collected on the Ocean drilling Program (ODP) Leg 204 at site 1244 (44°35.17N, 125°7.19W) on July 14th, 2002 (hole C and core 2). The ODP site is found at a water depth of 890 m on the eastern side of the South Hydrate Ridge on the Cascadia Margin. This site has been well characterized physically and geochemically^87^. Furthermore, the microbial community structure has been surveyed using 16S rRNA gene sequencing^88, 89^. Two sediment samples, designated DCO-2-5 (sample ID 1489929) and DCO-2-7 (sample ID 1489924), were collected at a sediment depth of 12.40 and 14.96 m below the seafloor, respectively, and stored at -80°C at GCR. A total amount of 10 g of each of the two sediment samples was used to extract DNA using the MoBio DNA PowerSoil Total kit. A total amount of 100 ng DNA was used to prepare sequencing libraries that were 150 bp paired-end sequenced at the Marine Biological Laboratory (Woods Hole, MA, USA) on an Illumina MiSeq sequencer. Adaptors and DNA spike-ins were removed from the forward and reverse reads using cutadapt v1.12^90^. Afterwards, reads were interleaved using interleave_fasta.py (https://github.com/jorvis/biocode/blob/master/fasta/interleave_fasta.py), and further trimmed using Sickle with default settings (Fass JN) (https://github.com/najoshi/sickle). Metagenomic reads from both samples were co-assembled using IDBA-UD using the following parameters: —pre_correction, -mink 75, -maxk 105, --step 10, --seed_kmer 55 ^81^. Metagenomic binning was performed on scaffolds with a length >3,000 bp using ESOM, including a total of 4,939 scaffolds with a length of 30,693,002 bp^67, 81^. CheckM v1.0.5 was employed to evaluate the accuracy of the binning approach by determining the percentage of completeness and contamination^76^.

*j. Heimdallarchaeote GBS09*

MAG was obtained as previously described^91^.

### Exploration of phylogenetic diversity in Asgard assemblies and MAGs

To assess the presence of potential Asgard-related lineages in our assemblies, we reconstructed a phylogeny of ribosomal proteins encoded in a conserved 15-ribosomal protein (RP15) gene cluster^16^. As ingroup, we used all MAGs presented in this study, plus all genomes classified as Asgard archaea in NCBI as of June 25^th^ 2021, plus those classified as “archaeon” corresponding to Hermodarchaeia (GCA_016550385.1, GCA_016550395.1, GCA_016550405.1, GCA_016550415.1, GCA_016550425.1, GCA_016550485.1, GCA_016550495.1, GCA_016550505.1). and all Asgard archaeal MAGs released by Sun et al.^21^. To obtain an adequate outgroup dataset, we downloaded all archaeal genomes from the Genome Taxonomy Database^92^, data revision 89, and selected one genome sequence per species-level cluster as defined in https://data.gtdb.ecogenomic.org/releases/release89/89.0/sp_clusters_r89.tsv. We then selected a set of 216 genomes classified as Bathyarchaeia, Nitrososphaeria and Thermoprotei, and used them as outgroup. Genes were detected and individually aligned and trimmed as previously described^10^. Ribosomal protein sequences were selected if they were encoded in a contig containing at least five of the 15 ribosomal protein genes. ModelFinder^93^ was run as implemented in IQ-TREE v. 2.0-rc2 to identify the best model among all combinations of the LG, WAG, JTT and Q.pfam models, as well as their corresponding mixture models by adding +C20, +C40 and +C60, and the additional mixture models LG4M, LG4X, UL2 and UL3, with rate heterogeneity (none, +R4 and +G4) and frequency parameters (none, +F). A PMSF approximation^94^ of the chosen model (WAG+C60+R4+F) was then used for a final reconstruction using 100 non-parametric bootstrap pseudoreplicates for branch statistical support. The obtained tree revealed a broad genomic diversity of Asgard lineages (Figure 1).

### Environmental distribution of Asgard archaea

16S rRNA gene sequences were predicted with Barrnap v 0.9 (https://github.com/tseemann/barrnap) with the option “--kingdom arc”. Since none of the two Helarchaeales bins contained 16S rRNA gene sequences, helarchaeal 16S rRNA gene sequences identified by Seitz et al.^50^ were used as representatives of this phylum. These sequences were submitted to the IMNGS platform as queries for Paraller similarity searches against all available NCBI sequence read archive (SRA) samples with a 95% similarity threshold and a minimum alignment size of 200 bp^95^. Available metadata for detected SRA samples were then manually assessed to link environmental context descriptions for individual SRA samples to broader environment categories. The sequence abundance output file generated by IMNGS was then analysed using R and the package tidyverse to calculate the number of SRA samples belonging to each environment per phylum^96^.

### Gene prediction

Gene prediction was performed using Prokka^97^ v1.12 (prokka --kingdom Archaea --norrna -- notrna). rRNA genes and tRNA genes were predicted with Barrnap (https://github.com/tseemann/barrnap) and tRNAscan-SE^96, 98^, respectively.

### Optimal growth temperature prediction

Optimal growth temperatures were predicted for the genomes presented here based on genomic and proteomic features^56^ (see Supplementary Information). Since ribosomal RNAs nucleotide composition are used in this method, only genomes with predicted rRNAs were analyzed.

### Identification of homologous protein families

All-versus-all similarity searches of all predicted proteins from the A64 taxon selection (64 Asgard, 76 TACK, 43 Euryarchaeota and 41 DPANN archaea; see Supplementary Table 2) were performed using diamond^99^ blastp (--more-sensitive --evalue 0.0001 --max-target-seqs 0 --outfmt 6). The file generated was used to cluster protein sequences into homologous families using SiLiX^100^ v.1.2.10, followed by Hifix^101^ v1.0.6. The identity and overlap parameters required by Silix were set to 0.2 and 0.7, respectively, after inspecting a wide range of values (--ident [0.15,0.4] and --overlap [0.55-0.9], with increments of 0.05) and selecting the values that maximized the number of clusters containing at least 80% of the taxa.

### Functional annotation of homologous protein families

Protein families, excluding singletons, were aligned using mafft-linsi^102^ v7.402 and converted into HHsearch format (.hhm) profiles using HHblits^103^ v3.0.3. Profile-profile searches were subsequently performed against a database containing profiles from EggNOG 4.5^104^, arCOGs^105^ and PFAM databases^106^ that had been previously converted to the hhm format using HHblits^103^ v3.0.3.

### Automatic functional annotation of individual proteins

Individual proteins were annotated using the HMMscan tool of the HMMer suite against PFAM v32^106^, Interproscan^107^ 5.25-64.0, EggNOG mapper v0.12.7 against the NOG database v4.5^104^, diamond aligner v0.9.9.110^99^ against the nr database, arCOG^105^, and GhostKoala annotation server108. Putative hydrogenases were further classified using HydDB109.

### Detailed analysis of ESPs

In-depth analysis of potential ESPs involved a combination of automatic screens and manual curation. We first manually searched for homologs of previously described ESPs^9, 10, 42^ by using a variety of sequence similarity approaches such as BLAST, HMMer tools, profile-profile searches using HHblits, combined with phylogenetic inferences, and, in some cases, the Phyre2 structure homology search engine^103, 110, 111^. We did not use fixed cutoffs, as the e-value between homologs will vary depending on the protein investigated, hence the need for manual examination of potential homologs and a combination of lines of evidence.

In addition, to identify potential new ESPs, we first used our profile-profile searches against EggNOG and manually investigated Asgard orthologous groups which had a best hit to a eukaryotic-specific EggNOG cluster. We also extracted PFAM domains whose taxonomic distribution is exclusive to eukaryotes as per PFAM v32, and investigated cases where they represented the best domain hit in Asgard archaea sequences identified by HMMscan. Finally, we manually investigated dozens of proteins known to be involved in key eukaryotic functions based on our knowledge and literature searches. In Figure 2, we are only reporting cases based on the strict cutoff that the diagnostic HMM profile had the best score among all profiles detected for a protein. An exception was made for the ESCRT domain Vps28, Steadiness box, UEV, Vps25, NZF, GLUE and Vps22 domains which are usually found in combination with other protein domains and thus do not necessarily represent the best scoring domain in a protein even if they represent true homologs.

### Phylogenetic analyses of concatenated proteins for species tree inference

Two sets of phylogenetic markers were used to infer the species tree. The first one (RP56) is based on a previously published dataset of 56 ribosomal proteins used to place the first assembled Asgard genomes^10^. The second one (NM57, for ‘new markers’) corresponds to 57 proteins extracted from a set of 200 markers previously identified as core archaeal proteins that can be used to robustly infer the tree of archaea^112^. These 57 markers were selected because they were found in at least a third of representatives of each of the 11 Asgard clades, as well as in 10 out of 14 eukaryotes, and were inherited from archaea in eukaryotes.

We initially assembled an RP56 dataset for a phylogenetically diverse set of 222 archaeal and 14 eukaryotic taxa. These included all 11 Asgard archaea MAGs and genomes available at the NCBI as of May 12, 2017, as well as the 53 most diverse novel MAGs from this work (out of 63). We gathered orthologs of these genes from all proteomes by using sequences from the previously published alignment^10, 112^ as queries for BLASTp. For each marker, the best BLAST hit from each proteome was added to the dataset. For the first iteration, each dataset was aligned using mafft-linsi^113^ and ambiguously aligned positions were trimmed using BMGE (-m BLOSUM30)^114^. All 56 trimmed RP alignments were concatenated into an RP56-A64 supermatrix (236 taxa including 64 Asgard archaea, 6332 amino acid positions). Once this taxon set was gathered, we identified homologs of the NM57 gene set as described above, thus generating supermatrix NM57-A64 (236 taxa, 14,847 amino acid positions).

We carried out a large number of phylogenomic analyses on variations of these two RP56-A64 and NM57-A64 datasets with different phylogenetic algorithms. Notably, preparing these datasets must be done with great care and is therefore time-consuming, and subsequent phylogenomic analyses generally require an enormous amount of computational running time. However, the rapid expansion of available Asgard archaeal MAGs, notably by Liu and colleagues as of April 2021^15^, urged us to update and re-run many of the computationally demanding analyses. As some of the work that was based on a more restrained taxon sampling is still deemed valuable, such as some of the Bayesian phylogenomic analyses and ancestral genome content reconstructions, we retained these in the present study.

An updated Asgard archaeal genomic sequence dataset was constructed by including all 230 Asgard archaeal MAGs and genomes available at the NCBI database as of May 12, 2021, as well as 63 novel MAGs described in the present work. All 56 trimmed RP alignments were concatenated into an RP56-A293 supermatrix (465 taxa including 293 Asgard archaea, 7112 amino acid positions), which was used to infer a preliminary phylogeny with FastTree v2^115^ (Supplementary Figure 16). Given the high computational demands of the subsequent analyses, we then used this phylogeny to select a subsample of Asgard archaea representatives. For this, we first removed the most incomplete MAGs encoding fewer than 19 ribosomal proteins (i.e., 1/3 of the markers) in the matrix. We also used the preliminary phylogeny to sub-select among closely related taxa: among taxa that were separated by branch lengths of <0.1, we only kept one representative. This led to a selection of 331 genomes, including 175 Asgard archaea, 41 DPANN, 43 Euryarchaeota, and 72 TACK representatives (RP56-A175 dataset). Out of these 175 Asgard archaea, 41 correspond to MAGs newly reported here. Once this taxon set was gathered, we identified homologs of the NM57 gene set as described above, thus generating supermatrix NM57-A175. All datasets and their composition are summarized in Supplementary Table 2.

To test for potential phylogenetic reconstruction artefacts, our datasets were subjected to several treatments. Supermatrices were recoded into four categories, using the SR4 scheme^27^. The corresponding phylogenies were reconstructed with IQ-TREE (using a user-defined previously described model referred to as ‘C60SR4’, based on the implemented ‘LG+C60’ model and modified to analyze the recoded data^10^) and Phylobayes (under the CAT+GTR model)^10^. We also used the estimated site rate output generated by IQ-TREE (-wsr) to classify sites into 10 categories, from the fastest to the slowest evolving, and we removed them in a stepwise fashion, removing from 10% to 90% of the data. Finally, we combined both approaches by applying SR4 recoding to the alignments obtained after each fast-site removal step. All phylogenetic analyses performed are summarized in Supplementary Table 2. See Supplementary Information for details and discussion.

### Analyses of individual proteins

For individual proteins of interest, we gathered homologs using various approaches, depending on the level of conservation across taxa. In order to detect putative Asgard homologs of eukaryotic proteins, we used a combination of tools including BLASTp^116^ and the HMMer toolkit (http://hmmer.org/) if HMM profiles were available, and queried a local database containing our 240 archaeal representatives (including all Asgard predicted proteomes). We then investigated the Asgard candidates by 1) using them as seed for BLASTp searches against nr; 2) by 3D modelling using Phyre2 and Swissmodel when sequence similarity was low; 3) by annotating them using Interproscan 5.25-64.0^107^, EggNOG mapper v0.12.7^117^, against the NOG database^117^, and GhostKoala annotation server^108^; 4) by annotating the archaeal orthologous cluster they belonged to using profile-profile annotation as described above. Eukaryotic homologs were gathered from the UniRef50 database^118^. Depending on the divergence between homologs, they were aligned using mafft-linsi and trimmed using TrimAl^119^ (--automated1) or BMGE^114^, or, in cases where we investigated a specific functional domain, we used the hmmalign tool from the HMMer package with the --trim flag to only keep and align the region corresponding to this domain. When divergence levels allowed, phylogenetic analyses were performed using IQ-TREE with model testing including the C-series mixture models (-mset option)^120^. Statistical support was evaluated using 1000 ultrafast bootstrap replicates (for IQ-TREE)^119^.

### Ancestral reconstruction

For the ancestral reconstruction analyses, only a subset of 181 taxa were included (64 Asgard, 74 TACK and 43 Euryarchaeota, see Supplementary Table 2 for details). Protein families with more than three members were aligned and trimmed using mafft-linsi v7.402^113^ and trimAl v1.4.rev15 with the --gappyout option^119^. Tree distributions for individual protein families were estimated using IQ-TREE v1.6.5 (-bb 1000 -bnni -m TESTNEW -mset LG -madd LG+C10,LG+C20 -seed 12345 -wbtl -keep-ident)^122^. The species phylogeny together with the gene tree distributions were subsequently used to compute 100 gene-tree species tree reconciliations using ALEobserve v0.4 and ALEml_undated^52, 53^, including the fraction_missing option that accounts for incomplete genomes. The genome copy number was corrected to account for the extinction probability per cluster (github.com/maxemil/ALE/commit/136b78e). The missing fraction of the genome was calculated as 1 minus the completeness values (in fraction) as estimated by CheckM v1.0.5 for each of the 181 taxa^76^. Protein families containing only one protein (singletons) were considered as originations at the corresponding leaf. The ancestral reconstruction of 5 protein families that included more than 2000 proteins raised errors and could not be computed. The minimum threshold of the raw reconciliation frequencies for an event to be considered was set to 0.3 as commonly done^123–126^ and recommended by the authors of ALE (Gergely Szölősi, personal communication).

### Ancestral metabolic inferences

Metabolic reconstruction of the Asgard ancestors was based on the inference, annotation and copy number of genes in ancestral nodes. The presence of a given gene was scored if its copy number in the ancestral nodes was above 0.3. A protein family was scored as “maybe present” if the inferred copy number was between 0.1 and 0.3. The protein annotation of each of the clusters containing the ancestral nodes was manually verified for each of the enzymatic steps involved in the pathways detailed in Supplementary Table 4.

## Data availability

The MAGs reported in this study have been deposited at DDBJ/EMBL/GenBank. BioProject IDs, BioSample IDs and GenBank assembly accession numbers are available in Supplementary Table 1 and will be released upon publication of the manuscript. All raw data underlying phylogenomic analyses (raw and processed alignments and corresponding phylogenetic trees) will be deposited on Figshare (https://figshare.com/account/home#/projects/111912) upon publication of the manuscript.

## Methods references

## Acknowledgements

We thank Stephan Köstlbacher for intellectual input. We thank the Uppsala Multidisciplinary Center for Advanced Computational Science (UPPMAX) at Uppsala University and the Swedish National Infrastructure for Computing (SNIC) at the PDC Center for High-Performance Computing for providing computational resources. We thank the Japan Agency for Marine-Earth Science & Technology (JAMSTEC) for taking sediment samples from the Taketomi shallow submarine hydrothermal system, the crew of the RV Roger Revelle for assisting with the sampling of the ABE and Mariner vent fields along the Eastern Lau Spreading Center during the RR1507 Expedition. The Ngāti Tahu - Ngāti Whaoa Runanga Trust is acknowledged as *mana whenua* of Radiata Pool and associated samples, and we thank them for their assistance in access and sampling of the Ngatamariki geothermal features. Sampling in the Eastern Lau Spreading Center and Guaymas Basin (Gulf of California) was supported by the US-National Science Foundation (NSF-OCE-1235432 to A.-L. R. and NSF-OCE-0647633 to A.T.). A subset of Guaymas sediments were sequenced by the U.S. Department of Energy Joint Genome Institute, a DOE Office of Science User Facility under Contract No. DE-AC02-05CH11231 granted to ND. We thank the captain and crew of RV Aurora for assistance during sampling at Aarhus Bay. Sampling at Aarhus Bay was supported by the VILLUM Experiment project “FISHing for the ancestors of the eukaryotic cell” (grant number 17621 to A.S. and K.U.K.). This work was supported by grants of the European Research Council (ERC Starting and Consolidator grants 310039 and 817834, respectively), the Swedish Research Council (VR grant 2015-04959), the Dutch Research Council (NWO-VICI grant VI.C.192.016), Marie Skłodowska-Curie ITN project SINGEK (H2020-MSCA-ITN-2015-675752) and the Wellcome Trust foundation (Collaborative award 203276/K/16/Z) awarded to T.J.G.E.. L.E. was supported by a Marie Skłodowska-Curie IEF (grant 704263), and by funding from the European Research Council (ERC Starting grant 803151). T.N. was supported by JSPS KAKENHI JP19H05684 within JP19H05679. W.-J.L. was supported by the National Natural Science Foundation of China (grant no. 91951205 and 92251302). D.T. was supported by the Swedish Research Council (International Postdoc grant 2018-06609). C.W.S. was supported by a Science for Life Laboratory postdoctoral fellowship (awarded to T.J.G.E. and C.W.S.) and funding from the Swedish research council (Vetenskaprådet Starting grant 2020-05071 to C.W.S.). J.L. was supported by the Wenner-Gren Foundation (fellowship 2016-0072). J.H.S. was supported by a Marie Skłodowska-Curie IIF grant (331291). This work was also supported by the Moore-Simons Project on the Origin of the Eukaryotic Cell, Simons Foundation 73592LPI to T.J.G.E. and B.J.B. (https://doi.org/10.46714/735925LPI) and Simons Foundation 812811 to L.E. (https://doi.org/10.46714/735923LPI), and NSF Division of Biological Science SBS Biodiversity: Discovery and Analysis program (1753661) to B.J.B.

## Author contributions

T.J.G.E. conceived and supervised the study. A.S., K.U.K, W.H.L., Z.-S.H., A.L.R., W.-J.L., T.N., A.-L.R., M.B.S., and A.P.T. collected and provided environmental samples. E.F.C., F.H., J.H.S., N.D., K.W.S., B.J.B., L-X.C., J.F.B., and E.St.J. performed metagenomic sequence assemblies and metagenomic binning analyses. L.E., D.T., E.F.C., K.W.S., C.W.S., J.L., B.J.B., and T.J.G.E. analyzed genomic data. L.E., D.T., E.F.C. and F.H. performed phylogenomic analyses. L.E., D.T., E.F.C., C.W.S., J.L. and T.J.G.E. investigated ESPs. E.F.C., L.E., and M.E.S. performed ancestral genome reconstruction analyses. V.D.A., B.J.B., C.W.S, L.E. and T.J.G.E. carried out metabolic inferences. L.E., D.T., E.F.C., C.W.S., V.D.A, B.J.B. and T.J.G.E. wrote, and all authors edited and approved, the manuscript.

These authors contributed equally: Laura Eme, Daniel Tamarit, Eva F. Caceres.

## Extended data figures

**Extended Data Figure 1.**
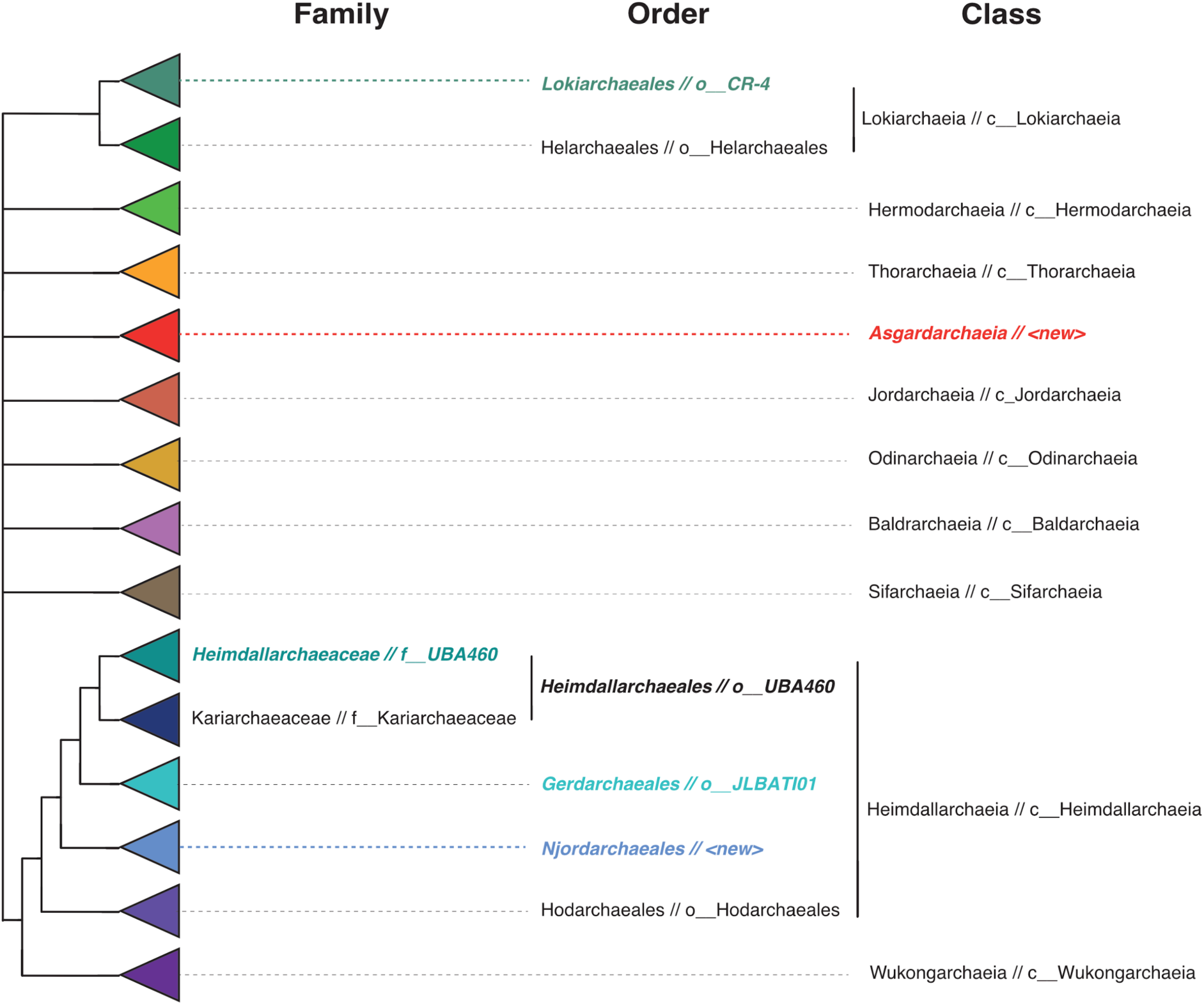
Cladogram of proposed taxonomic scheme for the ranks of family, order and class for Asgard archaeal lineages employed in this study. Equivalent names in GTDB are shown after a double slash (//). Cases with differing or new names have been highlighted in colored bold italics.

**Extended Data Figure 2.**
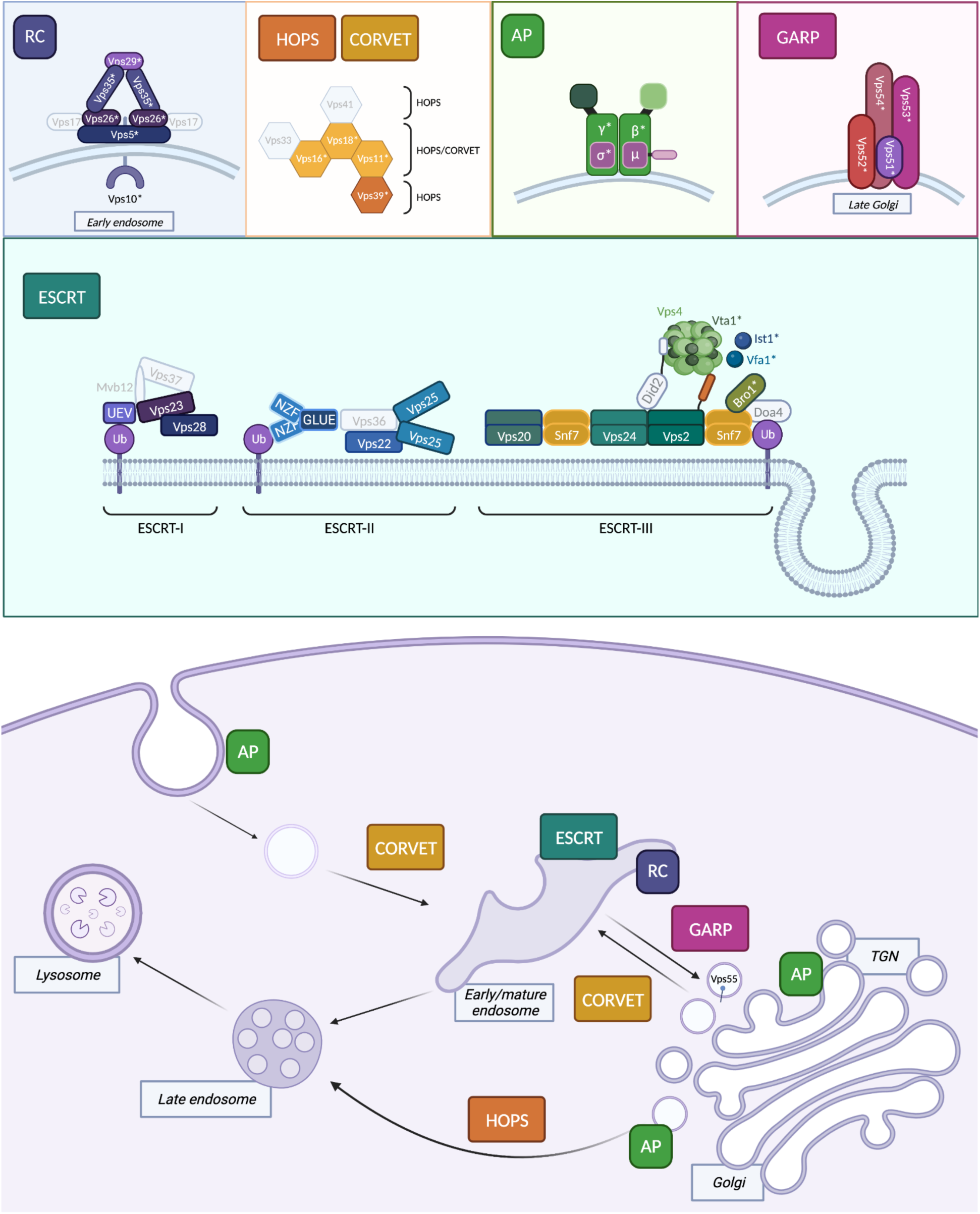
Identification of previously undetected vesicular trafficking ESPs in Asgard archaea. Schematic representation of a eukaryotic cell in which ESPs involved in membrane trafficking and endosomal sorting that have been identified in Asgard archaea are highlighted. Colored subunits have been detected in some Asgard archaea while grey ones seem to be absent from all current representatives. Only major protein complexes are depicted. Additional components can be found in Figure 2. From left to right, top to bottom: RC, Retromer complex. Retromer is a coat-like complex associated with endosome-to-Golgi retrograde traffic^36^. It is formed by Vacuolar protein sorting-associated protein 35, Vps5, Vps17, Vps26 and Vps29^127^. During cargo recycling, retromer is recruited to the endosomal membrane via the Vps5-Vps17 dimer. Cargo recognition is thought to be mediated primarily through Vps26 and possibly by Vps35. Finally, the BAR domains of Vps5-Vps17 deform the endosomal membrane to form cargo-containing recycling vesicles. Their distribution is sparse, but we have detected Asgard archaeal homologs of all subunits except for Vps17. Interestingly, the Thorarchaeota Vps5-BAR domain is often fused to Vps28, a subunit of the ESCRT machinery complex I, suggesting a functional link between BAR domain proteins and the thorarchaeial ESCRT complex. The best-characterized retromer cargo is Vps10. This transmembrane protein receptor is known in yeast and mammal cells to be involved in the sorting and transport of lipoproteins between the Golgi and the endosome. The Vps10 receptor releases its cargo to the endosome and is recycled back to the Golgi via the retromer complex^128^. CORVET: Class C core vacuole/endosome tethering complex; HOPS: Homotypic fusion and protein sorting complex. Endosomal fusion and autophagy depend on the CORVET and HOPS hexameric complexes^39^; they share the core subunits Vps11, Vps16, Vps18, and Vps33^40^. In addition, HOPS is composed of Vps41 and Vps39^41^. Vps39, found associated to late endosomes and lysosomes, promotes endosomes/lysosomes clustering and their fusion with autophagosomes^129^. AP, Adaptor Proteins. Asgard archaea genomes from diverse phyla encode key functional domains of the AP complexes. The eukaryotic AP tetraheteromeric structure is depicted, each color corresponding to a PFAM functional domain (Medium green: Adaptin, N terminal region; Dark green: Alpha adaptin, C-terminal domain; Light green: Beta2-adaptin appendage, C-terminal sub-domain; Dark pink/clear outline: Clathrin adaptor complex small chain; Light pink/dark outline: C-ter domain of the mu subunit); all five domains were detected in Asgard archaea, although not fused to each other. GARP: <u>G</u>olgi-<u>a</u>ssociated retrograde <u>p</u>rotein complex. The GARP complex is a multisubunit tethering complex located at the trans-Golgi network where it functions to tether retrograde transport vesicles derived from endosomes^37, 38^. GARP comprises four subunits, VPS51, VPS52, VPS53, and VPS54. ESCRT: Endosomal Sorting Complex Required for Transport system. This complex machinery performs a topologically unique membrane bending and scission reaction away from the cytoplasm. While numerous components of the ESCRT-I, II and III systems have been previously detected in Asgard archaea^9, 10, 42^, we here report Asgard homologs for several ESCRT-III regulators Vfa1, Vta1, Ist1, and Bro1. The bottom panel shows where these complexes mainly act in eukaryotic cells. Ub: Ubiquitin; Vps: <u>v</u>acuolar <u>p</u>rotein <u>s</u>orting. Subunit names in grey indicate that no homologs were detected in Asgard archaea. Domains newly identified as part of this study are indicated with an asterisk. Created with BioRender.com.

## Supplementary information

**Supplementary Information.** This file contains Supplementary Methods, Supplementary Discussions, Supplementary Figures 1-32, Supplementary Tables 1-8, Supplementary Data and Supplementary References.

Correspondence and requests for materials should be addressed to.

