## Supplementary Information for "Inference and reconstruction of the heimdallarchaeial ancestry of eukaryotes"

▲Equal contribution

†† Current address: Department of Biology, Lund University, Sölvegatan 35, 223 62 Lund, Sweden

††††† Current address: Department of Biochemistry, University of Cambridge, Cambridge, CB2 1QW, UK

### Table of contents


**Supplementary methods** 5

1. Functional annotation of protein clusters 5

2. Taxonomic and functional annotation of 'originations' in ancestors 5

3. Amino acid composition analyses 6

**Supplementary discussion** 8

1. Phylogenomics 8

1.1. Disentangling conflicting signals between the two gene marker sets 8

1.1.1. Unstable phylogenetic positions of eukaryotes and Njordarchaeales 8

1.1.2. The inclusion of Korarchaeota ribosomal protein sequences generates strong topological effects 9

1.1.3. Compositional adaptations to thermostability underlie phylogenetic conflict 11

1.2. Phylogenetic signal robustness assessment through systematic one-marker removal 15

1.3. Stable patterns after increased Asgard archaeal taxon sampling 16

1.4. Tackling long-branch effects reveals a nested eukaryotic placement within Heimdallarchaeia 18

1.5. Conclusions from phylogenomic analyses 20

1.5.2. Njordarchaeales represent a bona fide lineage of Asgard archaea 21

1.5.3. Eukaryotes robustly associate with Heimdallarchaeia 22

1.6. Comparisons with previous studies 23

2. Taxonomic classification of Asgard archaea: unification of novel high-ranking taxon names with the Genome Taxonomy Database taxonomic scheme 25

3. Optimal Growth Temperature prediction 29

4. Eukaryotic Signature Proteins 30

4.1. ESPs involved in informational processes 31

4.2. N-glycosylation (OST complex) 34

4.3. Vesicular trafficking and membrane remodelling 36

4.3.1. Adaptor protein complexes 36

4.4. Endosomal sorting 39

4.4.1. Retromer complex proteins Vps5, Vps26, Vps29 and Vps35 39

4.4.2. Vps10-Sortilin, a retromer cargo 40

4.4.3. Vps62 40

4.4.4. CORVET/HOPS complex proteins 40

4.4.5. Vps4 regulators: Vfa1, Vta1 and Ist1 41

5. Ancestral genome reconstruction 42

5.1 General considerations about the gene tree/species tree reconciliation approach used here 42

6. Genome content of the last archaeal common ancestor of the eukaryotes 45

6.1.1. The last archaeal ancestor of eukaryotes probably had a developed cytoskeleton but different from Lokiarchaeum 45

6.1.2. Metabolic features of Asgard archaeal ancestors 46

6.1.3. Proteins of bacterial origin represent a minority of ancestral Asgard archaeal proteomes 59

**Supplementary references** 61
**Supplementary figures 1-32** 70
**Supplementary tables** 114
**Supplementary data** 116

### Supplementary methods

#### 1. Functional annotation of protein clusters

Protein clusters have been aligned using mafft-linsi and used for profile-profile searches against a database that included profiles from NOGs, arCOGs and PFAM, using HHblits. Profile hits were considered if they showed a probability of being a true positive >= 0.95, a score_fraction >= 0.9 (i.e., the fraction of score with respect to the hit with the highest score) and an e-value <= 0.01. Such a filtering method allowed for local matches since no minimal length coverage of the query or template was required.

#### 2. Taxonomic and functional annotation of 'originations' in ancestors

We identified 2148 protein clusters that were inferred as an 'origination' (either transfers from outside the sampled archaeal species or *de novo* gene families) in any of 17 Asgard ancestral nodes displayed in Supplementary Figure 14. For 426 clusters that had a one-to-one correspondence to an EggNOG cluster at the LUCA level, we searched for the putative source of transfer similar to the approach described in [1]. We placed the sequences from these clusters onto the corresponding NOG trees using epa-ng[2], and extracted the most likely internal placement point. We also annotated all 'origination' clusters using the EggNOG v5 database and determined the affiliation to one of the broad functional categories used by EggNOG. Namely these are 'metabolism', 'cellular processes', 'information', 'unknown function', to which we added 'no homologies detected' (which differs from 'unknown function', where homologs are detected in the database but have no annotation that can be transferred) and the case where we found several of these categories to be present in the annotation (and therefore could not determine any one to be the 'correct' category).

#### 3. Amino acid composition analyses

Amino acid frequencies were calculated for all Njordarchaeales, other Heimdallarchaeia sequences and Korarchaeota sequences present in the RP and the NM dataset separately. We used those frequencies to carry out partition-around-medoids (PAM) clustering, principal component analysis (PCA) and their corresponding visualisation using R (https://www.R-project.org/) and the packages cluster and ggplot2 (Supplementary Figure 4). Thermostability-related amino acid metrics (the ratio of charged versus polar amino acids, and the fraction of residues represented by the amino acids isoleucine, leucine, valine, tryptophan, tyrosine, glycine, glutamate, arginine, lysine and proline) were calculated based on whole proteomes, genes within the RP dataset and genes within the NM dataset, and plotted them using ggplot2. To calculate thermostability-related metrics of alignment sites that favour two different topologies (monophyly of Njordarchaeales and Korarchaeota, versus monophyly of Njordarchaeales and other Heimdallarchaeia), we focused on the phylogenies obtained by running IQ-TREE2 using the LG+C60+G4+F+PMSF model on the RP56-A64--nDE and NM57-A64--nDE datasets. We obtained a consensus tree using IQ-TREE2 and keeping those branches where both phylogenies agreed. We then used this consensus tree to generate two constraints, one where Njordarchaeales were placed as the sister group of Korarchaeota, and another where they were placed as the sister group of other Heimdallarchaeia, reflecting the two topologies originally obtained by the RP56-A64--nDE and NM57-A64--nDE datasets, respectively. Finally, we used these two constrained trees to run IQ-TREE2 once again using both the RP56-A64--nDE and the NM57-A64--nDE datasets under the LG+C60+G4+F+PMSF model and printing the obtained site likelihood scores. These were used to identify which sites favoured one topology over the other. We calculated the thermostability-related amino acid compositional metrics of these sites, which were then plotted using ggplot2.

### Supplementary discussion

#### 1. Phylogenomics

##### 1.1. Disentangling conflicting signals between the two gene marker sets

###### 1.1.1. Unstable phylogenetic positions of eukaryotes and Njordarchaeales

Inferring evolutionary relationships between highly divergent lineages remains a major challenge in phylogenomics. As such, placing eukaryotes in relation to Archaea is a notoriously difficult object of study, which can only be assessed through careful methodology. But beyond this, other divergent (i.e., long-branching) lineages included in this work have been challenging to place. For example, the group to which we refer as Njordarchaeales has been placed within Asgard archaea[3] or as a sister group to Korarchaeota[4]. In the following sections, we put a strong emphasis on resolving the position of this group as it sometimes appeared as sister to eukaryotes in our reconstructions as well as in published phylogenies[3].

The phylogenies obtained using the untreated datasets (RP56-A64, RP56-A175, NM57-A64, and NM57-A175) were inconclusive with respect to the position of eukaryotes, and indicated important discrepancies with respect to the position of Njordarchaeales. Njordarchaeales appeared as a sister group to Korarchaeota in phylogenies obtained with RP56-A64 and RP56-A175, or together with Heimdallarchaeia in phylogenies obtained with NM57-A64 and NM57-A175, always with high bootstrap support (BS) (Supplementary Figure 17). In these phylogenies, eukaryotes appeared either affiliated with Asgard archaea (BS = 70% in RP56-A64, BS = 100% in NM57-A64 and NM57-A175) or with the TACK+Njordarchaeales group (BS = 88% in RP56-A175).

Given the notoriously long branches generated by DPANN sequences (which can generate phylogenetic reconstruction artefacts[5,6], we tested the effect of removing them from our dataset (RP56-A64-nD and NM57-A64-nD are versions of RP56-A64 and NM57-A64 without DPANN sequences). The topology obtained with NM57-A64-nD did not differ from that obtained with the original alignment (NM57-A64) (Supplementary Figure 2-3, 18). The topology obtained with RP56-A64-nD placed eukaryotes outside of the Asgard group, as sister to the TACK supergroup, albeit with low BS (Supplementary Figure 18), suggesting that the presence of DPANN sequences did not significantly affect the placement of eukaryotes and Korarchaeota. Consequently, in anticipation of more complex analyses, most subsequent datasets were generated without DPANN sequences.

Bayesian inference (BI) analyses did not produce converged chains after more than 20,000 iterations, and individual chains reflected the conflictual signal in the RP datasets. For example, when using the untreated RP56 concatenation (RP56-A64-nD) as input, all chains converged on the grouping of Njordarchaeales and Korarchaeota (Supplementary Figure 19). Meanwhile, the analyses of the SR4-recoded dataset (RP56-A64-nD-SR4) yielded two chains showing Njordarchaeales and Korarchaeota as sister groups, and the two remaining chains recovered the monophyly of Njordarchaeales together with Hodarchaeales, Gerdarchaeales and Heimdallarchaeales, nested within Asgard archaea (Supplementary Figure 20). The position of eukaryotes was unresolved.

###### 1.1.2. The inclusion of Korarchaeota ribosomal protein sequences generates strong topological effects

The topology obtained for the RP56 datasets is strongly affected by the removal of Korarchaeota (RP56-A64-nDK, RP56-A175-nDK), particularly regarding the position of Njordarchaeales. Using these datasets, we recover the grouping of Njord, Hod-, Gerd- and

Heimdallarchaeales (BS = 99%, RP56-A64-nDK) or Njord-, Hod-, Gerd-, Heimdallarchaeales and Wukongarchaeia (BS = 84%, RP56-A175-nDK), within Asgard archaea (Supplementary Figure 21). The position of eukaryotes remained unresolved, as they branched at the base of TACK+Asgard (RP56-A64-nDK) or TACK (RP56-A175-nDK) with low support (BS = 68% and 56%, respectively).

If we further exclude eukaryotes (RP-A64-nDEK, RP-A175-nDEK), the position of Njordarchaeales remains stable: Njord-, Hod-, Gerd-, and Heimdallarchaeales (and Wukongarchaeales, in the case of RP-A175-nDEK) cluster together (BS = 100%, for both RP-A64-nDEK and RP-A175-nDEK) (Supplementary Figure 22); this remained consistent and maximally supported in the SR4-recoded datasets analyses (Supplementary Figure 22).

In contrast, the position of Njordarchaeales inferred from the NM57 dataset variations appears more stable and not sensitive to the presence of Korarchaeota: ignoring the branching point of eukaryotes, the group formed by Njord-, Hod-, Gerd-, and Heimdallarchaeales consistently appears as monophyletic (with or without Wukongarchaeales), thus supporting the hypothesis that Njordarchaeales are *bona fide* Asgard archaea (Figure 2, Supplementary Figures 19-21, 23-25). In addition, the position of eukaryotes remained stable after the removal of Korarchaeota (NM57-A64-nDK and NM57-A175-nDK), as they clustered with Njordarchaeales (BS = 82% and BS = 98% for NM57-A64-nDK and NM57-A175-nDK, respectively) (Supplementary Figure 21).

Altogether, these results suggest that Njordarchaeales are artifactually attracted to Korarchaeota in the RP56 analyses. We observed that the Njordarchaeales and Korarchaeota RP56 homologs display similarly biased amino acid compositions typical of adaptation to hyperthermophily (see below), which led us to suspect that their shared lifestyle may explain their attraction to one another in the RP56-based phylogenetic analyses.

##### 1.1.3. Compositional adaptations to thermostability underlie phylogenetic conflict

###### 1.1.3.1. Growth temperature transitions and long branches

Asgard archaea include extremely diverse organisms, phylogenetically, metabolically and ecologically. Optimal growth temperature estimates (see Supplementary Discussion Section 3) indicate that some Asgard archaea thrive at low temperature, while others are thermophiles or even hyperthermophiles (Supplementary Table 5), in congruence with sample metadata (Supplementary Table 1). Lineages adapted to high temperatures are Jordarchaeia (median 65.4 °C), Wukongarchaeia (median 67.4 °C), Njordarchaeales (median 76.5 °C), and Baldrarchaeia (median 88.2 °C). Of these, however, only Wukongarchaeia and Njordarchaeales are separated by long branches from their closest relatives, particularly in the RP56-based phylogenies. For example, in the phylogeny obtained from the concatenated alignment NM57-A175-nDK, the stem branch of Njordarchaeales was 1.90 and 1.70 times longer than those of Jordarchaeia and Baldrarchaeia, and the equivalent branches in the phylogeny obtained from the RP56-A175-nDK alignment are 3.77 and 4.09 times larger for Njordarchaeales compared to Jordarchaeia and Baldrarchaeia.

Korarchaeota members are also known to be thermophilic[7,8]. Given these similarities in growth temperatures and the long branch at the base of Njordarchaeales, we hypothesized that their monophyly in certain phylogenetic reconstructions might have been caused by convergent amino acid composition patterns. Below, we performed compositional analyses to further investigate this.

###### 1.1.3.2. Compositional differences in protein marker sets

To visualize and interpret compositional data for the key taxa, we performed Partitioning Around Medoids (PAM) clustering, an unsupervised learning method, using the amino acid composition of the NM57 and RP56 proteins (NM57-A175 and RP56-A175 datasets) in Korarchaeota, Njordarchaeales and the traditionally named Heimdallarchaeia (here, Hodarchaeales, Gerdarchaeales Kariarchaeaceae and Heimdallarchaeaceae) (Supplementary Figure 17). In these representations, Hodarchaeales, Gerdarchaeales Kariarchaeaceae and Heimdallarchaeaceae form highly overlapping clusters, while Njordarchaeales and Korarchaeota cluster separately from them. In the plot corresponding to the RP56 dataset, a large overlap in amino acid composition is observed for the Njordarchaeales and Korarchaeota sequences, as these cluster altogether. Particularly, the first principal component (representing 60% and 47% of the variation present in the NM57 and RP56 sequences, respectively) shows Njordarchaeales and Korarchaeota clustering on the right-hand side of this axis, whereas the other Heimdallarchaeia cluster towards the left hand side (Supplementary Figure 4). This similarity in amino acid composition is possibly linked to adaptation to a (hyper)thermophilic lifestyle both in the Njordarchaeales and Korarchaeota lineages. A commonly observed compositional pattern for thermostability is the ratio between charged and polar amino acids[9–11]. We compared the values of this ratio between Hodarchaeales, Gerdarchaeales, Heimdallarchaeaceae, Kariarchaeaceae, Njordarchaeales and Korarchaeota (Supplementary Figure 5). Two-tailed t-tests (Supplementary Table 6) show that the differences between Hodarchaeales and the group formed by Gerdarchaeales+Kariarchaeceae+Heimdallarchaeceae are not statistically significant. However, the differences between both of these groups and Njordarchaeales and Korarchaeota are statistically significant for the NM57 and RP56 datasets (Bonferroni-corrected p-values $< 2.2e-16$). Yet, the difference between Korarchaeota and Njordarchaeales is not, or only marginally so (Bonferroni-corrected p-values = 0.79 and 1.3e-4, respectively). We also observed much higher ratio values for all four lineages in the RP56 proteins (mean±standard deviation: 1.94±1.05, 2.04±1.05, 2.58±1.25 and 2.93±1.80 for Hodarchaeales, Gerdarchaeales+Kariarchaeceae+Heimdallarchaeceae, Njordarchaeales, and

Korarchaeota, respectively) than in the NM57 proteins (1.36±0.40, 1.46±0.47, 2.04±1.21 and 2.04±0.86, respectively). More importantly, the difference and the spread of these values between (1) Hodarchaeales, Gerdarchaeales Kariarchaeaceae, and Heimdallarchaeaceae, and (2) the thermophilic Njordarchaeales and Korarchaeota, are stronger in the RP56 dataset compared to the NM57 dataset, consistent with a stronger bias for thermostability and a stronger impact on phylogenetic reconstructions.

A related but different metric for thermostability-related compositional bias is defined by the protein sequence fraction represented by the amino acids isoleucine, leucine, valine, tryptophan, tyrosine, glycine, glutamate, arginine, lysine, and proline (ILVWYGERKP)[11,12]. Similar to the previous metric, this fraction showed significant differences when comparing Hodarchaeales and Kari+Gerd+Heimdallarchaeales, to both Njordarchaeales and Korarchaeota, but not when comparing Njordarchaeales and Korarchaeota (Supplementary Table 6). This fraction also yielded higher differences and standard deviation in the RP56 dataset compared to the NM57 dataset (Supplementary Figure 5).

Finally, it is known that adaptation to (hyper)thermophily, while it impacts the whole genome[13], leads to a stronger composition bias in the tRNAs[14] and rRNAs, which tend to be more GC-rich than the rest of the genome[15]. The ribosome, by interacting intimately with those RNAs logically coevolve with them[16], presenting similarly stronger composition bias than the rest of the proteome and likely accumulating convergent adaptations in unrelated thermophilic lineages.

Altogether, this suggests that the attraction between Njordarchaeales and Korarchaeota in the phylogenetic reconstructions based on the RP dataset is, at least in part, the result of this compositional similarity.

###### 1.1.3.3. Site-likelihood analyses reveal artifactual topologies caused by compositionally biased sites

To further confirm the source of conflict between the two sets of markers, we identified the sites in the NM57-A64 and RP56-A64 concatenations that preferentially supported one topology over the other (i.e., whose likelihood was higher for a topology where Njord branched with Hod+Gerdarchaeales+Kari+Heimdallarchaeaceae than for one where Njordarchaeales branched with Korarchaeota, or *vice versa*). By doing so, we observed that, overall, the conflictual signal can be observed in both datasets, each containing a high number of sites supporting either topology. However, we observed that the NM57 concatenation includes ~1.3 times more sites supporting the topology where Njordarchaeales cluster with Hod+Kari+Gerd+Heimdallarchaeales topology than sites supporting the monophyly of Njordarchaeales and Korarchaeota. In contrast, the number of sites supporting one topology or the other is almost identical in the RP56 dataset (2834 versus 2913, ratio of 0.97, for Njord+Hod+Kari+Gerd+Heimdallarchaeales and Njordarchaeales+Korarchaeota, respectively).

We then calculated thermostability-related amino acid compositional patterns in the sites that favoured either the monophyly of Njordarchaeales and Korarchaeota, or the monophyly of Njordarchaeales and Hod+Kari+Gerd+Heimdall (Supplementary Figure 6). The ratio of charged *versus* polar amino acids in Njordarchaeales genomes was higher, both in the RP56 and NM57 gene markers, at sites favouring the monophyly of Njordarchaeales and Korarchaeota compared to sites that favoured the monophyly of Njordarchaeales and other Heimdallarchaeia (t-test p-values of 0.0011 and 1.28e-5 for NM57 and RP56 gene datasets, respectively). Moreover, this ratio was notably higher in the RP56 dataset (median for sites that supported Njordarchaeales+Korarchaeota=2.37; median for sites that supported Njordarchaeales+Heimdallarchaeia=2.09) than in the NM57 dataset (1.86 and 1.75, respectively). Similarly, the fraction ILVWYGERKP also showed significantly higher values in sites favouring the monophyly of Njordarchaeales and Korarchaeota (t-test p-values of 2.5e-12 and 7.78e-15 for NM57 and RP56 datasets, respectively), and consistently showed higher values in the RP56 dataset (median for sites that supported Njordarchaeales+Korarchaeota=0.71; median for sites that supported Njordarchaeales+Heimdallarchaeia=0.66) compared to the NM57 dataset (0.66 and 0.63, respectively).

These results are also consistent with a scenario where both sets of proteins have identical evolutionary histories, but have been under different evolutionary pressures for thermostability. Consequently, the ribosomal proteins display a stronger compositional bias compared to the rest of the proteome. We thus interpret that the NM57 dataset carries a more reliable phylogenetic signal when it comes to the position of Njordarchaeales. Based on these analyses and the phylogenomic results shown above (see 4.5), we conclude that Njordarchaeales affiliate with Hodarchaeales, Gerdarchaeales, and Kari+Heimdallarchaeaceaea, in a group forming the Heimdallarchaeia.

##### 1.2. Phylogenetic signal robustness assessment through systematic one-marker removal

To further study the robustness in phylogenetic signal carried by each gene set (NM57 and RP56), we inferred phylogenies from versions of the NM57-A175-nDK and RP56-A175-nDK datasets in which we systematically removed one protein at a time from the concatenation (i.e., we generated 57 supermatrices derived from the NM57-A175-nDK dataset based on all possible concatenations of 56 proteins out of 57; and 56 supermatrices derived from the RP56-A175-nDK dataset, corresponding to all possible concatenations of 55 of the 56 ribosomal proteins) (Supplementary Table 2). This allows to investigate how congruent the signal is across individual markers.

All supermatrices corresponding to the NM57 dataset subconcatenations yielded very consistent topologies. For example, all resolved with high support the monophyly of eukaryotes and Njord (85-100% BS), and the monophyly of eukaryotes, Heimdallarchaeia (Hod-, Gerd-, Njordarchaeales, and Kari- and Heimdallarchaeaceae) and Wukongarchaeales (84-100%). Moreover, the overall Asgard archaeal topology was largely consistently resolved.

On the other hand, the phylogenies obtained from the RP56 dataset variations produced highly unresolved and inconsistent results. For example, none of the phylogenies generated by any of the 55 RP variants resolved the position of eukaryotes in a supported (i.e., >= 70% BS) monophyletic group with Asgard archaea or TAC archaea. Additionally, 4 phylogenies showed moderately high support (70-85% BS) for the monophyly of TAC and Asgard archaea to the exclusion of eukaryotes. Moreover, the position of Njordarchaeales was unstable, branching with high support (>= 70%) with other Heimdallarchaeia and Wukongarchaeales in 23 of the 56 phylogenies, and with eukaryotes, outside of the Asgard archaeal group, in 8 phylogenies.

This analysis further indicates that the combination of low phylogenetic signal and excess of compositional biases make the RP56 a poorer gene marker set for phylogenomic analyses of eukaryotic and archaeal evolution, compared to the NM57 gene marker set.

##### 1.3. Stable patterns after increased Asgard archaeal taxon sampling

All the phylogenomic analyses presented above were performed on sequences obtained from two main taxon selections, both of which included Asgard archaea, DPANN archaea, Euryarchaeota, TACK archaea and eukaryotes. The first taxon sets included 68 Asgard archaea (published before March 2019), while the second included 175 Asgard archaea (representatives of published sequences as of June 2021).

Using the smaller taxon sampling to generate the NM57-A64 and RP56-A64 concatenations, we explored the effect produced by the presence of various groups that were typically characterised by long branches or unstable positions, such as DPANN, Korarchaeota, eukaryotes, or Njordarchaeales (Supplementary Table 2). As discussed already in parts, the NM57-A64 taxon sampling variants overall generated highly consistent results. For example, Njordarchaeales consistently associated with the Heimdallarchaeial group (Hod- and Gerdarchaeales, and Kari- and Heimdallarchaeaceae) in a monophyletic group that included eukaryotes whenever they were present in the analysis. This monophyly was routinely supported with high bootstrap values (generally equal to 100%). When both groups were present, eukaryotes associated with Njordarchaeales (BS > 81%), and jointly formed a monophyletic group with Heimdallarchaeia (BS > 97%). When Njordarchaeales were absent, eukaryotes still formed a monophyletic group with Heimdallarchaeia (BS = 100%). Other groups consistently found monophyletic were the whole Asgard archaeal clade (BS = 100%). In contrast, the RP56-A64 variants produced incongruent topologies. One such example was discussed above, where Njordarchaeales associated with Korarchaeota when both groups were present (BS > 91%), but branched with Heimdallarchaeia (plus eukaryotes if present) when Korarchaeota were absent (BS > 94%). Additionally, eukaryotes associated either with Heimdallarchaeia (BS = 100% in RP56-A64, BS = 92% in RP56-A64-nDNK –no DPANN, Njordarchaeales or Korarchaeota–), the Njord+Heimdallarchaeia group (BS = 99% if only including Asgard archaea and eukaryotes), or an unsupported position outside of Asgard archaea (BS < 70% in when no DPANN were included, with or without Korarchaeota and Njordarchaeales – RP56-A64-nD, RP56-A64-nDK, RP56-A64-nDN). The monophyly of Asgard archaea (with or without Njord) was generally unsupported (BS < 70%).

The second dataset we explored had a higher representation of Asgard archaea (175 taxa), and therefore a larger number of sequences overall. Given this and the high computational resources needed for the previous set of phylogenomic analyses, we generated a smaller number of taxon subsets based on these larger datasets. However, the general patterns we observed were very similar. For example, the NM57-A175 dataset variants generated phylogenies with topologies in which the Heimdall-, Njord- and Wukongarchaeales (the latter group was not included in the A64 datasets, due to their recent discovery[17]) formed a monophyletic group, with eukaryotes when present, with high support (BS > 91%). The RP56-A175 dataset instead generated phylogenies with an unsupported (BS < 70%) position of eukaryotes outside of Asgard archaea. RP56-A175 variants did not consistently resolve the position of Njordarchaeales, which associated with Korarchaeota when both groups were present (BS = 95%), and with Heimdallarchaeia when Korarchaeota was absent (BS > 83%).

##### 1.4. Tackling long-branch effects reveals a nested eukaryotic placement within Heimdallarchaeia

Based on all the above, we concluded that Njordarchaeales had been artifactually attracted to Korarchaeota due to convergent thermostability sequence adaptation, but instead belonged within Heimdallarchaeia. Additionally, in the phylogenies obtained with the untreated NM57-A175 and NM57-A175-nDK datasets, eukaryotes affiliated with the group formed by Heimdallarchaeia (including Njordarchaeales) and Wukongarchaeales. More specifically, these two phylogenies showed monophyly of Njordarchaeales and eukaryotes with high bootstrap support (98% in both cases). However, the eukaryotes position was not resolved in the phylogenies obtained with the untreated RP56-A175 and RP56-A175-nDK datasets.

To further investigate the position of eukaryotes, we aimed to alleviate potential artefactual signals commonly carried by fast-evolving sites. These are generally poorly modeled by standard reversible substitution models and can lead to artefactually longer branch estimates. We thus employed two main data treatments: SR4 recoding[18] and Fast-Site Removal (FSR).

Both aim to ameliorate potential phylogenetic artefacts arising from model misspecification at mutationally saturated or compositionally biased sites[19–22]. The SR4-recoded phylogenies were reconstructed with IQ-TREE (using a user-defined previously described model referred to as 'C60SR4', based on the implemented 'LG+C60' model and modified to analyze the recoded data[23]) and Phylobayes (under the CAT+GTR model)[23]. FSR datasets were generated by using the estimated site rate output by IQ-TREE to classify sites into 10 categories, from the fastest to the slowest evolving; we removed them in a stepwise fashion, removing from 10% to 90% of the data.

Disentangling the effect of taxon sampling, FSR and SR4-recoding is not straightforward. However, a common pattern arose after using FSR and/or SR4-recoding treatments on the NM57-A175-nDK dataset (without DPANN or Korarchaeota), where eukaryotes were robustly placed within Heimdallarchaeia, either with the Njordarchaeales or the Hodarchaeales groups. Unfortunately, the position of eukaryotes remained unresolved in the phylogenies obtained from treated versions of the RP56-A175-nDK alignment. As an illustration of the effect of FSR and SR4-recoding on the studied phylogenies, we have mapped the evolution of bootstrap support for the monophyly of either (1) eukaryotes and Njordarchaeales or (2) eukaryotes and Hodarchaeales, in phylogenies obtained from untreated and treated versions of the NM-A175-nDK dataset (Supplementary Figure 23, Supplementary Table 2). The effect of FSR is not linear, but we can nevertheless observe, for the NM57 datasets, that stepwise removal of the fastest-evolving sites yielded lower bootstrap support for eukaryotes branching with Njordarchaeales.

In contrast, the effect of SR4-recoding was much more consistent. The monophyly of Njordarchaeales and eukaryotes is never supported in SR4-recoded versions of the NM57-A175-nDK dataset. Meanwhile, the support for the monophyly of eukaryotes and Hodarchaeales increased. This is well displayed in SR4-recoded versions of the NM57-A175-

nDK alignment where 20% and 40% of the fastest-evolving sites were removed (Supplementary Figures 24 and 25). In the phylogenies obtained from these alignments, the monophyly of eukaryotes and Hodarchaeales was obtained with 71% bootstrap support. In phylogenies of both recoded and non-recoded alignments, removing over 50% of the fastest-evolving sites resulted in major loss of phylogenetic signal and consequent loss of support for monophyletic groups containing eukaryotes and any specific groups. All these trends are consistent with the results obtained from taxon sampling variations of the NM57-A64 dataset (NM57-A64-nD: without DPANN; NM57-A64-nDK: without DPANN and Korarchaeota; NM57-A64-nDN: without DPANN and Njordarchaeales; NM57-A64-nDNK: without DPANN, Njordarchaeales or Korarchaeota; NM57-A64-AsgE: only including Asgard archaea and eukaryotes).

The various treatments, when employed on the RP56-A175-nDK dataset or taxon sampling variants of the RP56-A64 dataset, did not help resolve the position of eukaryotes.

Combining FSR and recoding also showed an additional, clear trend with respect to the position of Njordarchaeales within Heimdallarchaeia, when using NM57-A175 and RP56-A175 alignment variants. In the absence of Korarchaeota, FSR-treated alignments (at shallow levels between 20-60% in recoded alignments, and deeper levels between 50-90% in non-recoded alignments) consistently placed Njordarchaeales as a sister group to the group formed by Gerd-, Kari- and Heimdallarchaeales. In these phylogenies, Njordarchaeales would not be the most divergent group within Heimdallarchaeia, but well nested within them.

#### 1.5. Conclusions from phylogenomic analyses

##### 1.5.1. Ribosomal proteins are artefact-prone gene markers in the presence of strong compositional biases

We performed compositional analyses that showed that the RP56 dataset carries stronger compositional differences between thermophilic (Njordarchaeales and Korarchaeota) and mesophilic (Hod-, Gerd-, Kari-, Heimdallarchaeales) lineages compared to the new marker dataset (NM57). This was reflected in overall amino acid usage as seen through principal component analysis and known compositional features related to thermostability. Moreover, site-likelihood analyses indicated that the NM57 dataset included a larger proportion of sites supporting a single topology in which Njordarchaeales are monophyletic with the rest of Heimdallarchaeia, while the RP56 dataset had a slightly larger number of sites supporting the monophyly of Njordarchaeales and Korarchaeota. Furthermore, the sites supporting the topology in which Njordarchaeales and Korarchaeota clustered together were significantly enriched in "thermostable" amino acids, suggesting that these sites were the results of convergent evolution.

##### 1.5.2. Njordarchaeales represent a *bona fide* lineage of Asgard archaea

Phylogenomic analyses of the ribosomal protein dataset (RP56) did not robustly place Njordarchaeales in the archaeal tree, as their position was heavily dependent on taxon sampling strategies. On the other hand, the new marker dataset (NM57) robustly placed Njordarchaeales within Asgard archaea, forming a monophyletic group with other Heimdallarchaeia. Given the previously described lines of evidence, we concluded that the placement of Njordarchaeales with Korarchaeota is the result of strong convergent compositional biases. Thus, indications that Njordarchaeales are indeed close relatives of Heimdallarchaeia were observed through the use of conserved non-ribosomal proteins, and the removal of Korarchaeota when using the ribosomal protein dataset. Moreover, the use of a more thorough taxon sampling (A175), following the publication of a large number of Asgard archaeal metagenome-assembled genomes in recent years, allowed a more precise placement of Njordarchaeales within Asgard archaea. In these phylogenies, Njordarchaeales was consistently placed nested within the Heimdallarchaeia, as a sister to the group formed by Gerdarchaeales, Kariarchaeaceae, and Heimdallarchaeaceae.

##### 1.5.3. Eukaryotes robustly associate with Heimdallarchaeia

Resolving the position of eukaryotes in the tree of life remains one of the most challenging themes in phylogenomics. In our analyses, eukaryotes were consistently placed within Asgard archaea, with few unsupported exceptions. More specifically, we retrieved very high support for topologies in which eukaryotes formed a monophyletic group with Heimdallarchaeia (Hod-, Gerd-, and Njordarchaeales, and Kari- and Heimdallarchaeaceae). Phylogenies obtained from untreated datasets often placed eukaryotes as a sister-group to Njordarchaeales (see section 1.1.1). However, phylogenies generated from alignments that were specifically treated to alleviate mutational saturation interestingly displayed a less strong affiliation between eukaryotes and Njordarchaeales, and a stronger affiliation between eukaryotes and Hodarchaeales. We interpret that the longer branch at the base of Njord, a lineage characterized by a strong compositional modification of the proteome caused by adaptation to (hyper)thermophily, was the result of poor model fit and yielded long-branch attraction between eukaryotes and Njordarchaeales. The analysis of slower evolving/recoded sites thus revealed a clearer affiliation between eukaryotes and Hodarchaeales, a lineage characterized by large genomes with a high number of Eukaryotic Signature Proteins (see section 3). While this specific affiliation between eukaryotes and Hodarchaeales still requires corroboration from additional studies, the nested branching of eukaryotes within the larger Heimdallarchaeia group is robustly supported by the phylogenomic investigations presented here.

##### 1.6. Comparisons with previous studies

Multiple phylogenomic analyses attempting to place eukaryotes in the archaeal tree of life have been published since the discovery of Asgard archaea. The pioneering study by Spang and colleagues[24] described the first Asgard archaeal MAGs, which were classified at the time as Lokiarchaea (here, Lokiarchaeales), and through phylogenomic analyses of a variety of single copy marker genes, showed the relatedness between eukaryotes and Lokiarchaea. Later in 2017, Zaremba-Niedzwiedzka et al[23] published a study in which phylogenetic analyses including representatives of additional Asgard groups named Thorarchaeota, Odinarchaeota and Heimdallarchaeota (here, Thorarchaeia, Odinarchaeia, and Heimdallarchaeia, respectively) were performed. Phylogenomic analyses on a set of 55 ribosomal proteins resulted in high confidence for the monophyly of Asgard archaea and eukaryotes[23]. In this study, both maximum-likelihood and Bayesian phylogenies obtained indicated (albeit with low support) monophyly of eukaryotes with Heimdallarchaeota (here, Heimdallarchaeia).

Other analyses, including Seitz et al. (2019)[25], Spang et al. (2019)[26], Liu et al. (2021)[17], Sun et al. (2021)[27], and Wu et al. (2022)[28] also recovered a relationship between eukaryotes and various Heimdallarchaeia clades, although only weakly supported. Moreover, Xie et al (2022)[3] placed eukaryotes as sister to the Njordarchaeales, also within Heimdallarchaeia. These studies, which were all based on a less broad Asgard archaeal taxon sampling, are consistent with the main findings reported in the present study (i.e., high confidence for the monophyly of eukaryotes and Heimdallarchaeia, and moderate support for Hodarchaeales as closest archaeal relatives of eukaryotes).

Some other studies have produced results that do not match our conclusions. In particular, Da Cunha and colleagues placed eukaryotes outside of Archaea altogether[29,30]. However, their arguments were disproved elsewhere[22,31]. In particular, Williams and colleagues reanalysed the 35-gene matrix of Da Cunha et al.[29] using best-fitting models in both maximum-likelihood and

Bayesian analyses and recovered the strongly supported monophyly of Asgard archaea and eukaryotes. Moreover, Bayesian inference of the dataset of Da Cunha et al under complex mixture models (CAT+GTR+G4) placed eukaryotes within Heimdallarchaeota (here, Heimdallarchaeia) in this study, although sister to the Kariarchaeaceae (only represented by one genome)[22]. Finally, the recent study by Aouad and colleagues[32] has yielded high support for the sister-relationship between Asgard archaea and eukaryotes, but their taxon selection of Asgard archaea was rather restricted, only including the 9 Asgard archaea MAGs from Zaremba-Niedzwiedzka et al. (2017)[23].

Some of the previously mentioned studies above also generated topologies showing a sister relationship between Asgard archaea and eukaryotes, such as Liu et al (2021)[17] when using a consensus topology of 129 trees containing bacteria, archaea and eukaryotes, constructed from the concatenated protein sequence alignments of bootstrap-like samples and leave-one-out sets of a small set of 29 universally conserved markers. In this study, only the site-homogeneous LG+R10 phylogenetic model was used to generate phylogenetic trees. The phylogeny yielded from a concatenation of these 29 markers resulted in a deeper-branching position of eukaryotes within Archaea, which we suspect is the result of a long-branch attraction artefact to the distantly related bacterial outgroup combined with the poor fit of the simple LG+R10 substitution model.

More recently, Rodrigues-Oliveira et al. (2022)[33] replicated this phylogeny by using 23 of the 29 markers from the same non-Asgard archaeal genomes as in Liu et al. 2021[17] plus a set of 94 diverse Asgard archaea, this time using the mixture model LG+C20+G4+F. They recovered here a moderately supported sister relationship between Asgard archaea and eukaryotes (ultrafast bootstrap support 84%).

We argue that the key differences between our study and those mentioned above include (1) the broad taxonomic representation of Asgard archaea, (2) the complex site-heterogeneous evolution models we employed (LG+C60+G4+F in maximum likelihood and CAT+GTR or CAT+LG model in Bayesian inferences, which allow the amino acid replacement patterns at different sites of a protein alignment to be described by distinct substitution processes), (3) the various alignment treatments we performed to alleviate potential long-branch attraction artefacts, and (4) the complex analyses we made to disentangle the signal yielded by compositional biases.

Altogether, the phylogenomic results shown here robustly place eukaryotes in a monophyletic group with Heimdallarchaeia and suggest a nested position as sister to the Asgard archaeal class Hodarchaeales (see next section).

#### 2. Taxonomic classification of Asgard archaea: unification of novel high-ranking taxon names with the Genome Taxonomy Database taxonomic scheme

Microbial taxonomy is currently undergoing profound changes, with conflicting taxonomic schemes provided by different parties. Originally, Asgard archaea was defined as a superphylum including multiple phyla (originally Lokiarchaeota, Thorarchaeota, Heimdallarchaeota, and Odinarchaeota[23], soon completed with Helarchaeota[25]). A different proposal has been put forward by the Genome Taxonomy Database (GTDB)[34,35], using *ad hoc* thresholds for the highest taxon ranks, and generating taxonomic categories based on a normalised, simple evolutionary tree (see Sun et al. 2021[27] for a recent use of this taxonomic scheme). According to this taxonomic scheme, Asgard archaea becomes a phylum ("Asgardarchaeota"), and the main Asgard archaeal divisions become classes (e.g., Lokiarchaeia, Thoarchaeia, Heimdallarchaeia, Odinarchaeia). To avoid perpetuating confusion, we have initiated a community effort to reconcile these conflicting taxonomic schemes into a unified Asgard archaeal taxonomy (Tamarit, Eme, Ettema *et al.*, in prep.).

We strongly believe that the phylogeny used as a basis to define an adequate taxonomic scheme for Asgard archaea will require additional attention (as detailed at length previously). However, we acknowledge the effort made by the GTDB to standardize phylogenetic ranking and the traction it has gained in the microbiology community. Consequently, here, we favour and expand on a taxonomic scheme compatible with the GTDB classification system (Ext. Data Fig. 1). As such, Asgard archaea would be defined at the phylum level (Asgardarchaeota) and be divided into the classes Lokiarchaeia, Thorarchaeia, Hermodarchaeia, Baldrarchaeia, Jordarchaeia, Asgardarchaeia, Sifarchaeia, and Heimdallarchaeia.

Recently, Liu and colleagues[17] proposed dividing the previously named Heimdallarchaeota into four phyla: Gerdarchaeota, Kariarchaeota, Heimdallarchaeota (thereby now confusedly formed only by a subset of the lineages previously known as Heimdallarchaeota) and Hodarchaeota. Given the clear monophyly of these groups and the ranking of this entire clade as 'class' according to the GTDB scheme, we suggest referring to them as Heimdallarchaeia, as currently proposed by the GTDB Release 207. The subgroups within Heimdallarchaeia would be considered under the taxonomic rank of 'order', i.e. Gerdarchaeales, Heimdallarchaeales, and Hodarchaeales, and 'family', i.e. Heimdallarchaeaceae and Kariarchaeaceae (both within Heimdallarchaeales; Tamarit, Eme, Ettema *et al.*, in prep.). Moreover, given our robust placement of Njordarchaeales (previously Njordarchaeota[3]) within Heimdallarchaeia, we also assign them the rank of 'order' and name them Njordarchaeales in this manuscript. The group here named Wukong (previously Wukongarchaeota[17]) did not have a robust placement in our phylogenomic analyses, often affiliating within Heimdallarchaeia or as sister to them. This group is currently only formed by two highly similar MAGs, separated from other Asgard lineages by a long branch. For this reason, we predict that they represent a single order (Wukongarchaeales) but concluding on whether this group belongs to Heimdallarchaeia or to a separate class (Wukongarchaeia) would require further analyses and additional genomic data.

Another difference between the taxonomy scheme employed here and the one in Liu et al. (2021)[17] refers to the group named Borrarchaeota in the latter study, defined as the phylum-level taxon containing *Borrarchaeum yapensis*. Here, we acknowledge the anteriority of the article by Farag and colleagues (2021)[36], which introduced the phylum Sifarchaeota, based on two bins used to define the genus *Sifarchaeotum*. Given the large distance between the two *Sifarchaeotum* genomes in our phylogenies (e.g. Supplementary Figure 26), it seems unlikely that they represent organisms that belong to the same genus. However, it does seem likely that they represent organisms related at a class level, as proposed in Sun et al. (2021)[27]; we here refer to them as Sifarchaeia (also see Tamarit, Eme, Ettema *et al.*, in prep.). While Sifarchaeia may be further subdivided into two orders (which could possibly be called Sifarchaeales and Borrarchaeales in recognition of the two previously cited studies), this difference is too fine-grained to be adequately resolved with the taxon sampling employed in the current study.

Another recent study, Xie et al. (2022), proposed the phyla Njordarchaeota, Sigynarchaeota and Freyrarchaeota. We have discussed extensively about the first group, which here we name Njordarchaeales. Additionally, while we have not included the genomes that Xie et al. used to define the Sigynarchaeota and Freyrarchaeota, we note that the former is clearly nested within the Lokiarchaeia (hence an invalidly proposed taxon) while the latter corresponds to the Jordarchaeia[27] (Tamarit, Eme, Ettema *et al.*, in prep.).

The taxonomic scheme proposed here thus coincides with the one proposed by the GTDB. There are, however, five notable exceptions: the groups here named Heimdallarchaeaceae ("f__UBA460" in GTDB), Gerdarchaeales ("o__JLBATI01" in GTDB), Lokiarchaeales ("o__CR-4" in GTDB), Njordarchaeales (no equivalent in GTDB) and Asgardarchaeia (no equivalent in GTDB) (Extended Data Figure 1). The first three groups result from the reclassification of previously proposed groups. To anchor them to specific genome representatives, here we propose that Heimdallarchaeaceae be represented by strain ABR16

(93.46% completeness and 3.27% contamination according to CheckM); that Gerdarchaeales be represented by strain SZ_4_bin5.60 (93.46% completeness and 8.88% contamination according to CheckM); and that Lokiarchaeales be represented by strain WORC5 (87.38% completeness and 3.27% contamination according to CheckM). For the first two strains we thus propose the candidate genus names *Ca.* Heimdallarchaeum and *Ca.* Gerdarchaeum, and in the latter case we classify strain WORC5 within the already proposed genus *Ca.* Lokiarchaeum[24]. Strain WORC5 is closely related to *Ca*. Lokiarchaeum sp. GC14_75, the first named Asgard archaeon found in the vicinity of Loki's castle (see e.g. Supplementary Figure 16). We calculated the Average Nucleotide Identity score (based on the calculator by Varghese et al. (2015)[37]) between strains GC14_75 and WORC5, obtaining a value of 74.5% with an alignment fraction of 0.33. These are well within the boundaries defined for genus demarcation according to Barco et al. (2020)[38]. Additionally, Njordarchaeales had been previously proposed as Njordarchaeota but with no ascribed representative genome – here, we propose strain GBS24 (92.06% completeness and 4.67% contamination according to CheckM) as representative genome.

The novel group named Asgardarchaeia (Tamarit, Eme, Ettema *et al.*, in prep.) is represented in our study by a single genome, strain B16_G1 (GCA_003662835.1). The lack of close relatives results in a long branch, difficult to place with confidence. Moreover, this MAG is unfortunately relatively incomplete (CheckM[39]: 59.51% completeness) with an estimated contamination level of 5.61% by CheckM. For these reasons, we were unable to robustly place it in our phylogenies. Moreover, although strain B16_G1 was confidently placed within Asgard archaea in most of our phylogenies, in some cases the presence of contaminating marker proteins caused this MAG to fall outside of the Asgardarchaeota clade (e.g. see Supplementary Figure 2). A robust phylogenomic analysis using additional newly sequenced genomes from this group has placed it confidently as a novel class-level taxon within Asgard archaea

(Tamarit, Eme, Ettema *et al.*, in prep.). We surveyed the NCBI database for recently published Asgard genomes and found three MAGs published as "Odinarchaeota" in December 2021 (AUK265: GCA_021160805, AUK159: GCA_021162905 and AUK204: GCA_021161985) that form a supported monophyletic group with B16_G1 in phylogenomic trees using the RP gene dataset (Supplementary Figure 27) and in most trees constructed from individual RP and NM gene markers (available in Figshare). Trees constructed from individual NM proteins show that sequences from 9 out of the 29 markers (M030, M032, M037, M039, M045, M060, M069, M212, MA56) identified in the B16_G1 MAG cluster with euryarchaeal sequences with strong support and short branches, while AUK265, AUK159 and AUK204 cluster together, with the rest of Asgard archaea. This probable contamination seemingly generates enough conflicting signal that B16_G1 is often misplaced in phylogenies constructed from concatenated NM markers. However, whenever B16_G1 is correctly placed within Asgard archaea, this genome is placed in a well-supported monophyletic group with genomes AUK265, AUK159 and AUK204, which themselves are consistently placed within Asgard archaea in phylogenies constructed from either RP or NM markers (Supplementary Figure 27). For this reason, we are confident that B16_G1, AUK265, AUK159 and AUK204 jointly form a novel class-level taxon within Asgard archaea.

#### 3. Optimal Growth Temperature prediction

Optimal growth temperatures were predicted for the genomes presented here based on genomic and proteomic features[11] (Supplementary Table 5). The used method applies an empirical equation obtained by regression analysis using a set of archaeal genomes for which the corresponding host optimal growth temperature is known. Since this set is dominated by

Crenarchaea and Euryarchaeota, estimates for Asgard archaea remain an estimate that will require experimental validation. Since ribosomal RNAs nucleotide composition are used in this method, only genomes with predicted rRNAs were analyzed. Because these predictions are made only based on a subset of genomes for each phylum, they may not reflect the full range of temperatures at which these organisms can live. Predictions based on genomic features are sometimes at odds with the reported sample temperatures, particularly for those coming from Guaymas Basin (Supplementary Table 1). This can be explained first because temperatures are measured using a probe *near* the sediment cores (not in the cores themselves). Second, we know that there can be large temperature gradients across short distances in those sediments, for example due to hot fluids rising in plumes in the sediments. However, the presence of the toprim DNA reverse gyrase allows us to be confident that Jord- and Baldrarchaeia and Wukong- and Njordarchaeales are *bona fide* hyperthermophiles.

#### 4. Eukaryotic Signature Proteins

We first updated the distribution of previously identified ESPs, including eukaryotic RLC7 family proteins, actin homologs, gelsolin-domain proteins, components of eukaryotic ESCRT (I, II and III) systems including two sub-families of SNF7 proteins, ubiquitin modifier system components and homologs of eukaryotic protein translocation and glycosylation pathways, and found that most of these are widely distributed across sampled Asgard archaeal lineages (Figure 3). Below we describe some of these patterns in more detail, and we report newly identified ESP homologs in Asgard archaea. Accession numbers for these are available in Supplementary Table 3.

#### 4.1. ESPs involved in informational processes

##### 4.1.1. RNA polymerase A

We reconstructed the evolutionary history of the RNA polymerase A (RNApol A) among archaea (Fig. 41.1.1_01 and 41.1.1_02), which exists in two versions: encoded by a single gene ("fused RNApol A"), or by two neighbouring genes ("split RNApol A", which correspond to the subunits A' and A''), whose distributions across organisms suggest a complex evolutionary history (Supplementary Figure 28)[23,31]. Eukaryotes, Thaum-, Aig-, Kor- and Bathyarchaeota (belonging to the TACK superphylum) encode a fused RNApol A, whereas Euryarchaeota, Crenarchaeota (incl. Geoarchaeales), and most DPANN archaea encode a split version of RNApol A. In Asgard archaea, among the previously published genomes, only Heimdallarchaeote LC_3 (reclassified as Hodarchaeales) was found to encode a fused RNApol A, which was claimed to represent a contaminant[29]. Including a much broader taxonomic sampling of Asgard archaea diversity confirms a complex history of this protein involving multiple fusion and fission events (Figure S4.1.1_03): RNApol A of Thor-, Odin-, Sif, Hermod- and Jordarchaeia sequences are split, while RNApol A homologs in Baldrarchaeia are fused for two out of three representatives and nested within the formerly mentioned clade of sequences; the third baldrarchaeion (Baldrarchaeia_Yap30_bin4_67) encodes a split version. In addition, all Lokiarchaeales homologs are split, whereas in its sister group, Helarchaeales, RNApol A has the fused arrangement in one subclade, and the split version in the sister-clade. Interestingly, among Hodarchaeales, the hodarchaeal LC_3 RNApol A is not the only fused version anymore since Hodarchaeales_S146_22 also encodes that version. All other Hodarchaeales RNApol A homologs are split. However, phylogenetic analyses of each of the subunits/domain show that they branch with other Heimdallarchaeia homologs, showing that they do not represent contaminants (Supplementary Figures 29-30). Finally, all

Njordarchaeales also display the single gene configuration, whereas Gerdarchaeales, Kari- and Heimdallarchaeaceae encode a split version.

###### 4.1.2. Diphthamide/EF2

Diphthamide is a modified histidine residue which is uniquely present in archaeal and eukaryotic elongation factor 2 (EF-2)[40]. Because of the essential role of diphthamide in translational fidelity, it was long assumed that diphthamide biosynthesis genes (*dph*) were conserved across all eukaryotes and archaea. A recent study showed that some Asgard archaea and other archaea lack *dph*[40] and that many of these *dph*-lacking archaea encode a second EF-2 copy missing key residues required for diphthamide modification. The study found that some Heimdallarchaeia maintain *dph* genes and a single gene encoding a canonical EF-2. Here, we confirm that all Hodarchaeales members encode *dph* genes, while all other Heimdallarchaeia have lost this pathway. Moreover, Hodarchaeales encode a single gene for a canonical EF-2 with the diphthamide modification motif, which branch at the base of their eukaryotic counterparts in phylogenetic reconstructions (Supplementary Figure 11).

###### 4.1.3. zf-PARP

A critical DNA damage response signaling molecule in eukaryotes is the posttranslational modification poly(ADP-ribose) (PAR). This molecule is produced by a family of structurally and functionally diverse proteins called poly(ADP-ribose) polymerases (PARPs). The amino-terminal region of PARPs consists of two PARP-type zinc fingers. This region acts as a DNA nick sensor. We identified homologs of those PARP-specific zinc fingers (zf-PARP), which could only be detected in Heimdall- and Thorarchaeia.

###### 4.1.4. Histone N-terminal extensions

Another interesting observation relates to histones -- highly alkaline DNA-binding proteins responsible for packaging and compacting the DNA in all eukaryotes and in most archaea.

Histone proteins are composed of the universal histone fold, and in eukaryotes, of an additional tail. These tails contribute to tighter DNA packaging and can undergo post-translational modifications (e.g. acetylation, methylation, phosphorylation and ubiquitylation) as a way to regulate DNA compaction and thus gene expression, DNA repair, and many other processes[41]. N-terminal tails were until very recently thought to exist only in eukaryotic histones, and have been identified in one Heimdallarchaeial lineage[42], as well as in two other archaea (a Huberarchaeote and a Bathyarchaeote)[43]. Here we identified N-terminal tails in another 3 lineages of Heimdallarchaeia (all 3 belonging to Hodarchaeales), but also in 3 Njordarchaeales genomes. Those are all of roughly the same length and sequence composition as eukaryotic tails, and like in eukaryotes, the numerous lysines in the Asgard archaeal histone tails may well be subject to acetylation. In addition, Asgard archaeal genomes encode a large number of Gcn5-related N-acetyltransferases, to which some histone acetyltransferases belong[43]. This raises the possibility that the ability to control DNA compaction and gene expression through the modification of the N-terminal histone tail in eukaryotes was inherited from their Asgard archaeal ancestor.

#### 4.1.5. E2F/DP

Some Asgard archaea genomes were found to encode E2F/DP proteins, a large family of transcription factors that are key in the progression of the cell cycle in eukaryotes. This represents another gene family shared between Asgard archaea and eukaryotes but absent from other archaeal lineages. Its presence in some Asgard archaeal genomes was recently pinpointed[44], and we identified homologs spanning most groups of Asgard archaea (Figure 3). The DNA-binding domain present in the Asgard archaeal E2F/DP proteins is highly conserved and closely resembles the eukaryotic domain, in particular the residues involved in base-contacts, which suggests that these proteins recognise similar DNA-motifs as their eukaryotic counterparts. However, the homology of Asgard archaeal E2F/DP proteins is restricted to the

DNA-binding domain, and they lack binding domains to their regulators as well as the dimerization domain (most eukaryotic E2F and DP members form heterodimers with each other). Altogether, this suggests a different regulation mechanism and function of those proteins in Asgard archaea. Nevertheless, the eukaryotic E2F/DP proteins likely have arisen from multiple duplications of the homolog inherited from Asgard archaea combined with fusion events to additional domains leading to their complex integration to the cell cycle.

###### 4.1.6. PAC4

The assembly of 20S proteasomes requires dedicated proteasome assembly chaperones (PAC)[45]. Among those are two dimeric chaperone complexes composed of PAC1-PAC2 and PAC3-PAC4, respectively. Interestingly, we could detect well conserved PAC4 domains in most Asgard phyla.

##### 4.2. N-glycosylation (OST complex)

We expanded on previous reports and uncovered novel Asgard archaeal homologs of ESPs associated with N-glycosylation processes. In eukaryotic cells, the translocon is responsible for transporting proteins across or inserting them into the endoplasmic reticulum (ER) membrane. The translocon is formed at its core by the Sec61 protein-conducting channel and several accessory components, which assist Sec61 or facilitate protein maturation by covalent modifications and chaperone-like functions[46] (Figure 3). The translocon-associated protein (TRAP) complex is formed by two subunits in most eukaryotes (four in animals and fungi) and represents one of these translocon accessory components. It aids translocation of proteins and is thought to have an important role in the biogenesis of N-glycosylated proteins[47], alongside with the multimeric oligosaccharyltransferase (OST) complex, another major translocon component involved in N-glycosylation. The eukaryotic OST generally comprises 6-8 subunits that are collectively embedded in the membrane of the rough ER, and are organized into 3 subcomplexes. STT3/AglB belongs to subcomplex II and represents the catalytic subunit, and its homologs are found among the three domains of life. In contrast, other subunits do not possess prokaryotic homologs. Two exceptions were previously reported to possess homologs in all or some Asgard archaea[23]: OST1/Ribophorin 1 (subcomplex I) and OST3/Tusc3 (subcomplex II). Consistent with this, we found homologs of these two proteins in most of the new Asgard archaea clades. Additionally, here, we report Asgard archaeal homologs of two other subunits: OST5/TMEM258 and WBP1/Ost48. OST5 (subcomplex I) was found in all Asgard archaea phyla and in no other archaea. Most interestingly, we identified divergent WBP1 homologs in all Heimdallarchaeia clades, including Njordarchaeales (confirmed by reverse BLAST and HHblits), further supporting the phylogenetic position of Njordarchaeales as close relatives of Heimdallarchaeia and eukaryotes, and making this the first subcomplex III subunit described in Asgard archaea. These findings indicate that the Asgard archaeal ancestor of eukaryotes likely encoded at least 5 subunits belonging to all three subcomplexes defining the eukaryotic OST complex.

In addition, we identified divergent Asgard archaeal homologs of all four subunits of the TRAP complex. Homologs of the TRAP alpha are very divergent but could be detected through profile-profile cluster annotation in several Loki-, Hermod-, Thor- and Heimdallarchaeia, the latter two encoding two copies, one much more divergent than the other. In parallel, we have identified proteins encoded by most Asgard archaeal lineages and containing a clear TRAP beta domain (PFAM domain PF05753), although these proteins are usually much longer than their eukaryotic counterparts. Finally, TRAP delta was identified only in Thorarchaeia, and TRAP gamma domains were identified in a few proteins that are broadly distributed across Asgardarchaeota (Figure 3).

#### 4.3. Vesicular trafficking and membrane remodelling

Vesicular transport is an essential process in eukaryotic cells and its emergence was key to eukaryogenesis. All eukaryotes have a set of vesicle coat proteins, which couple cargo selection to vesicle budding in the secretory and endocytic pathways[48]. Previously reconstructed Asgard archaeal genomes encoded predicted protein domains that in eukaryotes are associated with intracellular trafficking and secretion[23,24]. These include ESCRT (endosomal sorting complexes required for transport), TRAPP (transport protein particle), and homologs of the Sec23/24 COPII (coat protein complex II) vesicle coatomer protein complex, together with many small GTPases that are closely related to Rabs (i.e., important for transport-vesicle budding, motility, docking and fusion in eukaryotes). Below we describe further investigations of newly uncovered homologs of ESPs involved in membrane trafficking.

##### 4.3.1. Adaptor protein complexes

A key to the emergence of eukaryotes was the ability to regulate trafficking pathways between their intracellular membrane compartments. This is done in large part by the heterotetrameric adaptor complexes, AP1 to 5, and hexatetrameric TSET and COPI complexes[49]. Their main function is to select cargo for packaging into transport vesicles. In combination with membrane-deforming proteins such as clathrin and the COPI B-subcomplex, they facilitate protein and lipid trafficking between compartments in the secretory and endocytic pathways. All 7 complexes share a similar and homologous architecture, due to their emergence through duplication events that took place before the last eukaryotic common ancestor (LECA)[49]. Their core is composed of two large subunits (~100 kD), the β and γ families, a medium subunit μ (~50 kD), and a small subunit σ of ~20 kD.

The β and γ subunits both contain an Adaptin_N (PFAM PF01602) domain at the N-terminus, and a specific C-terminal domain, B2-adapt-app_C (PF09066) and Alpha_adaptinC2 (PF02883), respectively (Extended Data Figure 2). While Adaptin_N could be identified in many Asgard lineages, we could not detect either B2-adapt-app_C or Alpha_adaptinC2 right downstream of it. However, we identified standalone versions of B2-adapt-app_C and Alpha_adaptinC2 domains in sparsely distributed Asgard archaea (Figure 3).

The μ and σ subunits are composed of a Clat_adaptor_s (PF01217), which is fused to a Adap_comp_sub (PF00928) domain in the former, and is standalone in the latter. We identified both of those domains in most Asgard archaeal phyla, although with a sparse distribution across representatives, due to the high divergence to their eukaryotic counterparts. While we could identify both of those domains, we did not clearly see individual proteins displaying both domains. It is worth noting that Clat_adaptor_s was sometimes found to be fused to other ESPs involved in membrane-trafficking in eukaryotes, such as the Arf GTPase, Vacuolar fusion protein Mon1 and Roadblock/LC7 domains.

In summary, we detected Asgard homologs of all 5 domains constitutive of AP subunits. Although their arrangement is different from the one found in eukaryotic AP proteins, this suggests that eukaryotes inherited the building blocks of these key protein complexes from their Asgard archaeal ancestor and that a similar complex involved in membrane deformation processes could exist in some Asgard archaea.

###### 4.3.2. Yip1

We uncovered homologs of Yip1 proteins in all Asgard archaeal orders. In eukaryotes, Yip1 domain family (YIPF) proteins are multi-span, transmembrane proteins mainly localized in the Golgi apparatus[50]. YIPF proteins have been found in virtually all eukaryotes, suggesting that they have essential functions. Early analyses in *Saccharomyces cerevisiae* indicated that Yip1 plays a role in budding of transport vesicles and/or fusion of vesicles to target membranes, and it is required for transport between the endoplasmic reticulum and the Golgi[51]. Surprisingly, we found that the PFAM domain associated with this protein family (Yip1, PF04893) is commonly found in the three domains of life but no function has been reported for prokaryotic family members, and the significance of their similarity to the eukaryotic family members requires further investigation. Interestingly, investigations of genomic context indicate that *yip1* genes are often located in close vicinity to other genes encoding proteins involved in vesicle trafficking and membrane remodelling, such as ESCRT (I, II and III) and ubiquitin modifier system components. For example, they are flanking ESCRT-I subunit homologs in almost all Lokiarchaeia genomes. They are also often flanking genes encoding predicted proteins containing transmembrane domains (e.g., DUF2208, DUF5518, DUF1097). It is therefore tempting to speculate that Asgard archaeal Yip1 homologs have a role in vesicle biogenesis and/or trafficking pathways.

###### 4.3.3. HOOK domain

We identified a HOOK protein coiled-coil region found in representatives from several Asgard archaea phyla. In eukaryotes, the HOOK family of activating adaptors is one of the most conserved families of dynein adaptors. They take part in the 'FHF' complex (FTS, Hook, and FHIP (FHF complex subunit Hook Interacting Protein))[52]. Hook proteins contain a highly conserved N-terminal domain (mediating attachment to microtubules), and a more divergent C-terminal domains involved in binding to specific organelles. Additionally, a coiled-coil motif (PF05622) serves for homodimerization. We could only detect this central coiled-coil domain in Asgard archaea, but neither the N- or C-terminal domains. This is thus in itself only weak evidence for a role in membrane trafficking. However, it is interesting to observe that, in 25 homologs (9 Lokiarchaeales, 5 Thorarchaeia, 1 Hermodarchaeia, 9 Gerdarchaeales, 1 Njordarchaeales), this domain is found neighbouring an ATG16 domain (PF08614), which is itself a eukaryote-specific protein domain involved in the autophagy pathway[53]. The sequence divergence was however too high to align them accurately to their putative eukaryotic homologs.

##### 4.4. Endosomal sorting

Below we describe the identification of several new ESPs represented by Vacuolar protein sorting-associated protein (Vps). These are part of the Endosomal Sorting Complex Required for Transport (ESCRT) system, which performs the topologically unique membrane bending and scission reaction away from the cytoplasm. Many of these newly identified Vps homologs show a punctuated distribution across Asgard archaea. Although this could represent the true distribution, we suspect that they are here under-reported, first because they are small proteins, and second, because we took the conservative approach to only report cases in which their corresponding PFAM domain represents the best PFAM hit for a given protein.

###### 4.4.1. Retromer complex proteins Vps5, Vps26, Vps29 and Vps35

A particularly novel aspect reported here is the presence of homologs of the majority of proteins described as being involved in the retromer complex (Extended Data Figure 2). Retromer is a coat-like complex associated with endosome (or lysosome)-to-Golgi retrograde traffic[54]. It is formed by Vacuolar protein sorting-associated protein 35, Vps5, Vps17, Vps26 and Vps29[55]. These actually compose two subcomplexes: the cargo-selective complex (CSC), made of Vps26, Vps29, and Vps35[56]; and the sorting nexin (SNX) and Rvs (BAR) dimer, consisting of Vps17 and Vps5. During cargo recycling, retromer is recruited to the endosomal membrane via the Vps5-Vps17 dimer. Cargo recognition is thought to be mediated primarily through Vps26 and possibly by Vps35. Finally, the BAR domains of Vps5-Vps17 have the ability to sense and induce membrane curvature, are involved in various processes including endosome-to-Golgi retrograde trafficking and endocytosis[54]. Their distribution is sparse, but we have detected

Asgard archaeal homologs of all subunits except for Vps17 (Figure 3, Extended Data Figure 2). Interestingly, the Thorarchaeota Vps5-BAR domain is often fused to Vps28, a subunit of the ESCRT machinery complex I, suggesting a functional link between BAR domain proteins and the thorarchaeial ESCRT complex. This is, to our knowledge, a domain architecture that is not found in eukaryotic proteins.

##### 4.4.2. Vps10-Sortilin, a retromer cargo

The best-characterized retromer cargo is yeast Vps10, a member of the Sortilin receptor family. This transmembrane protein receptor is known in yeast and mammal cells to be involved in the sorting and transport of lipoproteins between the Golgi and the endosome. The Vps10 receptor releases its cargo to the endosome and is recycled back to the Golgi via the retromer complex[57]. We detected Vps10 domains in most Heimdallarchaeia orders, as well as in Loki- and Helarchaeales and in Hermod- and Thorarchaeia.

##### 4.4.3. Vps62

The function of Vps62 in eukaryotes has not attracted a lot of scrutiny but it has been shown to be located in vacuoles and to be required for protein targeting to the vacuole[58]. Interestingly, we have identified Vps62 homologs in all Thorarchaeia representatives as well as a handful of divergent homologs in Asgardarchaeia, Helarchaeales and in Kariarchaeaceae.

##### 4.4.4. CORVET/HOPS complex proteins

Endosomal fusion and autophagy depend on the class C core vacuole/endosome tethering (CORVET) and homotypic fusion and protein sorting (HOPS) that are hexameric complexes[59]; they share the class C core consisting of the subunits Vps11, Vps16, Vps18, and Vps33[60]. In addition, HOPS is composed of Vps41 and Vps39)[61]. Vps39, found associated to late endosomes and lysosomes, promotes endosomes/lysosomes clustering and their fusion with autophagosomes[62]. We could identify a few homologs of the Vps11, Vps16, Vps18 and Vps39 proteins in Asgard archaea.

###### 4.4.5. Vps4 regulators: Vfa1, Vta1 and Ist1

Vfa1, in yeast, is an endosomal protein that interacts with the ATPase Vps4 and is involved in regulating the trafficking of other proteins to the endocytic vacuole[63]. It is involved in the transport of biosynthetic membrane proteins from the prevacuolar/endosomal compartment to the vacuole and is required for multivesicular body (MVB) protein sorting[64]. Vfa1 also catalyzes the ATP-dependent dissociation of class E VPS proteins from endosomal membranes, such as the disassembly of the ESCRT-III complex[64].

Vta1 is another positive regulator of Vps4 ATPase through the promotion of correct assembly of Vps4 and stimulation of its ATPase activity[65].

Finally, Ist1 appears to regulate the recruitment and oligomerisation of Vps4, thereby regulating the flow of cargo through the MVB pathway[66].

We identified Vfa1 protein domain as the best hit in Gerd- and Lokiarchaeales, as well as in Thorarchaeia proteins; Vta1 was detected in Njordarchaeales and Hodarchaeales, in Lokiarchaeales and Thorarchaeia; Ist1 was the best domain detected in a handful of proteins from Hodarchaeales and Kariarchaeaceae, Lokiarchaeales and Thorarchaeia.

###### 4.4.6. GARP complex subunits Vps51, Vps52, Vps53 and Vps54

The Golgi-associated retrograde protein (GARP) complex is a multisubunit tethering complex located at the trans-Golgi network where it functions to tether retrograde transport vesicles derived from endosomes [67,68]. GARP comprises four subunits named vacuolar protein sorting 51 (VPS51), VPS52, VPS53, and VPS54. We detected all of these domains as best hits in several lineages across Asgard archaea.

##### 4.4.7. Vps55/68 Sorting Complex

Genome-wide screens in yeast have shown that Vps68 localizes to endosomes and that it forms a complex with Vps55[69]. Very recently, it has been demonstrated that Vps68 physically interacts with ESCRT-III and that it cooperates with ESCRT-III in intraluminal vesicles formation at late endosomes [70]. Interestingly, we uncovered homologs of these two proteins in a few Asgard lineages.

##### 4.4.8. ESCRT-III accessory protein Bro1

Bro1 proteins are involved in (1) cargo recognition in concert with or in parallel to the early ESCRTs, (2) regulating ESCRT-III dynamics by facilitating Snf7 activation and inhibiting Vps4 disassembly of ESCRT-III, and (iii) ESCRT-dependent MVB biogenesis in vivo by facilitating ILV formation[71]. These diverse contributions suggest Bro1 domain family members may serve roles coordinating cargo entry into budding ILVs during MVB sorting. The Bro1 domain is the best domain detected in Hod- and Njordarchaeales, as well as in Thorarchaeia.

In conclusion, despite the somewhat patchy distribution, the repertoire of Asgard homologs of proteins involved in intracellular membrane trafficking in eukaryotes is even vaster than previously thought[17,23,24] and broadly distributed. Given the predicted functional cohesiveness of all these components, it is tempting to speculate on the existence of a form of intracellular trafficking in Asgard archaea which would be possibly supported by an actin-based cytoskeleton, similar to the one observed in Lokiarchaeia[33,72].

#### 5. Ancestral genome reconstruction

To study the evolutionary dynamics of gene families and to estimate the gene content of ancestral genomes we used Amalgamated likelihood estimation (ALE), a probabilistic gene-tree aware method. This approach uses information from species- and gene-trees to distinguish vertical inheritance (tree-like) from horizontal transfers through a process called reconciliation, which aims to fit a gene tree into a species tree. The discord between topologies is used to infer the duplication, transfer and loss (DTL) events in each gene family. Additionally, by using gene-tree distributions (e.g., based on bootstrap trees or posterior distribution), rather than a single gene-topology (that would correspond to the maximum likelihood or consensus tree), ALE takes into account the uncertainty associated with phylogenetic reconstructions of individual gene families that, e.g. due to their short length, might lack enough phylogenetic signal to confidently establish their evolutionary history. As a result, ALE reduces the impact that poorly supported bipartitions present in individual gene trees may have in the estimation of DTL events. Furthermore, this approach also accounts for the fact that numerous species are extinct or not represented in the dataset, and allows for horizontal transfer events involving unrepresented lineages.

##### 5.1. General considerations about the gene tree/species tree reconciliation approach used here

Ancestral reconstruction approaches have been developed to be used in conjunction with complete genomes and the impact of including incomplete MAGs in the DTL inferences still needs to be evaluated. Notwithstanding, given that complete genomes are not available for numerous archaeal lineages, in particular from any Asgard archaeal member at the time that these analyses were first performed, the use of MAGs becomes inevitable to study the evolution of such groups. We reasoned that, if MAGs are incomplete, the biggest effect will be observed at the terminal nodes in which genome incompleteness will be incorrectly regarded as gene losses, and, hence, the number of losses will be overestimated. If the dataset contains several related MAGs, and there is no systematic bias in the genes that are lacking, the effect of including those incomplete genomes in the analyses will be minimal at internal nodes. In addition, contamination present in MAGs could also affect the DTL modeling by incorrectly regarding genes that are the result of erroneous binning as if they were gene gains (lateral gene transfers or de novo originations), or duplications. Similarly, we argue that if the contaminated sequences present in MAGs are random, such artefacts will mostly affect the terminal nodes. Given that the present analyses focus on the reconstruction of the gene content in ancestral nodes, the above-mentioned artefacts are unlikely to significantly impact our results.

To assess how the inclusion of MAGs might affect the DTL modeling, we thus examined the number of DTL events predicted in internal and terminal nodes (Supplementary Figure 31). We observed a higher number of losses predicted at terminal nodes, but no increased numbers of gene gains or duplications. This observation indicates that most biases in the DTL analyses originate from incomplete, rather than contaminated, genomes. This is in agreement with the estimated quality of the Asgard archaeal MAGs used in the present study, which are often incomplete but show, in general, low levels of contamination.

Furthermore, the reported biases are further supported by the correlations observed between the number of events predicted and the completeness and contamination values estimated for the terminal nodes (Supplementary Figure 31). Taking into consideration these observations, and to reduce any impact of including incomplete genomes on the number of inferred events, we excluded the terminal nodes (i.e., with a potentially overestimated number of losses) when inspecting the evolutionary dynamics of genomes of Asgard archaea and other archaea.

Yet, the use of MAGs is expected to affect some of the reconciliations performed and, therefore, the accuracy of the inferences. Properly controlled simulation analyses are needed to fully understand expected biases and identify problematic cases, which is beyond the scope of the current analyses. Of note, several recent studies have successfully implemented the analysis of MAGs using gene tree/species tree reconciliation approaches to infer ancestral gene content in prokaryotes[73–75].

#### 6. Genome content of the last archaeal common ancestor of the eukaryotes

##### 6.1.1. The last archaeal ancestor of eukaryotes probably had a developed cytoskeleton but different from *Lokiarchaeum*

Based on the reported cellular morphology of two closely related Lokiarchaeales lineages, *Ca.* P. syntrophicum[72] and *Ca.* L. ossiferum[33], scenarios for the origin of the eukaryotic cell have been suggested in which the archaeal ancestor could form cellular protrusions and membrane vesicles[72]. In this hypothesis, these protrusions are predicted to have 'entangled' and facilitated the engulfment of the proto-mitochondrial symbiont by the archaeal host. Yet, it is unlikely that the morphological features of *Lokiarchaeum* resemble the ancestral state of the last Asgard ancestor of eukaryotes. In fact, although the ancestor of Asgard archaea is inferred to harbour homologs of ESPs such as actin, profilin, proteins of the villin/gelsolin superfamily and ARP2/3 components, the number of copies predicted shows a wide variation among the ancestors of the various Asgard archaeal phyla. Specifically, the results of the ancestral reconstruction indicate that the profilin and villin/gelsolin families experienced numerous duplication events in several of the Asgard archaeal phyla and that the ARP2/3 proteins were frequently lost. In particular, we observe elevated levels of duplication of genes encoding cytoskeletal proteins in Lokiarchaeia. Additionally, the ancestors of all Heimdallarchaeia including Hodarchaeales are predicted to encode for another actin-related protein, a putative homolog of MreB, which is absent in Lokiarchaeia. More generally, our inferences suggest that, during the evolution of Lokiarchaeia, there were numerous duplication events that resulted in the expansion of several gene families with function associated with the cytoskeleton dynamics and the trafficking machinery. Despite the fact that the Hodarchaeales ancestor was inferred to contain most of these gene families, these were usually in fewer copies. As a result, we expect fundamental differences in their cytoskeletal abilities and morphologies. Therefore, we raise a note of caution regarding considering the features observed in present-day organisms as ancestral, especially when based on the study of very few closely related lineages. While having access to cultured representatives of Asgard archaea lineages is extremely valuable for increasing our understanding of their cell biology, physiology and metabolism, inferences of ancestral characteristics should be done within an evolutionary framework. Insights obtained from culturing diversified Asgard archaeal lineages, in conjunction with gene-content inferences of various Asgard archaea ancestors, will prove indispensable to determine the nature of the last Asgard archaeal ancestor of eukaryotes.

###### 6.1.2. Metabolic features of Asgard archaeal ancestors

We used the inference of ancestral gene presence to investigate the metabolic potential of various Asgard archaea ancestors. In particular, we focused on the last common ancestor of all Asgard archaea, the last common ancestor of Heimdallarchaeia, and the ancestor of Hodarchaeales.

**Gluconeogenesis and glycolysis**. Across the tree of life, organisms utilize the Embden-Meyerhof-Parnas (EMP) and Entner-Doudoroff (ED) pathways to metabolize glucose. The ED pathway is the prominent glycolytic pathway in bacteria and is rarely found in archaea and eukaryotes. The main difference of the ED pathway compared to the EMP pathway lies in the early-stages of the reaction sequence yielding glyceraldehyde-3-phosphate (GAP, Supplementary Figure 14). In our ancestral state reconstruction analysis, we predict that the Asgard archaea ancestors encoded the majority of the EMP and ED pathways (Supplementary Figure 15). The Asgard ancestor likely lacked the EMP pathway-specific enzyme ATP-dependent phosphofructokinase (ATP-PFK; COG0205/arCOG03641; broadly distributed in eukaryotes and Bacteria). However, the ancestor of both Heimdallarchaeia and Hodarchaeales likely encoded a ATP-PFK, suggesting a complete sugar degradation pathway in these ancestors (Supplementary Figure 15). The majority of the ED pathway was predicted to be present in the Asgard archaea ancestors, however, we suspect this pathway was not glycolytic owing to the absence of some components, namely 2-Keto-3-Deoxy-(6-Phospho) gluconate aldolase (KDPGA). KDPGA produces pyruvate and glyceraldehyde 3-phosphate (GAP) (Supplementary Figure 14), which is present in all archaeal ED pathway utilizers, is not present in any of the Asgard archaeal ancestors. This suggests that the early reactions of the ED pathway might have been important for 6-phosphogluconate production for the pentose phosphate pathway and not glycolysis (Supplementary Figure 14).

We predict that the Asgard archaeal ancestors had at least two possible ways to generate 6-carbon sugars: i) from phosphoenolpyruvate (PEP) via the reverse EMP gluconeogenic pathway, necessary for the formation of fructose 1,6-bisphosphate (FBP), and ii) via a reversed EMP pathway through the conversion of glyceraldehyde 3-phosphate to fructose 6-phosphate by FBP aldolase/phosphatase (FBP A/P). FBP A/P is thought to be an ancestral gluconeogenic enzyme that has a restricted distribution in archaea and some bacteria[76]. We predict that FBP A/P was present in the Asgard archaeal ancestor and subsequently lost in the Heimdallarchaeia and Hodarchaeales ancestors. Since the Asgard ancestor is predicted to encode an FBP A/P (an ancestrally gluconeogenic enzyme[76]), we hypothesize it was able to make sugars. In contrast, we failed to identify the FBP A/P enzyme in either the Heimdallarchaeia or Hodarchaeales ancestors and therefore hypothesize these ancestors did not perform FBP A/P-mediated gluconeogenesis. Whether these ancestors used the EMP pathway for glycolytic or gluconeogenic purposes cannot be determined. Indeed, the EMP pathway could have been co-opted for glycolytic purposes following a shift toward heterotrophic growth as has been suggested in other systems[76]. For example, the change in global sugar supply may have relaxed selection for the maintenance of the FBP A/P ancestral unidirectional enzyme in heterotrophic bacteria and eukaryotes[76].

**Reducing power formation via the oxidative pentose phosphate pathway.** The pentose phosphate pathway (PPP) is widespread in bacteria and eukaryotes and provides reducing equivalents (NADPH) for reductive biosynthesis (via the oxidative pentose phosphate pathway, OPPP) and precursors for the biosynthesis of nucleotides and aromatic amino acids (via the non-oxidative pentose phosphate pathway, NOPPP). Although the OPPP is rare in Archaea[77], our results indicate the presence of a partial OPPP in the Asgard and Hodarchaeales ancestors (Supplementary Figure 14). Interestingly, our analyses indicate that the OPPP enzyme 6-phosphogluconolactonase (6GPL) mainly found in Bacteria and eukaryotes (and recently identified in the archaeon *Haloferax volcanii*)[78,79], was present in Heimdallarchaeia and Hodarchaeales ancestors and was lost in the rest of the lineages except the Helarchaeales ancestor (Supplementary Figure 15, Supplementary Table 4). These results suggest that Hodarchaeales and eukaryotes have analogous central carbon pathways (EMP and OPPP), although the phylogenetic ancestry of these pathways requires further investigation.

**Nucleotides biosynthesis via the reverse ribulose monophosphate pathway.** Ribulose-5-phosphate (Ru5P) is a key precursor for nucleotide biosynthesis and can be synthesized by the OPPP, NOPPP and the ribulose monophosphate pathway (RuMP) operating in the reverse reaction. We failed to identify two key components of the NOPPP (i.e., transketolase TK and transaldolase TA) in any extant Asgard archaeal genomes, suggesting this pathway is not used and likely absent in the ancestors. However, we did identify components of the RuMP pathway, the 3-hexulose-6-phosphate synthase (HPS, COG0269) and 3-hexulose 6-phosphate (PHI, COG0794) that converts the reversible interconversion of Ru5P to F6B respectively. The presence of key enzymes for nucleotide biosynthesis via R5P (phosphoribosylpyrophosphate (PRPP) synthase (COG0462); orotate phosphoribosyltransferase (COG0461), and glutamine phosphoribosyl amidotransferase (COG0034)) in all Asgard archaeal ancestors, suggest that this pathway could be operating in reverse direction (rRuMP) as a possible alternate means of producing Ru5P from F6P derived from glycolysis or gluconeogenesis.

**Aromatic amino acid biosynthesis.** The Asgard archaeal ancestors likely lacked key genes for the production of erythrose 4-phosphate (E4P), an important precursor for chorismate, and ultimately aromatic amino acid (AroAA), biosynthesis. However, we did uncover a pathway for producing chorismate via a partial shikimate pathway that does not require E4P. This alternative route has been suggested for other anaerobic archaea via *de novo* synthesis of F1,6BP derived from FBPA A/P[80]. Under this scenario, FBPA A/P could have played an important role in providing the precursor for the alternative aromatic biosynthesis pathway, indicating a crucial role in the anabolic metabolism of the Asgard archaeal ancestors. Other pathways for aromatic amino acid biosynthesis have been suggested[80], for example the incorporation of exogenous aryl acids via indolepyruvate oxidoreductase (IOR, arCOG01609); however this enzyme was not found to be present in any Asgard archaeal ancestors.

**Acetate utilization.** Our ancestral reconstructions predict that all Asgard archaeal ancestors encoded an ADP-dependent, ADP-forming acetyl-CoA synthetase (ACDs, arCOG01340). This enzyme can convert acetyl-CoA into acetate with the concomitant production of ATP by substrate-level phosphorylation (Supplementary Figure 14, “Acetyl-CoA to Acetate”). This is a major energy-conserving enzyme for sugar and peptide fermentation in hyperthermophilic archaea, and is considered to be an ancient strategy for ATP synthesis[81]. In the absence of other substrates, acetate can also serve as a source of carbon via the conversion to acetyl-CoA, in all the Asgard archaeal ancestors given the presence of the (ACS) (arCOG06112).

**The tricarboxylic acid cycle as a source of reducing power.** In agreement with previous results[26], all Asgard archaeal ancestors appear to have encoded a complete tricarboxylic acid (TCA) cycle, and lacked ATP citrate lyase for the reverse TCA cycle. One of the enzymes of the TCA cycle, isocitrate dehydrogenase (IDH, ENOG41122A1) can play an important role in the generation of reducing equivalence for the ETC (i.e., NADH) but also for biosynthetic processes (i.e., NADPH)[82]. In eukaryotes, IDH exists in two forms: an allosteric $NAD^+$-linked IDH found only in mitochondria, and a non-allosteric $NADP^+$-linked IDH that is found in both mitochondria and cytoplasm. Some archaea also have IDH proteins with different preferences (e.g., arCOG01163 and arCOG01164 for NAD+ and NADP+, respectively). At present, we cannot distinguish between the specificity ($NAD^+$ or $NADP^+$) of the IDH of the Asgard archaeal ancestors based solely on our orthology assignment. However, studies have suggested that organisms that encode isocitrate lyase (COG2513) always encode an $NADP^+$-dependent IDH[83]. The only Asgard ancestor that encodes isocitrate lyase and IDH is the Hodarchaeales, suggesting that, like some eukaryotes, the Hodarchaeales IDH might be able to use NADP+ as a substrate. This would provide an important source of reducing power when growing on acetate (which could enter the TCA following conversion to acetyl-CoA).

**RuBisCO utilization and nucleoside assimilation.** In agreement with previous studies[84], each Asgard archaeal ancestors likely encoded proteins belonging to the ribulose 1,5-bisphosphate carboxylase (RuBisCO) family (COG1850). Based on previous phylogenetic analyses[26], the RuBisCO encoded in extant Asgard archaeal is affiliated with type IV or archaeal type III, thus is likely important for the salvaging and assimilation of nucleosides and not carbon fixation. Our results indicate that most of the genes encoding proteins important for adenosine monophosphate (AMP) salvage were present in the Asgard archaeal ancestors (Supplementary Table 5), supporting assimilatory-type RuBisCO for the generation of 3-phosphoglycerate from AMP that can enter the EMP pathway to produce acetyl-CoA (assimilating salvaged nucleotides into carbon central metabolic pathways). Interestingly, two enzymes belonging to the AMP salvage pathway show a clear distinction in their copy numbers in the Asgard archaeal ancestors. Adenylate kinase (COG0563) and phosphoribulokinase/uridine kinase (PRK/UK) family (IPR006083, arCOG05133) were predicted to be present in the Heimdallarchaeia and Hodarchaeales ancestors (also likely present in the ancestor of Heimdall-Njordarchaeales) but lost in the other ancestors (except for Loki- ancestor). Although the PRK/UK homologs that have been detected in extant Heimdallarchaeia are thought to be UKs based on domain composition, we cannot rule out PRK activity in the Heimdallarchaeia and Hodarchaeales ancestors, given the many enzymatic links (pentose bisphosphate pathway, glycolysis, gluconeogenesis, and amino acid metabolism) present in the Asgard archaeal ancestors (Supplementary Figure 14). Our results are in agreement with previous studies[85] where it has been suggested that the photosynthetic Calvin-Benson-Bassham pathway originated from a primitive carbon metabolism utilizing RuBisCO, like the reductive hexulose-phosphate pathway (RHP), that was potentially operational in Heimdallarchaeia and Hodarchaeales ancestors.

**Hydrogen metabolism present in the Asgard ancestors.** Apart from group 3b and 3c [NiFe]-hydrogenases, various membrane-bound group 4 [NiFe]-hydrogenases have previously been identified in Odin- and Heimdallarchaeia genomes[26]. Using this expanded taxon sampling of Asgard archaea, we could also identify homologs in several other Asgard archaeal phyla (Njordarchaeales, Jord-, Baldr- and several Thorarchaeia lineages), indicating that group 4 [NiFe]-hydrogenases were present in the Asgard archaeal ancestors (Supplementary Table 4). Although it was not possible to specifically establish the ancestral state of [NiFe]-hydrogenases of the group 3b and 3c using ALE, because they belong to the same gene family as other [NiFe]-hydrogenases, manual inspection of their phyletic pattern suggests that these two types were already present in the Asgard archaeal ancestor (Supplementary Table 4). Further analyses are needed to predict whether these homologs participated in hydrogen evolution or consumption (diversity of [NiFe] hydrogenases are reviewed elsewhere[86].

**The Wood-Ljungdahl but not the methyl-CoM reductase pathway was present in the Asgard archaeal ancestor.** Our ancestral reconstructions predicted that the last common Asgard archaeal ancestor encoded most components of the Wood-Ljungdahl pathway (WLP) including carbon monoxide dehydrogenase/acetyl-CoA synthase (CODH/ACS) and the formylmethanofuran dehydrogenase (*fmdABCDE*) (Supplementary Figure 14, Supplementary Table 4). This is in agreement with recent studies[26] that suggest that the Asgard archaeal ancestor was able to use $H_2$ lithoautotrophically from reduced compounds, or produce $H_2$ as a by-product of fermentation of molecules with higher oxidation states. Interestingly, our ancestral modelling suggests that the WLP was lost before the Heimdallarchaeia-Njordarchaeales ancestor; thus, it was also missing in LAECA (Supplementary Figure 15). The Helarchaeales ancestor appears to be the only Asgard archaeal ancestor which encoded a methyl-CoM reductase (MCR), which is likely involved in anaerobic hydrocarbon degradation[25].

**Biomass and energy conservation from formate, trimethylamine, and formaldehyde.** All Asgard archaeal ancestors were also predicted to encode aldehyde ferredoxin oxidoreductase (AFOR) for assimilation of several aldehydes (*i.e.,* crotonaldehyde, acetaldehyde, formaldehyde and glyceraldehyde), into central carbon pathways. Furthermore, most Asgard archaeal ancestors (except for Odinarchaeia and Helarchaeales) encode a trimethylamine methyltransferase (MttB) for the utilization of methylated compounds and methylamines[87]. A recent study[101] showed that this gene has a broad distribution across the tree of life including a newly-discovered non-methanogenic Archaea, Brockarchaeota, that is likely assimilating trimethylamine (TMA) via acetate production. Similar to Brockarchaeota, the Asgard archaeal ancestors lack methanogenesis-related genes including MCR, and methylamine-specific corrinoid protein MtbA (which transfers the methyl group to coenzyme M in methanogens or tetrahydrofolate $H_4F$ in acetogens). Furthermore, the Asgard archaeal ancestors encode components for the assimilation of TMA and formate into central metabolism. More specifically, all ancestors were predicted to contain the B12-binding corrinoid protein (COG5012), the glycine cleavage system (GCS) for the production or degradation of glycine, a serine-dehydratase-like enzyme (for conversion of the serine to pyruvate), and key enzymes of the $H_4F$ methyl branch of the WLP (such as the Methylene-$H_4F$ reductase and MTHFR). The presence of these pathways suggests the ability to assimilate methylated and single-carbon compounds was present in the Asgard archaeal ancestors. Overall, this suggests that the ancestors (except the Njordarchaeales ancestor) had the potential to use TMA and that genes for formate utilization were likely to have been present in Heimdallarchaeia and Hodarchaeales ancestors (Supplementary Figure 14).

**Operational oxidative phosphorylation in the Hodarchaeales ancestor.** The bioenergetic potential of the host lineage prior to the acquisition of the alphaproteobacterial endosymbiont has been a major gap in most eukaryogenesis models. In our ancestral state reconstructions, we predict that the ancestor of Hodarchaeales used nitrate as a terminal electron acceptor for energy conservation via nitrate respiration. The ability to respire with nitrate or oxygen would have provided metabolic flexibility based on the availability of terminal electron acceptors for the archaeal host. The subsequent evolution of the respiratory chain after the endosymbiotic event may have been a response of the organism to enhance its oxidative capabilities. Under this scenario, the Hodarchaeales ancestor likely produced ATP coupled to electron transport ultimately terminating in the reduction of nitrate.

Previous studies have shown that some Heimdallarchaeia, which lack the WLP, encode several components of the electron transport chain (ETC) that may support anaerobic and aerobic respiration[88]. We found that Heimdallarchaeia and Hodarchaeales ancestors likely encoded nitrate reductases and all the components of the ETC (except for complex III). Below, we describe our findings regarding each component of the ETC found in the Asgard archaeal ancestors.

**Complex I.** The proton-pumping NADH:ubiquinone oxidoreductase (Complex I, CI), is the first of the respiratory complexes generating the proton motive force essential for ATP production. homologs of this complex have been found in bacteria, archaea, mitochondria, and chloroplasts[89]. This complex can transfer electrons from NADH (Bacteria and mitochondria) or $F_{420}H_2$ (Archaea) to a quinone electron carrier (e.g., ubiquinone) to generate quinol with the concomitant transfer of protons across the membrane. These complexes can be distinguished based on the presence of the NADH-interacting module (NuoEFG and accessory subunit NuoA) or $F_{420}H_2$-interacting module (FpoF). The Asgard archaeal ancestors (specifically

Heimdallarchaeia, including Hodarchaeales) likely encoded most of the Complex I subunits (NuoA-K) including NADH interacting subunits (NuoAEFG). This suggests these ancestors could have used electrons from NADH analogous to the bacterial NADH:quinone oxidoreductase-like complex to ultimately generate a proton gradient. In contrast, in the Asgard archaeal ancestor, the NADH-interacting module (NuoAEFG) is incomplete. Therefore, the absence of a complete $F_{420}H_2$ or NADH-interacting module in the Asgard archaeal and Njordarchaeales ancestors could indicate that these organisms might have used a membrane-bound [NiFe] hydrogenase composed of NuoBCDHIL subunits. This would allow for the translocation of protons via hydrogen oxidation and not NADH oxidation and has been previously reported in other archaea[90].

**Complex II.** The second component of the ETC is succinate dehydrogenase (SDH). This complex does not contribute to the proton motive force directly but contributes electrons to the quinol pool. The complex is composed of two soluble subunits (SdhA and SdhB) and two membrane anchors (SdhC and SdhD) that funnel electrons from SdhB to the membrane-associated quinone species. We identified SdhA and SdhB in all Asgard archaeal ancestors. However, SdhC and SdhD were only detected in the Heimdallarchaeia and Hodarchaeales ancestors. Whether this is owing to the divergent nature of these subunits or their genuine absence cannot be determined with present data. The presence of SdhA and SdhB and not SdhC and SdhD in the Asgard archaea ancestor, could suggest these ancestors possessed a non-membrane bound SDH complex likely playing a role in the TCA cycle and the transfer of electrons through Fe-S clusters to an unknown electron carrier molecule in the cytoplasm. Our data suggest the Heimdallarchaeia and Hodarchaeales ancestors likely encoded SdhC and SdhD that could transfer electrons to a quinone such as menaquinone. In agreement with this, several genes coding menaquinone biosynthesis (3-polyprenyl-4-hydroxybenzoate decarboxylase,

COG0043; alpha/beta superfamily hydrolase, COG1506; multimeric flavodoxin WrbA, COG0655) were predicted to be present in Heimdallarchaeia and Hodarchaeales ancestors (Supplementary Table 4). Furthermore, in extant Asgard archaeal genomes, the set of genes involved in ubiquinone/menaquinone biosynthesis are near complete in Heimdallarchaeia, suggesting quinone species are components of their electron chain[26].

**Nitrate reductase and Complex III.** Previous investigations of extant Asgard genomes uncovered a putative operon of a cytoplasmic-facing nitrate reductase (NR), a cupredoxin-like protein and putative nitrate transporter (NarK)[26]. This complex is predicted to transfer electrons from a quinol to nitrate. Our ancestral state reconstructions suggest that in addition to the cupredoxin-like proteins the Heimdallarchaeial ancestor encoded the NarGHJ subunits while the Hodarchaeales ancestor likely encoded the NarGHIJ subunits. We previously identified a NarK/NarU-like nitrate transporter in some Heimdallarchaeia (one in the Hodarchaeales and the Kariarchaeaceae members LC_3 and LC_2, respectively)[26] and here we additionally identified a homolog in the kariarchaeaceae S139. These putative NarK/NarU-like sequences were clustered into a large cluster of 3481 major facilitator superfamily proteins (MFS), which was too large to be analyzed with ALE (see methods). However, manual investigations show that despite the sparse presence of these transporters, the sequences form a monophyletic group in phylogenies, suggesting that they were present in the last common ancestor of Hodarchaeales and Kariarchaeaceae, i.e. in the ancestor of Heimdallarchaeia.

The cytochrome bc1 complex contains subunits with heme groups that bind and transfer electrons from quinol to cytochrome c[91] while pumping protons to the periplasm. We failed to identify clear homologs of this complex in any of the Asgard ancestors. However, at least one subunit of CIII is homologous to a subunit of the NR. For example, the nitrate reductase gamma subunit (NarI, arCOG02194) is a membrane-embedded heme-iron unit that resembles cytochrome b and is capable of accepting electrons from quinols. We inferred the NarI only in the Hodarchaeales ancestor. We also uncovered a potential Rieske iron-sulphur protein (arCOG01720) in most of the Asgard archaeal ancestors (except from Njordarchaeales). These observations provide two potential hypotheses for predicting the function of the ETC in the Asgard archaeal ancestors. Firstly, the nitrate reductase found in the Hodarchaeales ancestor was not acting in conjunction with a Rieske center and the reduction of nitrate to nitrite could have occurred on the plasma membrane facing the cytoplasm[92]. The second possibility is that the Hodarchaeales ancestor used the NarI subunit of a nitrate reductase and the Rieske centre in a membrane-bound complex, to shuttle electrons from quinones to cupredoxin. In this scenario, the Hodarchaeales ancestor is the only ancestor predicted to possess a CIII-like complex.

**Complex IV**. The fourth complex, cytochrome c oxidase, oxidizes cytochrome c from CIII ultimately reducing oxygen to water while pumping protons. This complex is composed of three main subunits (I, II and III) that are conserved throughout evolution. Given its wide distribution across the tree of life, this complex is thought to have been present in the last universal common ancestor of all lineages and likely emerged prior to the rise of global oxygen levels[93]. This observation implies that cytochrome c oxidase might have used an alternative electron acceptor to oxygen. We found that the Heimdallarchaeia ancestor encoded at least subunits I and II while the Hodarchaeales ancestor encoded all three subunits (Supplementary Figure 14). These CIVs might have used electrons from cupredoxins (or other electron carrier molecules) to reduce nitrate to nitrite while pumping protons. Whether this complex could also use oxygen as a terminal electron in the Asgard archaea cannot be determined with present data. However modern Asgard archaeal genomes have been recovered from metagenome data generated from oxic zones[88] where oxygen could be used as an electron acceptor.

**Complex V.** The last component of the ETC is complex V, an ATPase that can harness the proton motive force to biosynthesize ATP. There are three distinct types of ATPases found across the tree of life: the ATP-producing F-type (found in Bacteria, mitochondria and chloroplasts), ATP-consuming V-type (vacuolar-type, common in eukaryotic intracellular membrane compartments), and ATP-producing A-type (archaeal-type, found in the plasma membrane of Archaea and some Bacteria)[94]. We found that all of the Asgard archaeal ancestors likely coded A/V-type ATPase. V- and A-ATPases have similar structures and differ from the F-ATPases by having additional peripheral stalks and connecting subunits $V_1$ and $V_o$. However, our inferences suggest that subunit H, which encodes the peripheral stalk necessary for rotational movements[94], was likely absent in all the Asgard archaeal ancestors.

**Cupredoxin, a potential ancestral electron carrier.** We did not detect genes related to the electron carrier molecule cytochrome *c* in any of the Asgard archaeal ancestors. However, we did identify potential quinone-utilizing proteins, such as CI and CII, and a cupredoxin. In bacteria, cupredoxins have been shown to act as electron donors to terminal oxidases[95,96]. Thus, cupredoxins could have played a key role in shuttling electrons across the membrane-anchored energy-converting complexes of the ETC of ancestral Asgard archaeal lineages.

**Beta oxidation.** All enzymes required for the beta-oxidation and fatty acid biosynthesis pathway are found in the Asgard ancestors suggesting that LAsCA had the ability to degrade and, potentially, synthesize fatty acids. Although the direction of this reaction remains unknown, this is consistent with a scenario in which LAsCA had the ability to degrade organic compounds that were transferred to one or more syntrophic partners[26]. In eukaryotes and bacteria, fatty acids are synthesized or oxidized on two different carrier molecules, acyl-carrier protein (ACP) or CoA respectively. However, in most archaea, fatty acids are likely synthesized and oxidized using CoA[97,98]. ACP has been previously reported in several

Heimdall- and Thorarchaeia genomes[26], yet we could not predict homologs of this protein (IPR003231) in the Asgard archaeal ancestors, indicating that these were acquired during the evolution of these two phyla.

**Aerobic degradation of tryptophan in Heimdallarchaeia ancestors.** Related to oxygen dependence, we further inspected the presence of the enzymes responsible for the aerobic degradation of tryptophan via the kynurenine pathway which has been shown to be present in three Heimdallarchaeia genomes[88]. The two protein families that are specific to this pathway (3-hydroxyanthranilate 3,4-dioxygenase and tryptophan 2,3-dioxygenase) were not found in any other Asgard archaea apart from the three previously described Heimdallarchaeia genomes. The results from the ancestral reconstruction suggest that this pathway was absent in the last common ancestor of Asgard archaea but was acquired later by the ancestor of Heimdallarchaeia.

##### 6.1.3. Proteins of bacterial origin represent a minority of ancestral Asgard archaeal proteomes

As part of the ALE reconciliation analysis, we aimed to identify genes of bacterial origin, as described in the Supplementary Methods. In summary, only ~25% of the 2148 clusters inferred to have originated in the various Asgard archaeal ancestors had a one-to-one correspondence to an EggNOG cluster at the LUCA level, suggesting that most of the others correspond to Asgard archaea innovations. For the 426 clusters that had a one-to-one correspondence to an EggNOG cluster, we aimed to identify a putative source of transfer by placing the sequences from these clusters onto the corresponding NOG trees using epa-ng[2], and extracted the most likely internal placement point. Most of the inferred placements yielded only a vague taxonomic label (‘Bacteria’ was the most frequent one), indicating either a complex history involving multiple transfers across prokaryotes, or an ancestral protein family shared between

Bacteria and Archaea (vertically inherited) that has been lost in non-Asgard archaea. In addition, the investigation of the functional assignment of laterally acquired and *de novo* clusters that had an EggNOG annotation, but most of those corresponded to an "unknown function". Although these results do yield clear patterns regarding the evolutionary origin and function of the genes that originated in Asgard archaeal ancestors, they nonetheless suggest that the majority of those are Asgard archaea innovations, whose function will warrant further investigation.

Reverse gyrase. Our reconciliation analyses suggest that reverse gyrase, a protein found ubiquitously in hyperthermophilic organisms but absent from mesophiles, was already present in the Last Asgard Common Ancestor (Supplementary Table 4). However, the evolution of this protein family is notoriously complex, and given that the Asgard homologs do not form a monophyletic clade in our phylogeny (Supplementary Figure 32), we cannot exclude that the reverse gyrase was acquired several times independently at the base of Asgard hyperthermophilic lineages.

36. Farag, I. F., Zhao, R. & Biddle, J. F. 'Sifarchaeota' a novel Asgard phylum from Costa Rica sediment capable of polysaccharide degradation and anaerobic methylotrophy. *Appl. Environ.*

*Microbiol.* (2021).
37. Varghese, N. J. *et al.* Microbial species delineation using whole genome sequences. *Nucleic Acids Research* vol. 43 6761–6771 Preprint at https://doi.org/10.1093/nar/gkv657 (2015).
38. Barco, R. A. *et al.* A Genus Definition for Bacteria and Archaea Based on a Standard Genome Relatedness Index. *MBio* **11**, (2020).
39. Parks, D. H., Imelfort, M., Skennerton, C. T., Hugenholtz, P. & Tyson, G. W. CheckM: assessing the quality of microbial genomes recovered from isolates, single cells, and metagenomes. *Genome Res.* **25**, 1043–1055 (2015).
40. Narrowe, A. B. *et al.* Complex Evolutionary History of Translation Elongation Factor 2 and Diphthamide Biosynthesis in Archaea and Parabasalids. *Genome Biol. Evol.* **10**, 2380–2393 (2018).
41. Zhao, Y. & Garcia, B. A. Comprehensive Catalog of Currently Documented Histone Modifications. *Cold Spring Harb. Perspect. Biol.* **7**, a025064 (2015).
42. Mattiroli, F. *et al.* Structure of histone-based chromatin in Archaea. *Science* **357**, 609–612 (2017).
43. Henneman, B., van Emmerik, C., van Ingen, H. & Dame, R. T. Structure and function of archaeal histones. *PLoS Genet.* **14**, e1007582 (2018).
44. de Mendoza, A. & Sebé-Pedrós, A. Origin and evolution of eukaryotic transcription factors. *Curr. Opin. Genet. Dev.* **58-59**, 25–32 (2019).
45. Kusmierczyk, A. R. & Hochstrasser, M. Some assembly required: dedicated chaperones in eukaryotic proteasome biogenesis. *Biol. Chem.* **389**, 1143–1151 (2008).
46. Wang, L. & Dobberstein, B. Oligomeric complexes involved in translocation of proteins across the membrane of the endoplasmic reticulum. *FEBS Lett.* **457**, 316–322 (1999).
47. Pfeffer, S. *et al.* Dissecting the molecular organization of the translocon-associated protein complex. *Nat. Commun.* **8**, 14516 (2017).
48. Dacks, J. B. & Robinson, M. S. Outerwear through the ages: evolutionary cell biology of vesicle coats. *Curr. Opin. Cell Biol.* **47**, 108–116 (2017).
49. Hirst, J. *et al.* Characterization of TSET, an ancient and widespread membrane trafficking

complex. *Elife* **3**, e02866 (2014).

50. Shaik, S., Pandey, H., Thirumalasetti, S. K. & Nakamura, N. Characteristics and Functions of the Yip1 Domain Family (YIPF), Multi-Span Transmembrane Proteins Mainly Localized to the Golgi Apparatus. *Front Cell Dev Biol* **7**, 130 (2019).
51. Heidtman, M., Chen, C. Z., Collins, R. N. & Barlowe, C. Yos1p Is a Novel Subunit of the Yip1p–Yif1p Complex and Is Required for Transport between the Endoplasmic Reticulum and the Golgi Complex. *MBoC* **16**, 1673–1683 (2005).
52. Christensen, J. R. *et al.* Cytoplasmic dynein-1 cargo diversity is mediated by the combinatorial assembly of FTS-Hook-FHIP complexes. *Elife* **10**, (2021).
53. Roy, M. & Roux, S. Rab GTPases in Osteoclastic Bone Resorption and Autophagy. *Int. J. Mol. Sci.* **21**, (2020).
54. Seaman, M. N. J. The retromer complex - endosomal protein recycling and beyond. *J. Cell Sci.* **125**, 4693–4702 (2012).
55. Kim, E. *et al.* Implication of mouse Vps26b–Vps29–Vps35 retromer complex in sortilin trafficking. *Biochem. Biophys. Res. Commun.* **403**, 167–171 (2010).
56. Shi, H., Rojas, R., Bonifacino, J. S. & Hurley, J. H. The retromer subunit Vps26 has an arrestin fold and binds Vps35 through its C-terminal domain. *Nat. Struct. Mol. Biol.* **13**, 540–548 (2006).
57. Suzuki, S. W., Chuang, Y.-S., Li, M., Seaman, M. N. J. & Emr, S. D. A bipartite sorting signal ensures specificity of retromer complex in membrane protein recycling. *J. Cell Biol.* **218**, 2876–2886 (2019).
58. Wiederhold, E. *et al.* The yeast vacuolar membrane proteome. *Mol. Cell. Proteomics* **8**, 380–392 (2009).
59. Villaseñor, R., Kalaidzidis, Y. & Zerial, M. Signal processing by the endosomal system. *Curr. Opin. Cell Biol.* **39**, 53–60 (2016).
60. Graham, S. C. *et al.* Structural basis of Vps33A recruitment to the human HOPS complex by Vps16. *Proc. Natl. Acad. Sci. U. S. A.* **110**, 13345–13350 (2013).
61. Jiang, P. *et al.* The HOPS complex mediates autophagosome–lysosome fusion through interaction with syntaxin 17. *Molecular Biology of the Cell* vol. 25 1327–1337 Preprint at

https://doi.org/10.1091/mbc.e13-08-0447 (2014).
62. Balderhaar, H. J. K. & Ungermann, C. CORVET and HOPS tethering complexes - coordinators of endosome and lysosome fusion. *J. Cell Sci.* **126**, 1307–1316 (2013).
63. Arlt, H., Perz, A. & Ungermann, C. An overexpression screen in Saccharomyces cerevisiae identifies novel genes that affect endocytic protein trafficking. *Traffic* **12**, 1592–1603 (2011).
64. Vild, C. J. & Xu, Z. Vfa1 binds to the N-terminal microtubule-interacting and trafficking (MIT) domain of Vps4 and stimulates its ATPase activity. *J. Biol. Chem.* **289**, 10378–10386 (2014).
65. Azmi, I. *et al.* Recycling of ESCRTs by the AAA-ATPase Vps4 is regulated by a conserved VSL region in Vta1. *J. Cell Biol.* **172**, 705–717 (2006).
66. Dimaano, C., Jones, C. B., Hanono, A., Curtiss, M. & Babst, M. Ist1 regulates Vps4 localization and assembly. *Mol. Biol. Cell* **19**, 465–474 (2008).
67. Liewen, H. *et al.* Characterization of the human GARP (Golgi associated retrograde protein) complex. *Exp. Cell Res.* **306**, 24–34 (2005).
68. Pérez-Victoria, F. J. *et al.* Structural basis for the wobbler mouse neurodegenerative disorder caused by mutation in the Vps54 subunit of the GARP complex. *Proc. Natl. Acad. Sci. U. S. A.* **107**, 12860–12865 (2010).
69. Schluter, C. *et al.* Global analysis of yeast endosomal transport identifies the vps55/68 sorting complex. *Mol. Biol. Cell* **19**, 1282–1294 (2008).
70. Alsleben, S. & Kölling, R. Vps68 cooperates with ESCRT-III in intraluminal vesicle formation. *J. Cell Sci.* **135**, (2022).
71. Tseng, C.-C. *et al.* Correction: Bro1 stimulates Vps4 to promote intralumenal vesicle formation during multivesicular body biogenesis. *Journal of Cell Biology* vol. 220 Preprint at https://doi.org/10.1083/jcb.20210207007022021c (2021).
72. Imachi, H. *et al.* Isolation of an archaeon at the prokaryote-eukaryote interface. *bioRxiv* 726976 (2019) doi:10.1101/726976.
73. Sheridan, P. O. *et al.* Gene duplication drives genome expansion in a major lineage of Thaumarchaeota. *Nat. Commun.* **11**, 1–12 (2020).
74. Martijn, J. *et al.* Hikarchaeia demonstrate an intermediate stage in the methanogen-to-halophile

transition. *Nat. Commun.* **11**, 5490 (2020).

75. Williams, T. A. *et al.* Integrative modeling of gene and genome evolution roots the archaeal tree of life. *Proceedings of the National Academy of Sciences* **114**, E4602–E4611 (2017).
76. Say, R. F. & Fuchs, G. Fructose 1,6-bisphosphate aldolase/phosphatase may be an ancestral gluconeogenic enzyme. *Nature* **464**, (2010).
77. Bräsen, C., Esser, D., Rauch, B. & Siebers, B. Carbohydrate Metabolism in Archaea: Current Insights into Unusual Enzymes and Pathways and Their Regulation. *Microbiol. Mol. Biol. Rev.* **78**, 89 (2014).
78. The oxidative pentose phosphate pathway in the haloarchaeon Haloferax volcanii involves a novel type of glucose-6-phosphate dehydrogenase – The archaeal Zwischenferment. *FEBS Lett.* **589**, 1105–1111 (2015).
79. FEBS Press. https://febs.onlinelibrary.wiley.com/doi/epdf/10.1016/j.febslet.2015.03.026.
80. Soderberg, T. Biosynthesis of ribose-5-phosphate and erythrose-4-phosphate in archaea: a phylogenetic analysis of archaeal genomes. *Archaea* **1**, 347 (2005).
81. -J. Weiße, R. H., Faust, A., Schmidt, M., Schönheit, P. & Scheidig, A. J. Structure of NDP-forming Acetyl-CoA synthetase ACD1 reveals a large rearrangement for phosphoryl transfer. *Proc. Natl. Acad. Sci. U. S. A.* **113**, E519–E528 (2016).
82. Spaans, S. K., Weusthuis, R. A., Van Der Oost, J. & Kengen, S. W. M. NADPH-generating systems in bacteria and archaea. *Front. Microbiol.* **6**, (2015).
83. Zhu, G., Brian Golding, G. & Dean, A. M. The Selective Cause of an Ancient Adaptation. *Science* **307**, 1279–1282 (2005).
84. Liu, Y. *et al.* Comparative genomic inference suggests mixotrophic lifestyle for Thorarchaeota. *ISME J.* **12**, 1021–1031 (2018).
85. Kono, T. *et al.* A RuBisCO-mediated carbon metabolic pathway in methanogenic archaea. *Nat. Commun.* **8**, 1–12 (2017).
86. Greening, C. *et al.* Genomic and metagenomic surveys of hydrogenase distribution indicate H2 is a widely utilised energy source for microbial growth and survival. *ISME J.* **10**, 761–777 (2016).
87. Vanwonterghem, I. *et al.* Methylotrophic methanogenesis discovered in the archaeal phylum

Verstraetearchaeota. *Nature Microbiology* **1**, 1–9 (2016).

### Supplementary figures 1-32

**NB. High resolution versions for all Supplementary Figures is available in separate PDF file (see zip file 'Supplementary_figures_2021-04-06336R1.zip').**

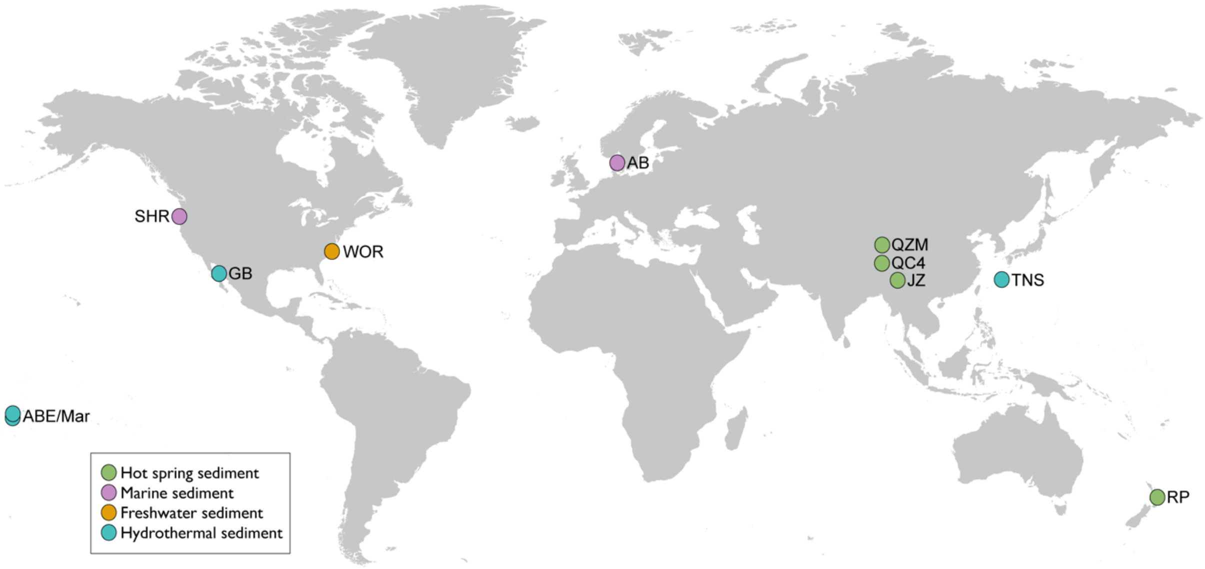


**Supplementary Figure 1.** World map showing the sampling locations of the MAGs described in the current study. SHR: South Hydrate Ridge; AB: Aarhus Bay (Denmark); ABE: ABE vent field; GB: Guaymas Basin (Mexico); JZ: Jinze (China); Mar: Mariner vent field; QC: QuCai village (China); QZM: QuZhuoMu village (China); RP: Radiata Pool, Ngatamariki (New Zealand); TNS: Taketomi Island (Japan).

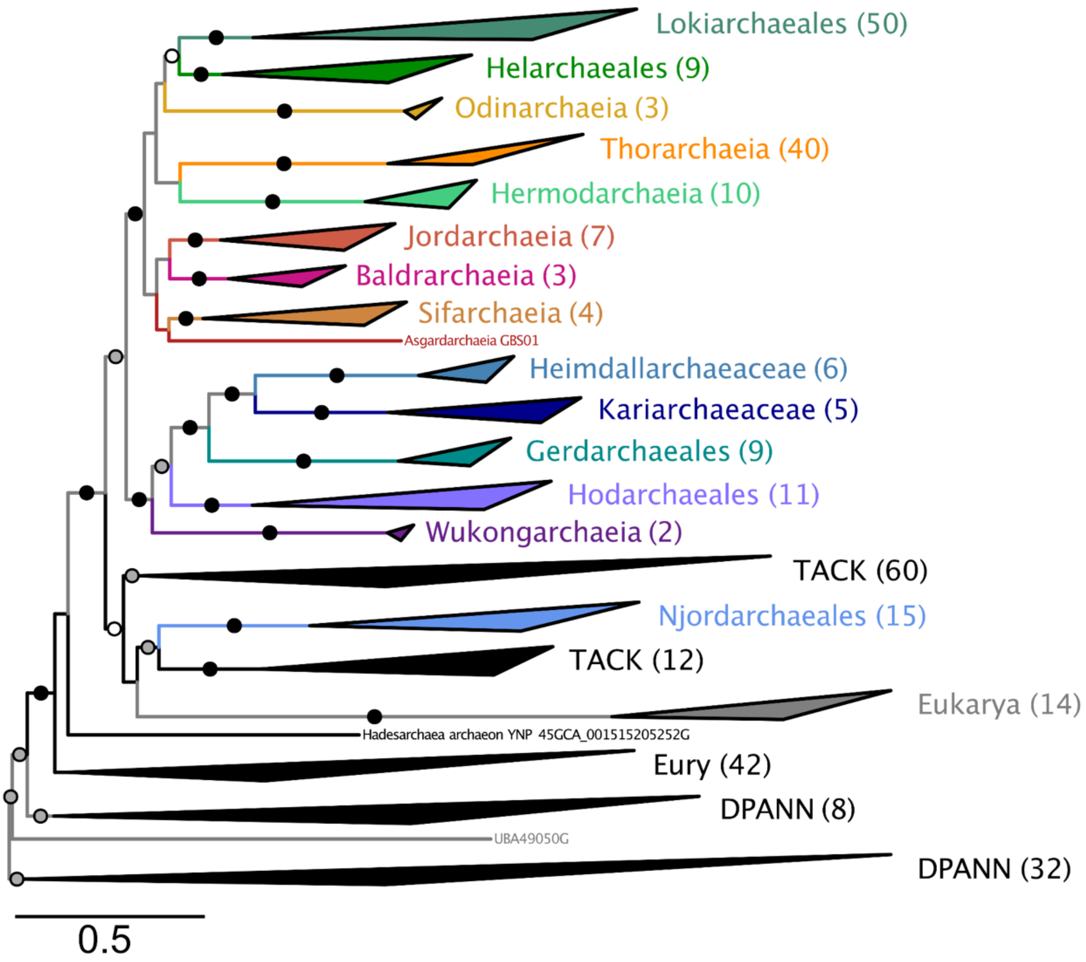


**Supplementary Figure 2**. Phylogenomic analysis based on 56 concatenated ribosomal proteins reveals 11 major Asgard archaea clades. Phylogeny inferred from the RP56-A175 dataset (7112 sites and 345 taxa), using IQ-TREE under the LG+C60+F+Γ model. Support at branches was estimated using the PMSF bootstrap approximation under the same model (100 pseudo-replicates). Black dots indicate maximum support values (100%), grey dots indicate bootstrap support 95-99% and white dots bootstrap support 70-95%. Tree is midpoint rooted. Scale bar denotes the average expected substitutions per site. These analyses revealed the existence of 12 major Asgard archaeal clades: the previously described Lokiarchaeales, Odinarchaeia, Heimdallarchaeia (comprising the orders Gerdarchaeales and Hodarchaeales, and the families Kariarchaeaceae and Heimdallarchaeaceae), Thorarchaeia and Helarchaeales, the recently reported Hermod-, Sif, Jord, Baldr and Wukongarchaeia, and Njordarchaeales as well as the newly identified class Asgardarchaeia. While they correspond to different taxonomic ranks, which we discuss in the Supplementary Information, we decided to adhere as closely as possible to the clade compositions as they were discussed in the literature, while revising their suffixes. Uncollapsed tree is available in Figshare.

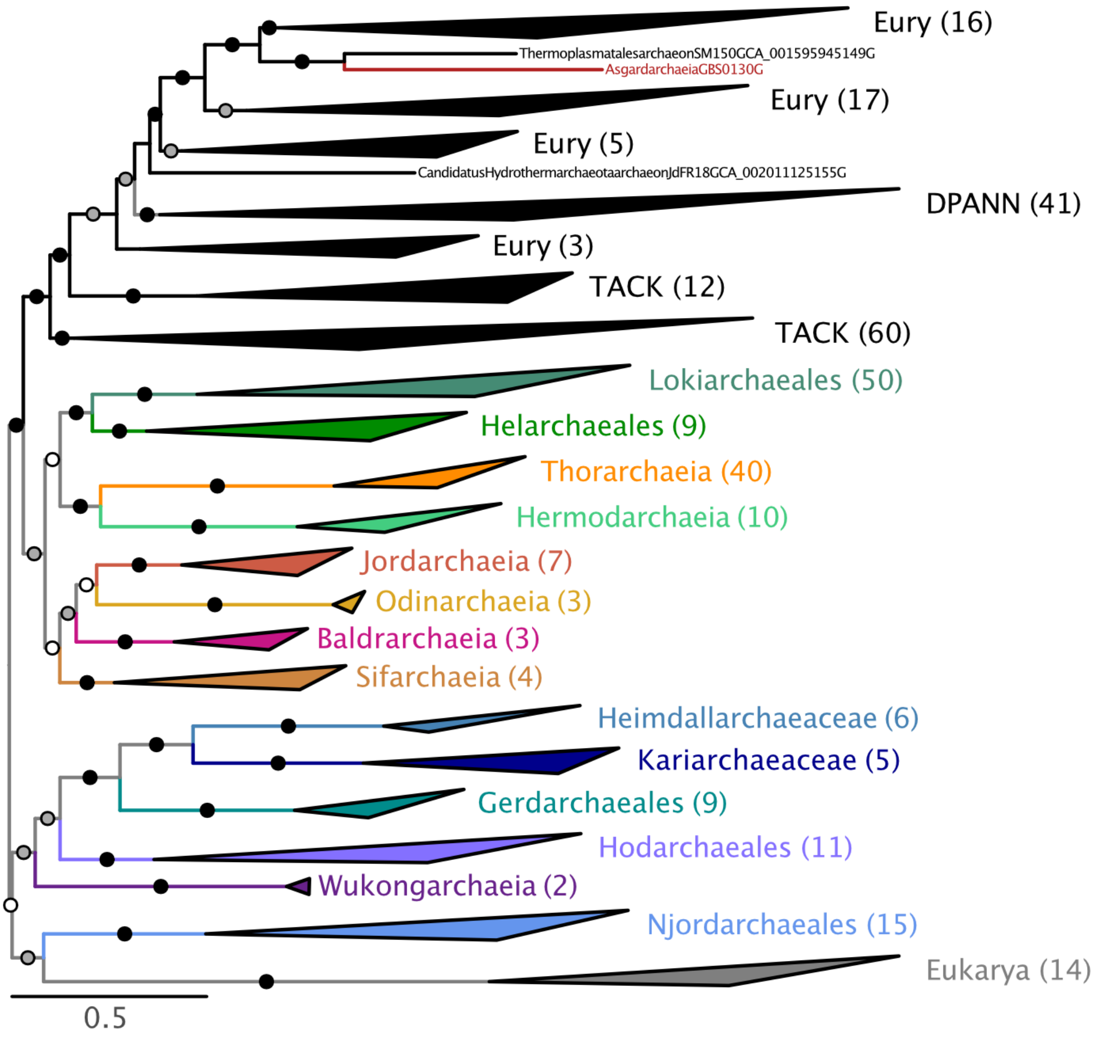


**Supplementary Figure 3.** Maximum likelihood phylogenomic analysis of the NM-A175 dataset (57 concatenated proteins, 15733 sites after trimming, 345 taxa). Phylogeny was inferred using IQ-TREE under the LG+C60+F+Γ model. Support at branches was estimated using the PMSF bootstrap approximation under the same model (100 pseudoreplicates). Scale bar denotes the average expected substitutions per site. The uncollapsed tree is available in Figshare.

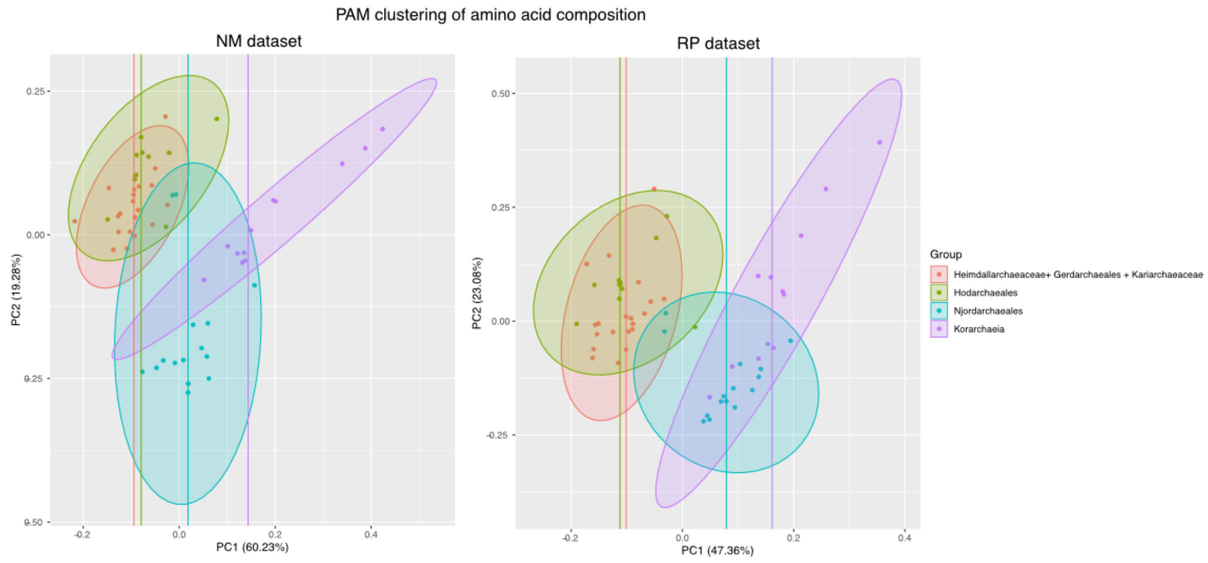


**Supplementary Figure 4.** PAM clustering amino acid composition. PAM (Partition Around Medoids) clustering of amino acid compositions in the NM and RP gene datasets. Colors indicate taxonomy as indicated in the legend.

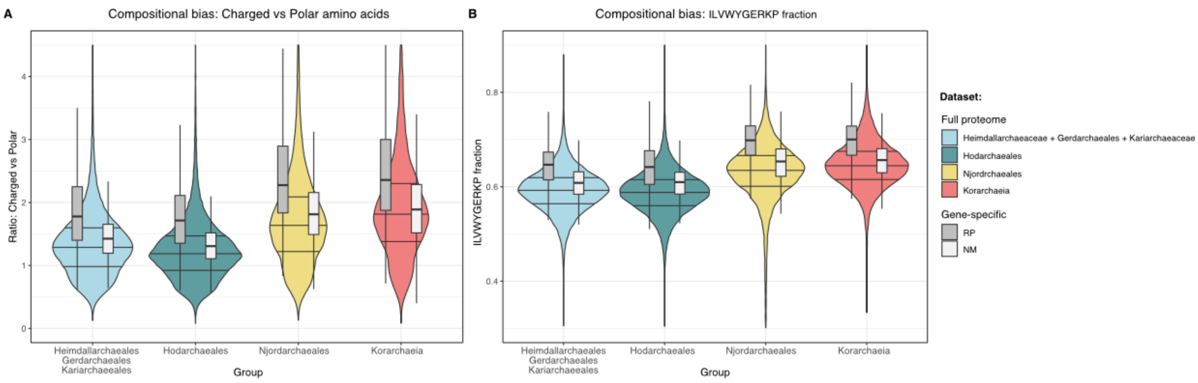


**Supplementary Figure 5.** Thermostability compositional patterns -- violin plots. Thermostability-related compositional bias is represented by (A) the ratio of charged versus polar amino acids and (B) the fraction of residues represented by the amino acids isoleucine, leucine, valine, tryptophan, tyrosine, glycine, glutamate, arginine, lysine, and proline. Background violin plots represent the distribution of values in the entire proteome of all genomes included in the four groups under focus. Boxplots represent the distribution of values corresponding to the NM and RP datasets.

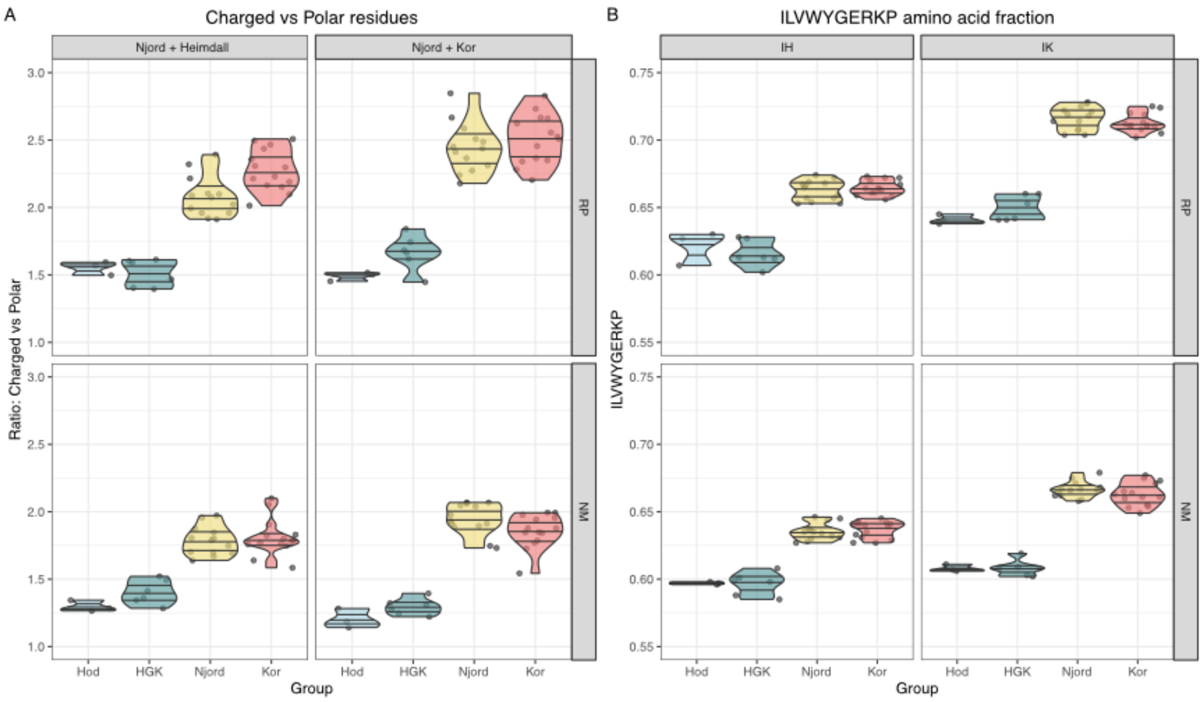


**Supplementary Figure 6.** Thermostability compositional patterns on sites with topological preferences. Thermostability-related compositional bias is represented by (A) the ratio of charged versus polar amino acids and (B) the fraction of residues represented by the amino acids isoleucine, leucine, valine, tryptophan, tyrosine, glycine, glutamate, arginine, lysine, and proline. Left and right-side plots represent sites with a higher site-likelihood for a topology in which Njord clusters with other Heimdallarchaeia (left) or Korarchaeia (right). The top and bottom plots represent sites from the RP (top) and NM (bottom) datasets. Dots and their distributions represent compositional features for genomes within each taxonomical category.

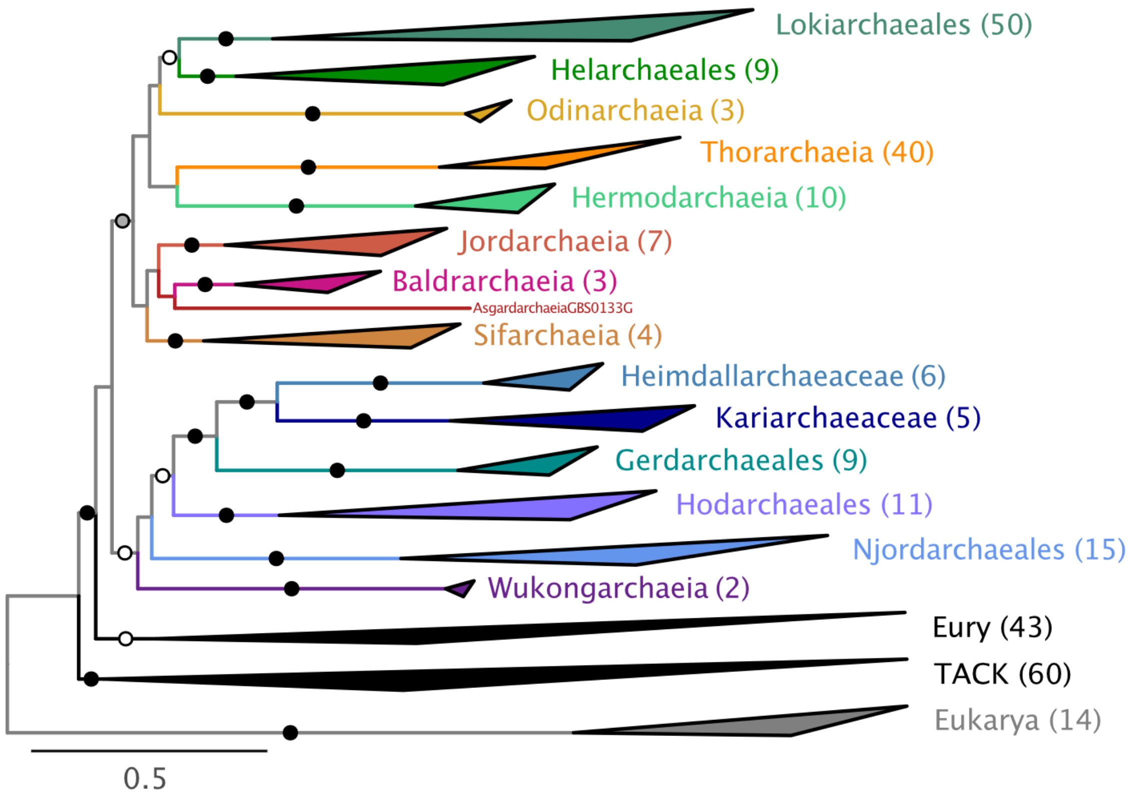


**Supplementary Figure 7.** Phylogenomic analysis based on the RP56-A175-nDK dataset (7093 sites and 292 taxa), using IQ-TREE under the LG+C60+F+Γ model. DPANN and Korarchaeota have been discarded. Support at branches was estimated using the PMSF bootstrap approximation under the same model (100 pseudoreplicates). Black dots indicate maximum support values (100%), grey dots indicate bootstrap support 95-99%, and white dots bootstrap support 70-95%. The tree is midpoint rooted. Scale bar denotes the average expected substitutions per site. Note the shift in the position of Njordarchaeales which now branch at the base of Heimdallarchaeia instead of as sister to Korarchaeota (Figure S2). Uncollapsed phylogeny is available in Figshare.

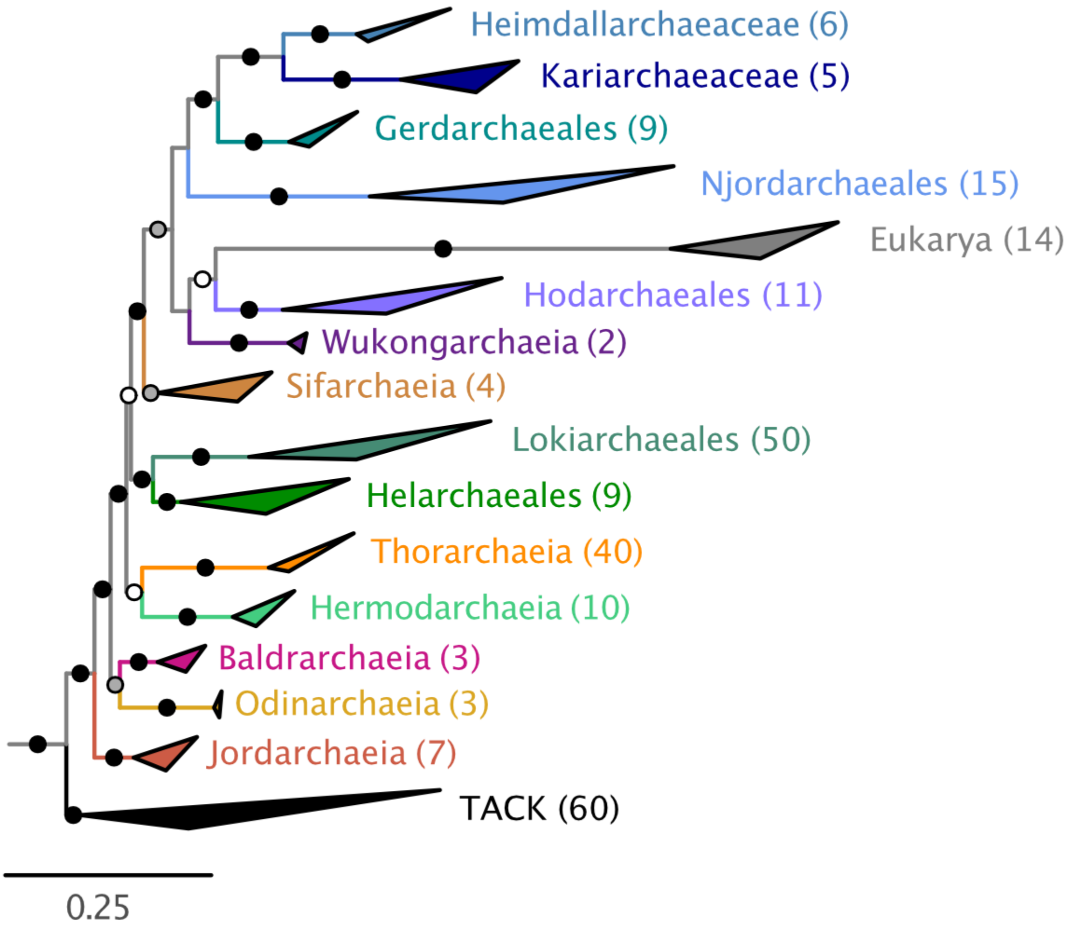


**Supplementary Figure 8.** Phylogenomic analysis based on the NM57-A175-nDK_SR4_FSR20 dataset (12584 sites and 292 taxa), using IQ-TREE under a user-defined model referred to as 'C60SR4' as described in[23]. DPANN and Korarchaeota have been discarded. Alignment was SR4-recoded and the 20% fastest-evolving sites were removed. Support at branches was estimated using the PMSF bootstrap approximation under the same model (100 pseudo-replicates). Black dots indicate maximum support values (100%), grey dots indicate bootstrap support 95-99%, and white dots bootstrap support 70-95%. The tree is midpoint rooted. Scale bar denotes the average expected substitutions per site. Euryarchaeota were pruned for the figure. Uncollapsed phylogeny is available in Figshare.

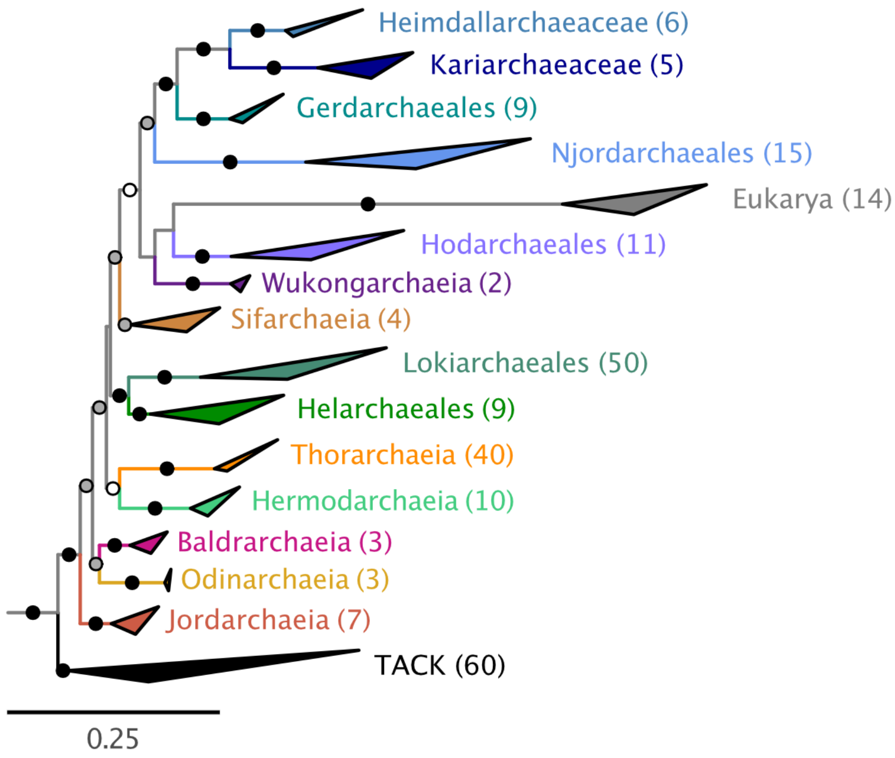


**Supplementary Figure 9.** Maximum-likelihood phylogenomic analysis based on the NM57-A175-nDK_SR4_FSR30 dataset (11011 sites and 292 taxa), using IQ-TREE under a user-defined model referred to as 'C60SR4' as described in[23]. DPANN and Korarchaeota have been discarded. Alignment was SR4-recoded and the 30% fastest-evolving sites were removed. Support at branches was estimated using the PMSF bootstrap approximation under the same model (100 pseudo-replicates). Black dots indicate maximum support values (100%), grey dots indicate bootstrap support 95-99% and white dots bootstrap support 70-95%. The tree is midpoint rooted. Scale bar denotes the average expected substitutions per site. Euryarchaeota were pruned for the figure. Note the nested position of Njordarchaeales within Heimdallarchaeia (BS = 95% for the monophyly of Njordarchaeales, Gerdarchaeales, Kariarchaeaceae and Heimdallarchaeaceae). Uncollapsed phylogeny is available in Figshare.

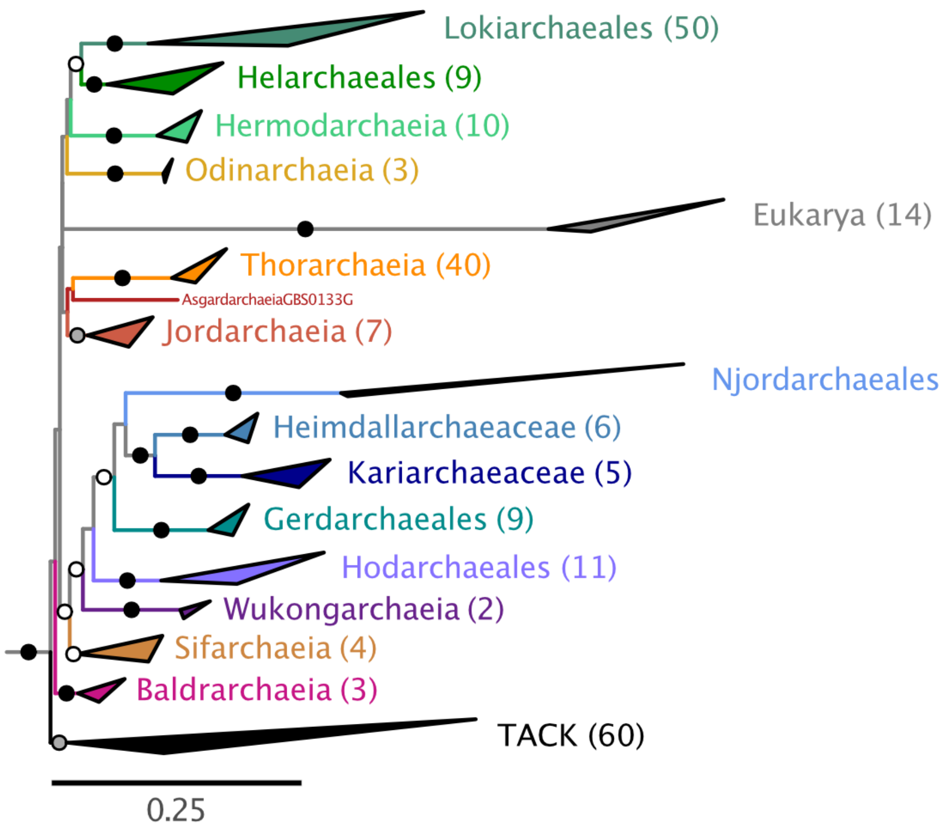


**Supplementary Figure 10.** Maximum-likelihood phylogenomic analysis based on the RP56-A175-nDK_SR4_FSR30 dataset (4977 sites and 292 taxa), using IQ-TREE under a user-defined model referred to as 'C60SR4' as described in[23]. DPANN and Korarchaeota have been discarded. Alignment was SR4-recoded and the 30% fastest-evolving sites removed. Support at branches was estimated using the PMSF bootstrap approximation under the same model (100 pseudo-replicates). Black dots indicate maximum support values (100%), grey dots indicate bootstrap support 95-99% and white dots bootstrap support 70-95%. Tree is midpoint rooted. Scale bar denotes the average expected substitutions per site. Euryarchaeota were pruned for the figure. Note the nested position of Njordarchaeales within Heimdallarchaeia (BS = 83% for the monophyly of Njordarchaeales, Gerdarchaeales, Kariarchaeaceae and Heimdallarchaeaceae). Uncollapsed phylogeny is available in Figshare.

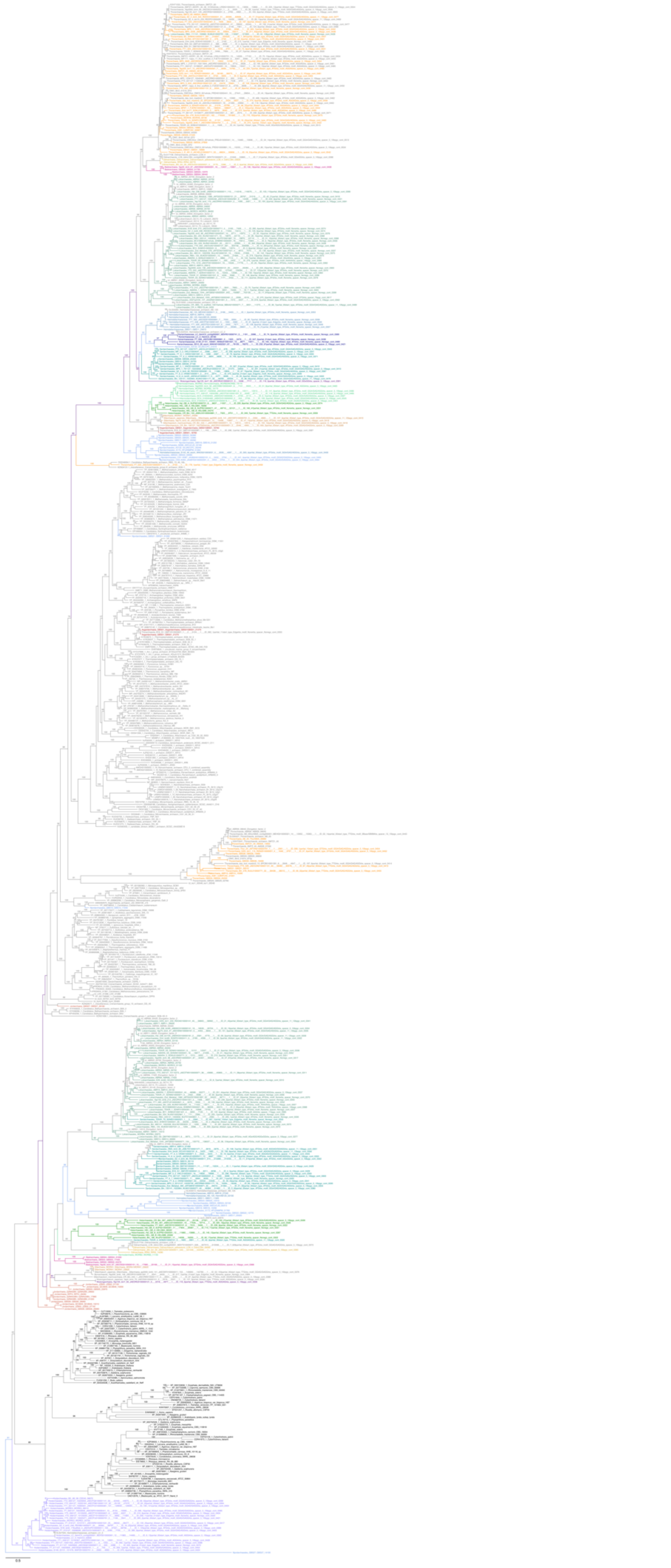

1765

**Supplementary Figure 11.** Maximum likelihood phylogeny of the EF-2 protein family. Homologs identified in our set of Asgard archaea genomes (coloured) were added to the sequences obtained from Narrowe, et al. 2018[40] (in grey), aligned with mafft-linsi and trimmed with BMGE (-m BLOSUM30 -b 3). Tree was inferred using IQ-TREE under the LG+C20+F+Γ model. Support at branches was estimated using ultrafast bootstrap (1000 pseudo-replicates). Support values <80% are not displayed. Scale bar denotes the average expected substitutions per site. Hodarchaeales members possess a single homolog (at the bottom, with eukaryotes in black) displaying the diphthamide modification motif.

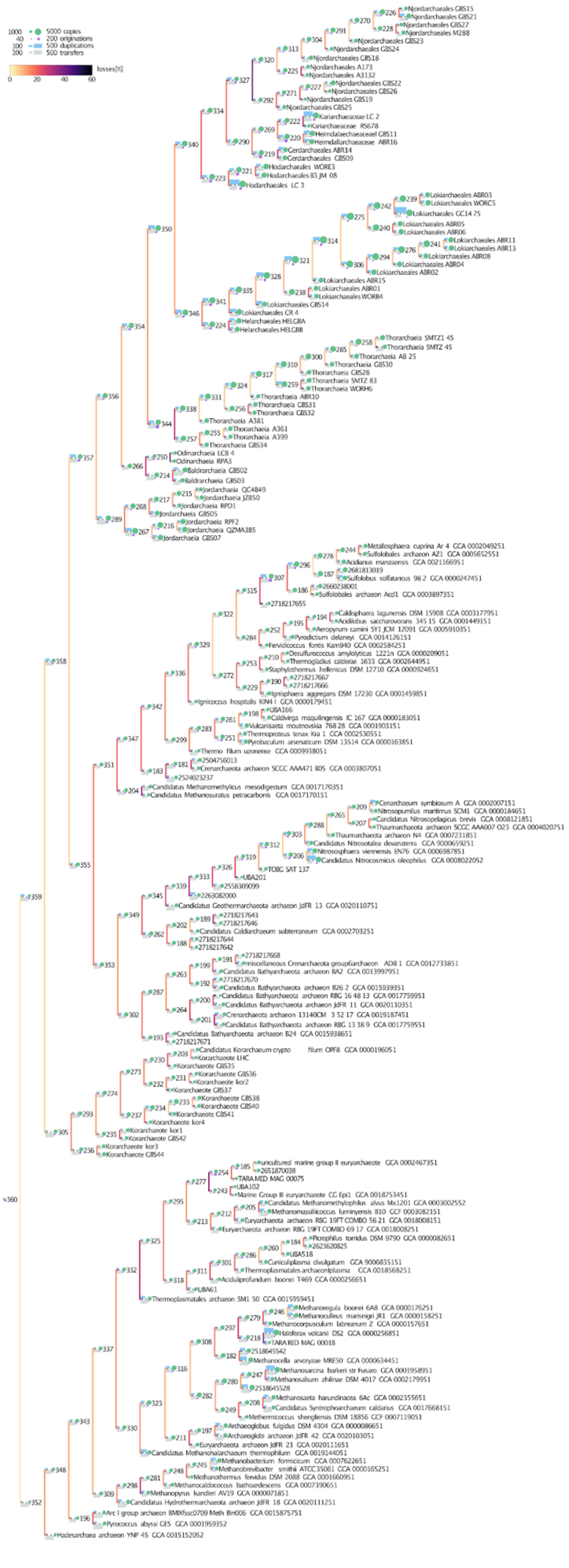

**Supplementary Figure 12.** Ancestral proteome sizes and numbers of gene loss, duplication, and gain were inferred from reconciliation analyses using ALE on the A64 dataset, across a selection of archaeal representatives belonging to Euryarchaeota, TACK and Asgard archaea. This figure was generated using a modified version of https://github.com/Boussau/plotODTLTree/blob/master/PlotTreeWithODTL.Rmd. The underlying data for this figure, including values for proteome size, and for gene transfer, origination, duplication, and loss events for each node (specified per node number) are provided in Supplementary Table 8.

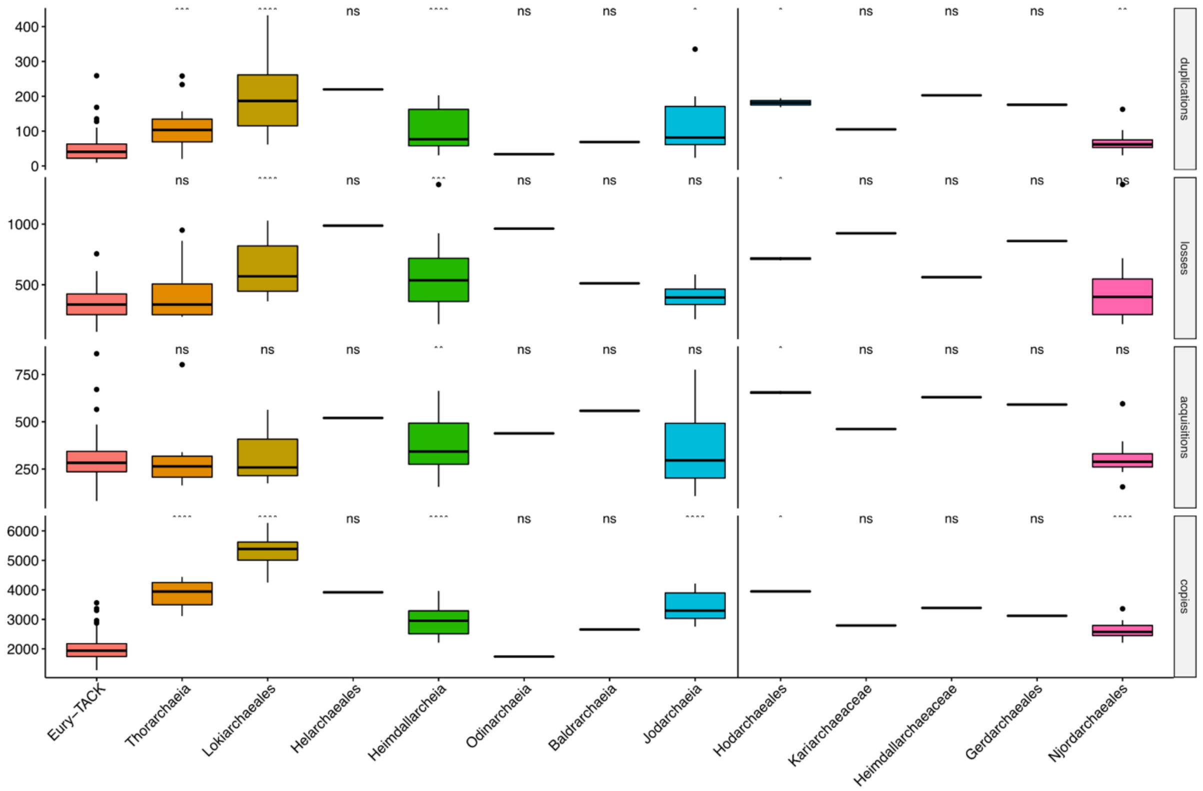


**Supplementary Figure 13.** Absolute number of predicted events inferred for Asgard archaeal ancestors, plotted by major clade. P-values for each Wilcoxon test against the median values of internal nodes belonging to TACK and Euryarchaeota are shown above each category, where *: p-value <= 0.05, **: p-value <=0.01, ***: p-value <=0.001. Most Asgard archaeal ancestors had predicted proteome size (i.e. protein copy numbers) which are significantly higher than other archaea. This can be in parts explained by the numbers of gene duplications which is itself estimated to have been significantly higher in several Asgard archaea clade ancestors compared to Euryarchaeota and TACK archaea.

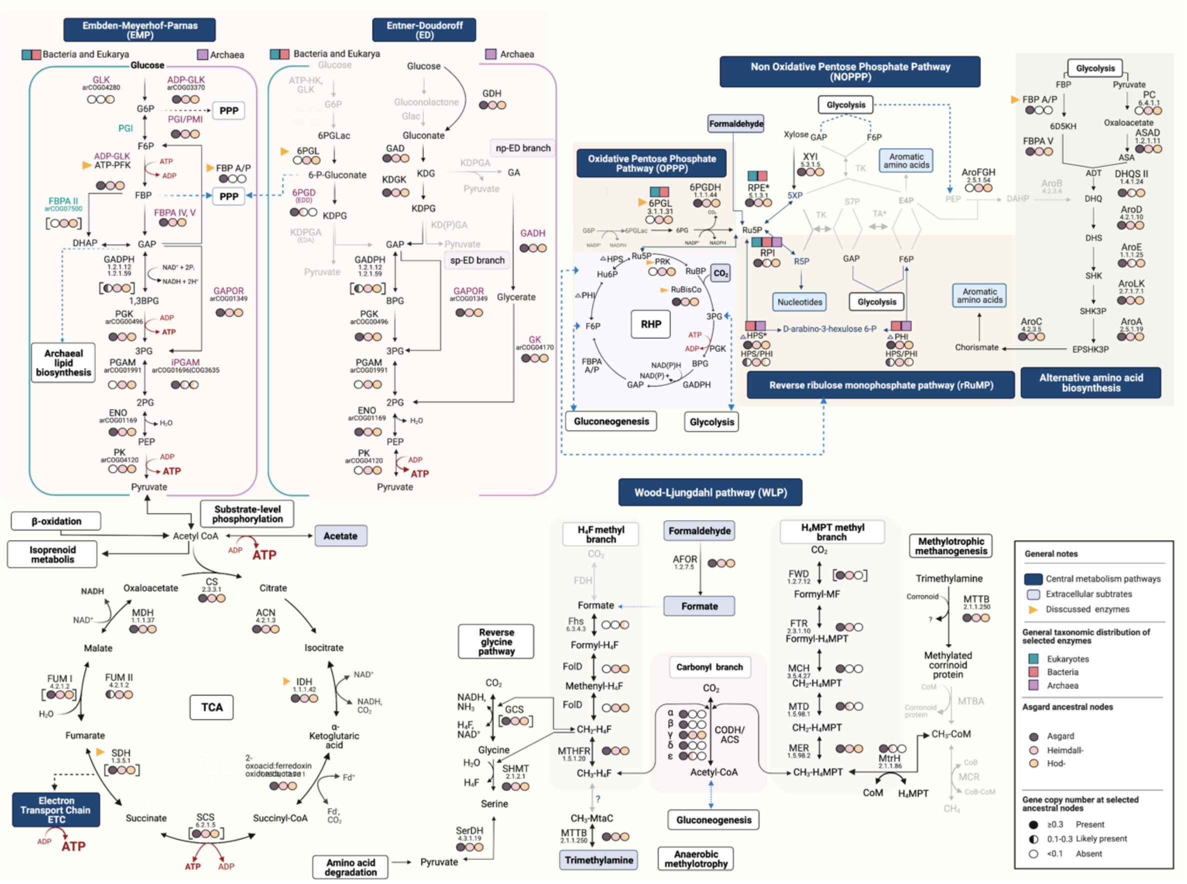


**Supplementary Figure 14.** Metabolic map of several Asgard ancestors. Carbon metabolism pathways (glycolysis, gluconeogenesis, pentoses phosphate pathway, amino acid biosynthesis, and Wood-Ljungdahl pathway) are indicated on a blue background. All the enzymatic steps with the corresponding enzyme name abbreviations and E.C. number can be cross-referenced in Supplementary Table 4. Coloured circles indicate whether an enzyme was likely present (filled circle), possibly present (half filled), and likely absent (empty circle) in the ancestor of all Asgard archaea, all Heimdallarchaeia (including Njordarchaeales), and of Hodarchaeales only, from left to right, respectively. Square brackets around coloured circles indicate consensus annotation of a given enzymatic complex. Phylogenetic distribution of selected enzymes mainly found in eukaryotes, Bacteria and Archaea are shown in green, red and purple squares. Potential extracellular compounds that can be used as a source of carbon or energy are highlighted in dark gray (acetate, amino acids and formaldehyde, formate, $CO_2$, trimethylamine: TMA). Key enzymes that display a clear variation in the copy numbers of theother investigated Asgard ancestors compared to Hodarchaeales are shown in yellow triangles. Enzymes and arrows displayed in light grey indicate steps that could not be identified in any Asgard ancestor. Blue dashed arrows indicate substrates that are supplied for, or derived from other central pathways. ATP production steps are highlighted in red. Central carbon pathways from sugar degradation and synthesis are represented by the Embden-Meyerhof-Parnas (EMP) and Entner-Doudoroff (ED) pathways, including the non-phosphorylative branch (np-ED branch) and the semi-phosphorylative branch (sp-ED) in Archaea. The reactions known for Bacteria and eukaryotes and the modified versions in Archaea are shown in green and pink, respectively. Enzymes displayed in black in the EMP and ED pathways represent shared steps in Bacteria, eukaryotes and Archaea. EMP and ED pathways were modified from [77]. The Pentose phosphate pathway (PPP) is divided into nonoxidative pentose phosphate pathway (NOPPP), oxidative pentose phosphate pathway (OPPP) and the reversed ribulose monophosphate pathway (rRuMP). An alternative aromatic amino acid biosynthesis pathway that does not involve Erythrose 4-Phosphate (E4P) as precursors includes 2-Amino-3,7-dideoxy-D-threo-hept-6-ulosonic acid, ADT; 3-Dehydroquinate, DHQ; 3-Dehydroxyshikimate, DHS; Shikimate, SHK; Shikimate-3-phosphate, SHK3P. The reductive hexulose-phosphate (RHP) pathway that involve ribulose-1,5-bisphosphate carboxylase/oxygenase (RuBisCO), responsible for $CO_2$ fixation, and phosphoribulokinase (PRK) was modified from [85]. Wood-Ljungdahl pathway (WLP) was modified from [99]. Abbreviations: ENO, enolase; FBPA, fructose-1,6-bisphosphate aldolase; GAPDH, glyceraldehyde-3-phosphate dehydrogenase; GAPN, non-phosphorylating GAPDH; GAPOR, GAP:glyceraldehyde-3-phosphate; GLK, glucokinase; HK, hexokinase; PEPS, PEP synthe-tase; PFK, phosphofructokinase; PGI, phosphoglucose isomerase; PGI/PMI, phosphoglucose isomerase/phosphomannose isomerase; cPGI, cupin-type phos- phoglucose isomerase; PGAM, phosphoglycerate mutase (dPGAM, 2,3-bisphosphoglycerate [2,3BPG] cofactor dependent; iPGAM, 2,3BPG cofactor independent); PK, pyruvate kinase; PPDK, pyruvate:phosphate dikinase; G6P, glucose 6-phosphate; F6P, fructose 6-phosphate; DHAP, dihydroxyacetone phosphate; GAP, glyceraldehyde 3-phosphate; 1,3BPG, 1,3-bisphosphoglycerate; 3PG, 3-phosphoglycerate; 2PG, 2-phosphoglycerate; PEP, phosphoenolpyruvate; EDA, Entner-Doudoroff Aldolase; EDD, Entner-Doudoroff dehydratase; $H_4F$, tetrahydrofolate methyl branch; FDH, formate dehydrogenase; FHS, 10-formyl-H4F synthetase; FolD, 5,10-methenyl-H4F cyclohydrolase/ 5,10-methylene-H4F dehydrogenase. $H_4MPT$, $H_4MPT$ methyl branch tetrahydromethanopterin; FWD, Both the tungsten (Fwd) or molybdenum (Fmd) formylmethanofuran dehydrogenase complex (Fwd–FmdABCD); FTR, Formylmethanofuran:H4MPT-formyltransferase; MCH, methenyl-H4MPT cyclohydrolase; MTD, F420-dependent methylene (CH2)-H4 MPT dehydrogenase; MER, methylene-H4 MPT reductase. MTTB, Trimethylamine methyltransferase; GCS, glycine-cleavage system; rRuMP, Reversed ribulose monophosphate pathway; RHP, reductive hexulose-phosphate pathway; RuBisCO, Ribulose-1,5-bisphosphate carboxylase/oxygenase; PRK, phosphoribulokinase. Created with BioRender.com

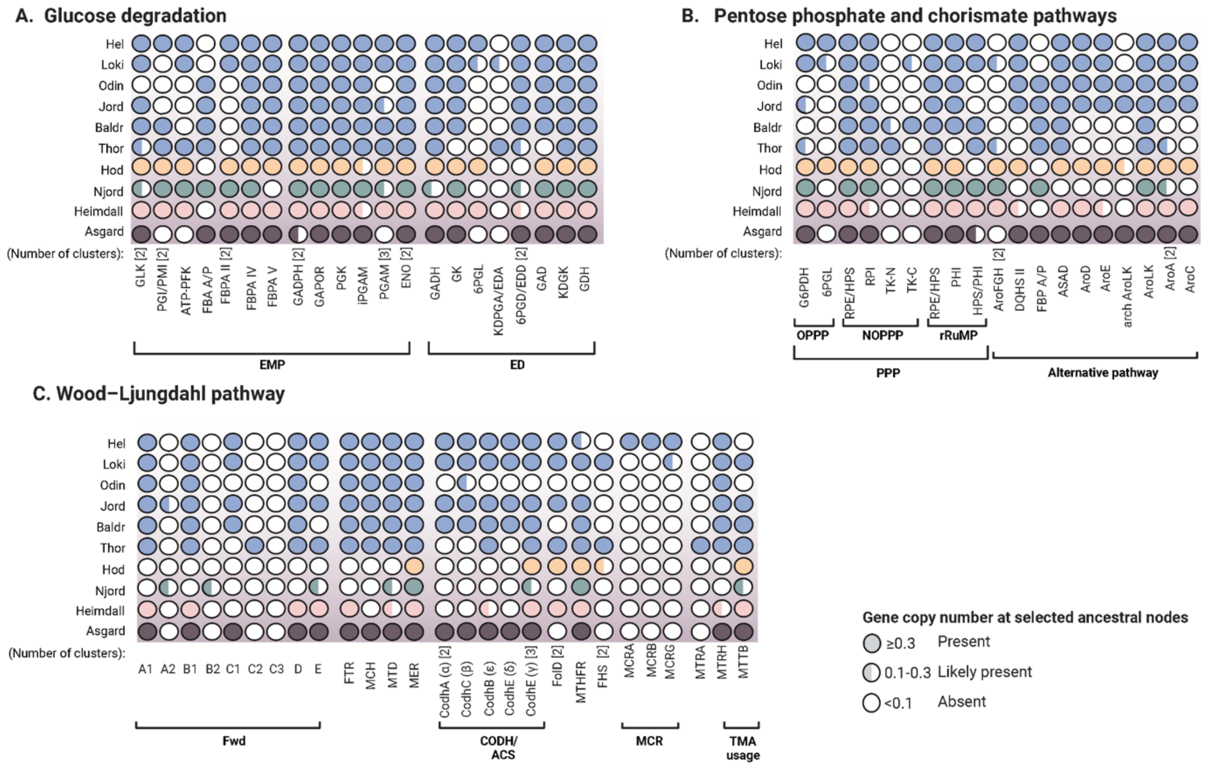


**Supplementary Figure 15. Predicted presence of key enzymes from selected carbon central metabolic pathways.** A) Glucose degradation includes glycolytic pathways and gluconeogenic pathways: Embden-Meyerhof-Parnas (EMP) and Entner-Doudoroff (ED). B) Precursors synthesis includes: Pentoses phosphate pathway (PPP) divided in nonoxidative pentose phosphate pathway (NOPPP), oxidative pentose phosphate pathway (OPPP) and the reversed ribulose monophosphate pathway (rRuMP). The alternative aromatic amino acid biosynthesis pathway is also included. C) Wood-Ljungdahl pathway (WLP). Filled, half-filled and empty circles indicate the inferred presence, potential presence and absence of a given enzymatic step in the ancestral nodes, respectively, based on the predicted copy number at that node. Blue circles indicate Asgard archaea ancestors not included in the main Figure. Enzymatic abbreviations can be cross-reference in Supplementary Figure 15 and Table S7.

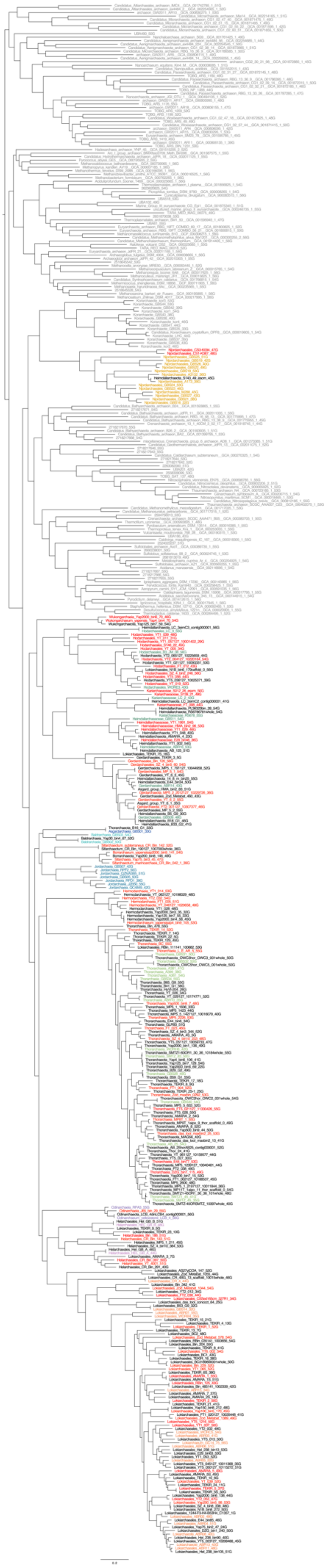

1862

**Supplementary Figure 16. Approximate Maximum likelihood phylogenomic analysis showing the relationship between all Asgard archaeal MAGs and genomes available at the NCBI database as of May 12, 2021, as well as 63 novel MAGs described in the present work.** Phylogenomic reconstruction is based on the RP56-A293 supermatrix (465 taxa including 293 Asgard archaea, 7112 amino acid positions). In the case of closely related lineages, we selected only one representative (based on distance and completeness) for downstream phylogenomic analyses in order to reduce the computational burden. Asgard taxa included in the phylogenomic investigations presented in this work are colored in red if they were part of the larger dataset (RP-A175 and NM-A175), and in other colors if they were included in the smaller dataset (RP-A64 and NM-A64). DPANN archaea, Euryarchaea, and TACK archaea are part of both datasets and indicated in grey. Scale bar denotes the average expected substitutions per site.

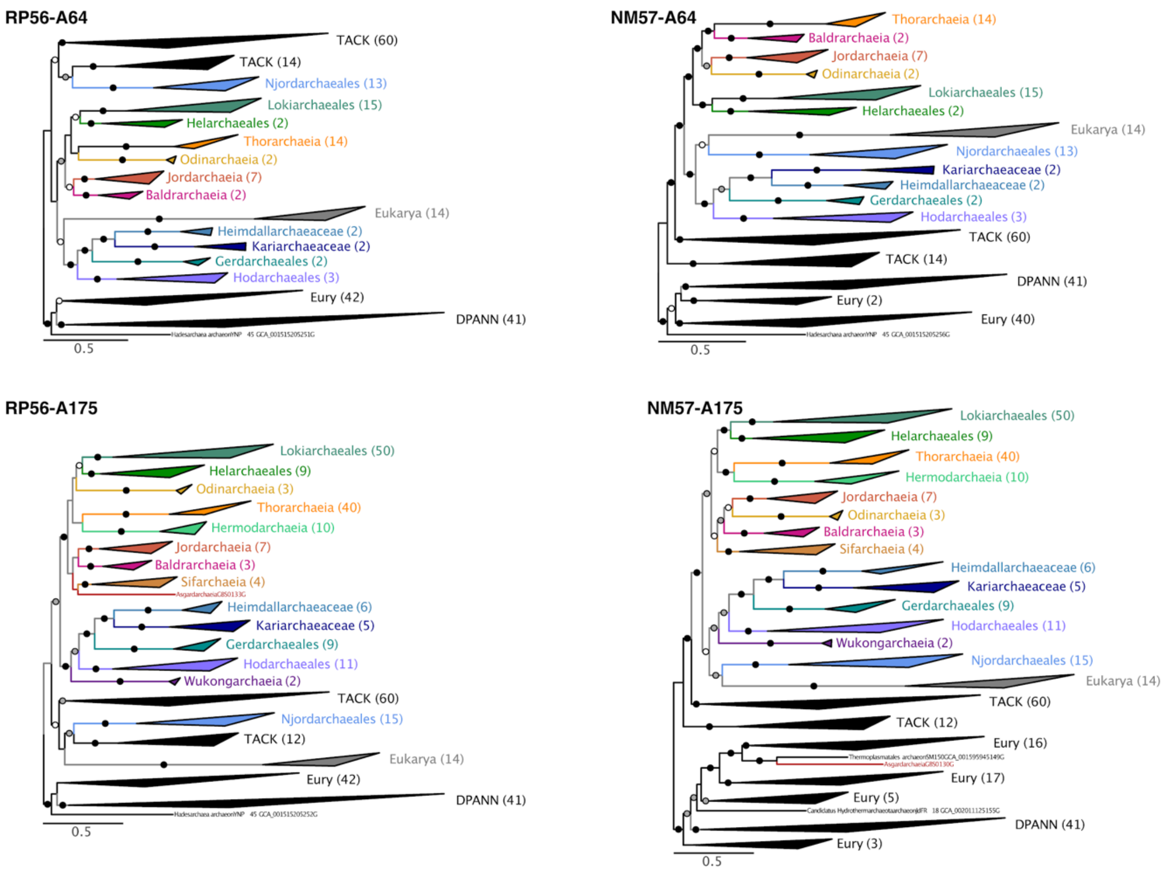


**Supplementary Figure 17**. Maximum-likelihood phylogenomic analysis based on the RP56-A64 (6332 sites, 236 taxa), NM57-A64 (14847 sites, 236 taxa), RP56-A175 (7112 sites, 345 taxa), NM57-A175 (15733 sites, 345 taxa), using IQ-TREE under the LG+C60+F+Γ model. Support at branches was estimated using the PMSF bootstrap approximation under the same model (100 pseudoreplicates). Black dots indicate maximum support values (100%), grey dots indicate bootstrap support 95-99% and white dots bootstrap support 70-95%. Tree is rooted on Euryarchaeota and DPANN. Scale bar denotes the average expected substitutions per site. Uncollapsed phylogenies are available in Figshare.

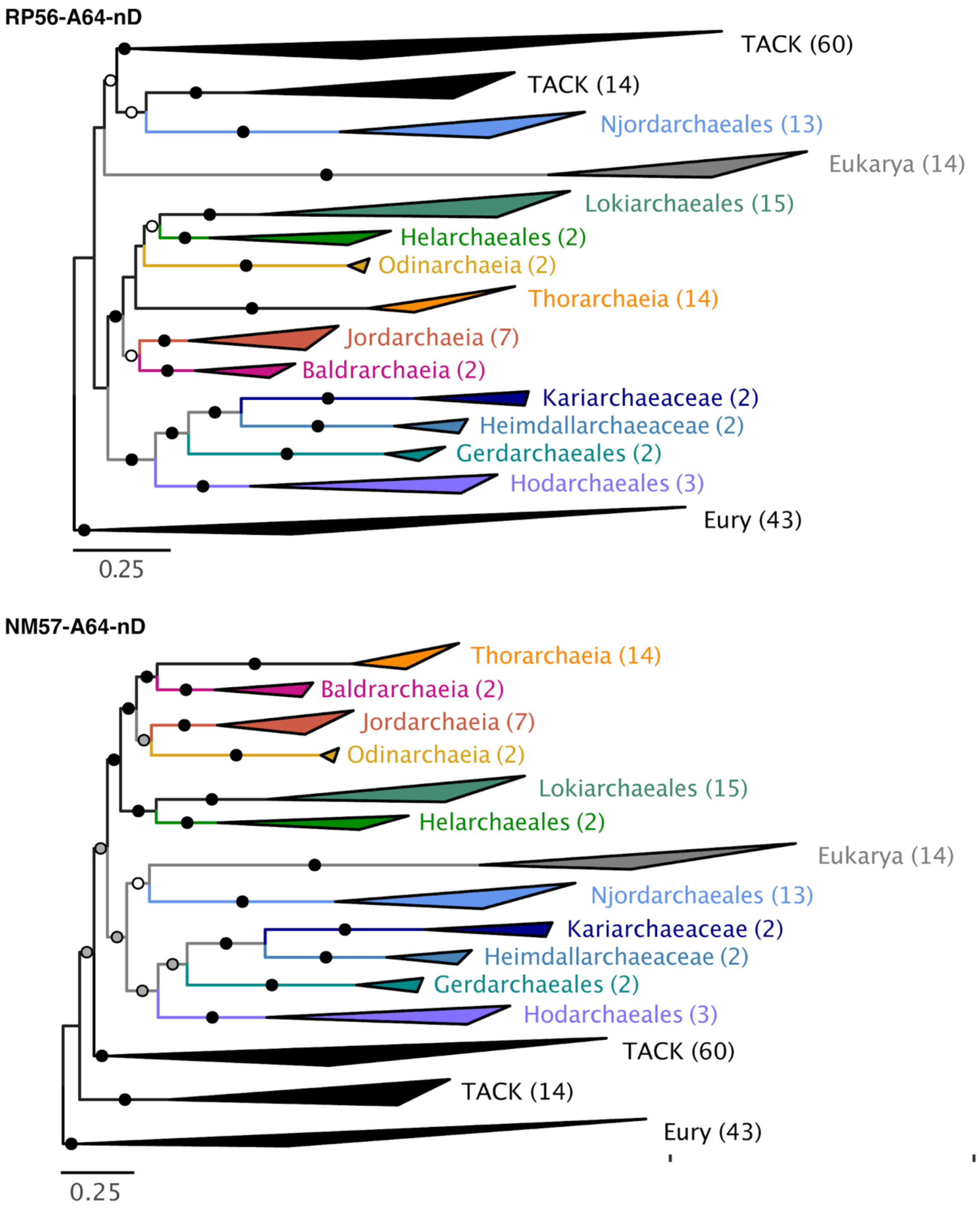


**Supplementary Figure 18**. Maximum-likelihood phylogenomic analysis based on the RP56-A64-nD (5647 sites, 195 taxa) and NM57-A64-nD (13485 sites, 195 taxa), using IQ-TREE under the LG+C60+F+Γ model. DPANN have been discarded. Support at branches was estimated using the PMSF bootstrap approximation under the same model (100 pseudoreplicates). Black dots indicate maximum support values (100%), grey dots indicate bootstrap support 95-99% and white dots bootstrap support 70-95%. Tree is rooted on Euryarchaeota. Scale bar denotes the average expected substitutions per site. Uncollapsed phylogenies are available in Figshare.

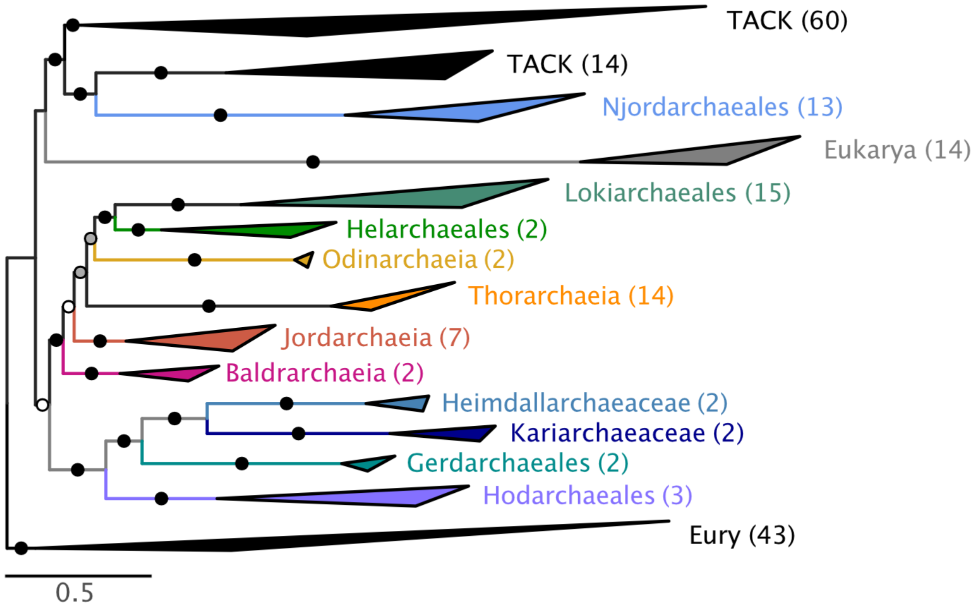


**Supplementary Figure 19**. Bayesian phylogenomic analysis based on the RP56-A64-nD (5647 sites, 195 taxa), using IQ-TREE under the CAT+GTR model. DPANN have been discarded. Black dots indicate maximum support values (posterior probability, PP = 1.0), grey dots indicate PP = 0.95-0.99 and white dots PP = 0.70-0.95. Tree is rooted on Euryarchaeota. Scale bar denotes the average expected substitutions per site. Uncollapsed phylogenies are available in Figshare.

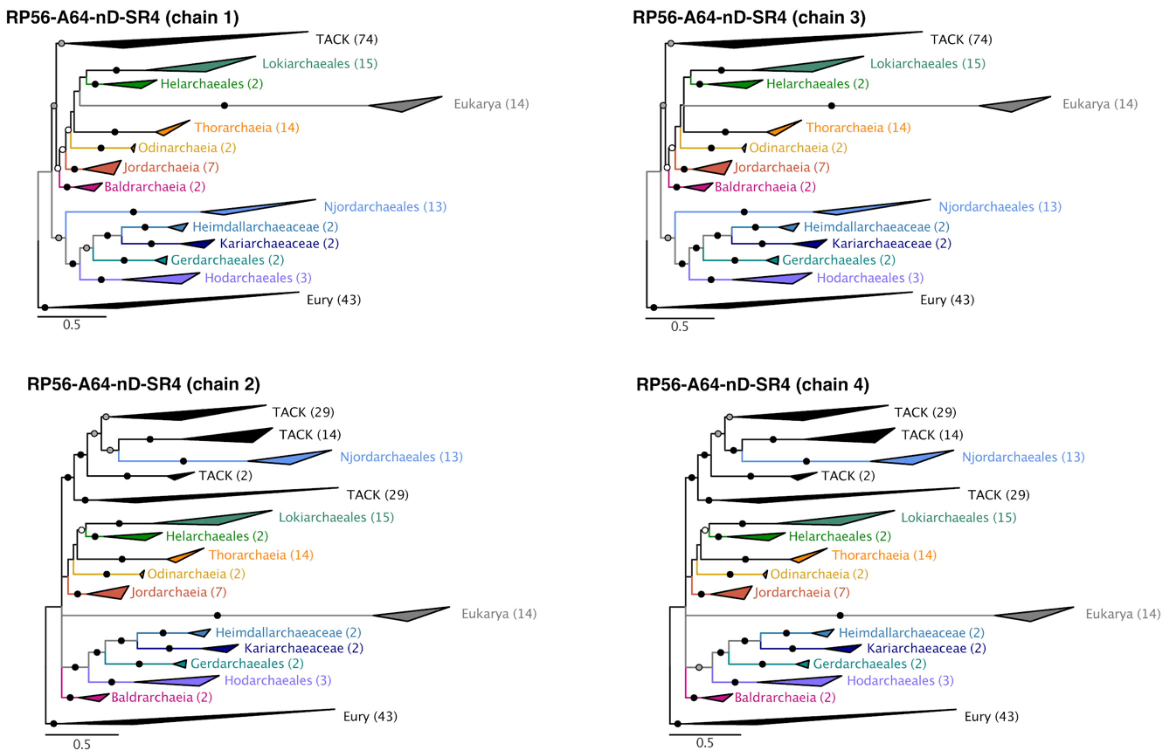


**Supplementary Figure 20**. Bayesian inferences based on the RP56-A64-nD-SR4 (5647 sites, 195 taxa), using IQ-TREE under the CAT+GTR model. Consensus tree for each chains are depicted. Black dots indicate maximum support values (posterior probability, PP = 1.0), grey dots indicate PP = 0.95-0.99 and white dots PP = 0.70-0.95. Trees are rooted on Euryarchaeota.

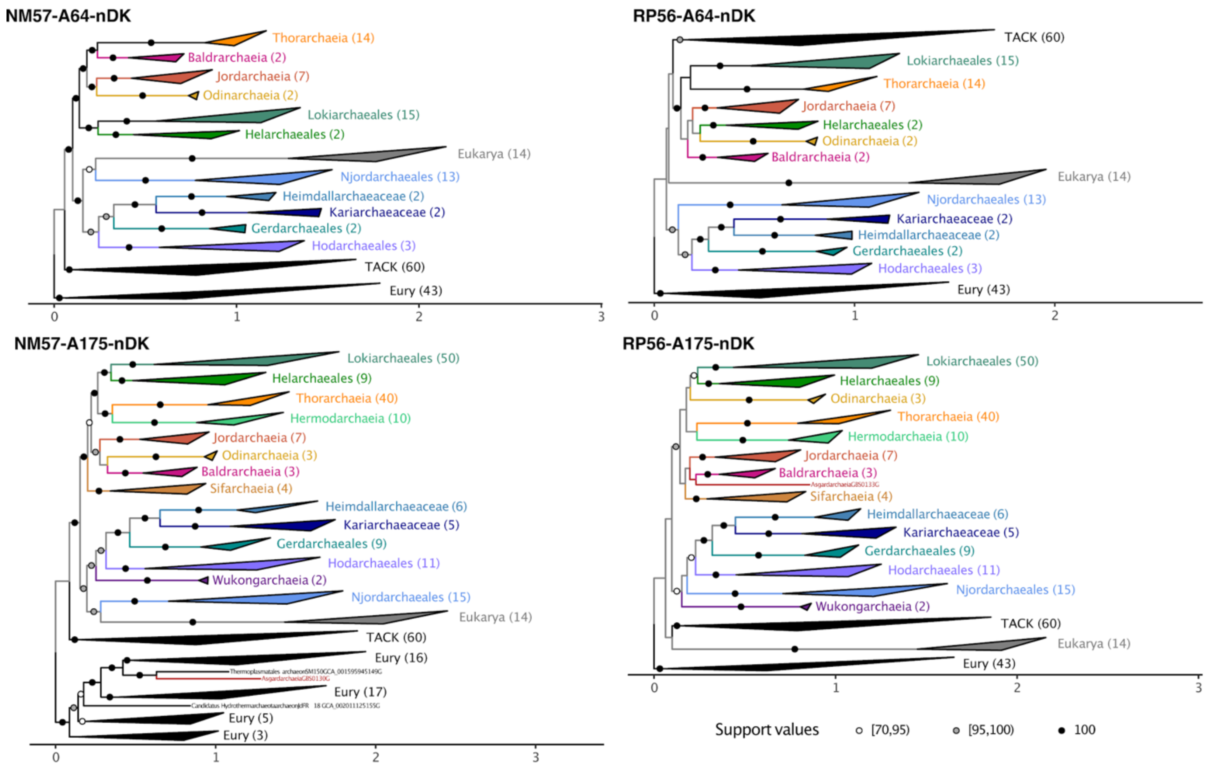


**Supplementary Figure 21**. Maximum-likelihood phylogenomic analysis based on the RP56-A64-nDK (6422 sites, 181 taxa), RP56-A175-nDK (7093 sites, 292 taxa), NM57-A64-nDK (14774 sites, 181 taxa), NM57-A175-nDK (15679 sites, 292 taxa), using IQ-TREE under the LG+C60+F+Γ model. DPANN and Korarchaeota have been discarded. Support at branches was estimated using the PMSF bootstrap approximation under the same model (100 pseudoreplicates). Black dots indicate maximum support values (100%), grey dots indicate bootstrap support 95-99% and white dots bootstrap support 70-95%. Tree is rooted on Euryarchaeota. Scale bar denotes the average expected substitutions per site. Uncollapsed phylogenies are available in Figshare.

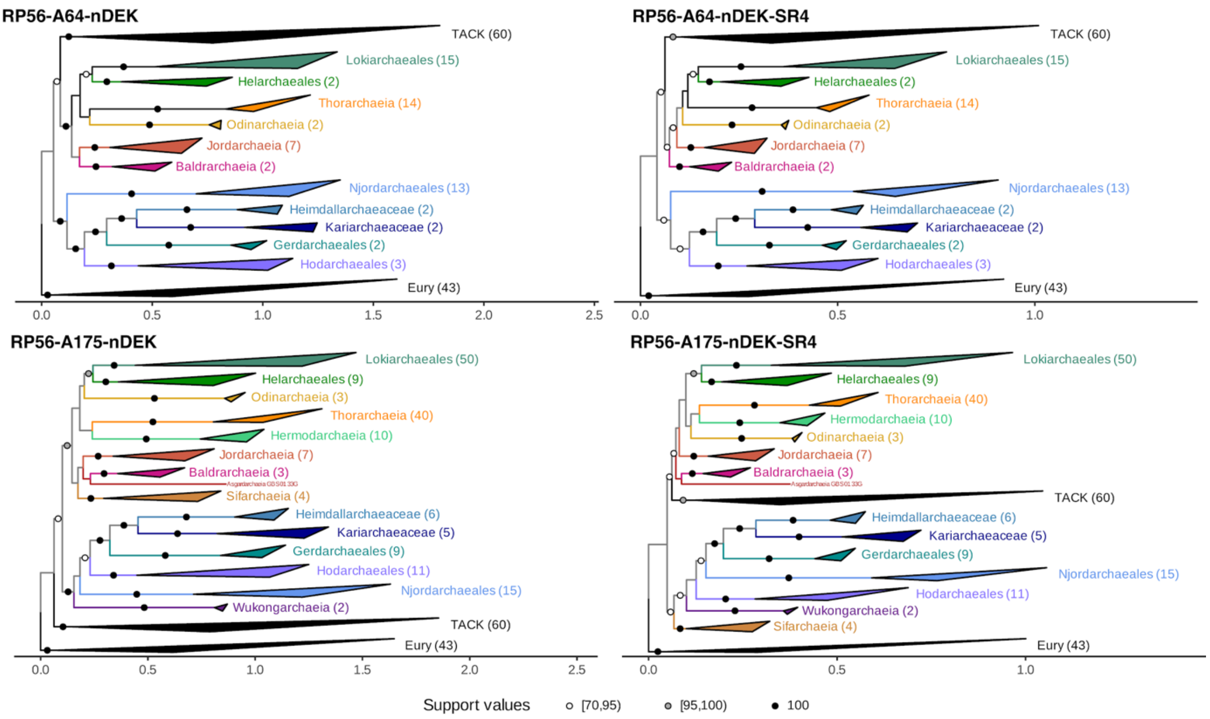


**Supplementary Figure 22.** Maximum-likelihood phylogenomic analysis based on the RP56-A64-nDEK and RP56-A64-nDEK-SR4 (6229 sites, 167 taxa), NM57-A175-nDEK-SR4 and NM57-A175-nDEK (13985 sites, 167 taxa) using IQ-TREE under the LG+C60+F+Γ model for the untreated alignments, and under a user-defined 'C60SR4' model for the SR4-recoded alignments. DPANN, Korarchaeota and eukaryotes have been discarded. Support at branches was estimated using the PMSF bootstrap approximation under the same model (100 pseudoreplicates). Black dots indicate maximum support values (100%), grey dots indicate bootstrap support 95-99% and white dots bootstrap support 70-95%. Tree is rooted on Euryarchaeota. Scale bar denotes the average expected substitutions per site. Uncollapsed phylogenies are available in Figshare.

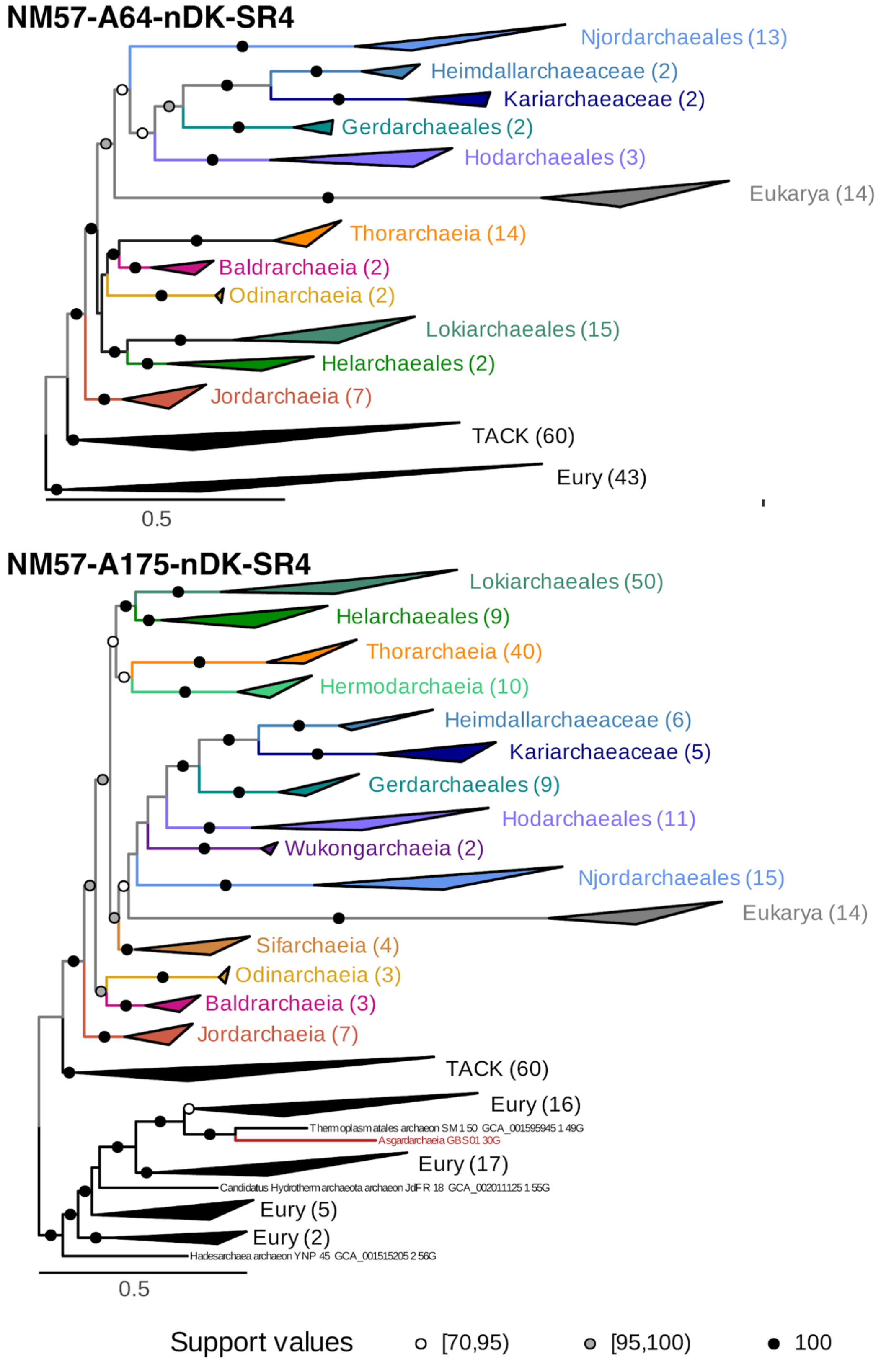


**Supplementary Figure 23.** Maximum-likelihood phylogenomic analysis based on the NM57-A64-nDK-SR4 (14774 sites, 181 taxa) and NM57-A175-nDK-SR4 (15679 sites, 292 taxa)

using IQ-TREE under the LG+C60+F+Γ model for the untreated alignments, and under a user-defined 'C60SR4' model for the SR4-recoded alignments. DPANN, Korarchaeota and eukaryotes have been discarded. Support at branches was estimated using the PMSF bootstrap approximation under the same model (100 pseudoreplicates). Black dots indicate maximum support values (100%), grey dots indicate bootstrap support 95-99%, and white dots bootstrap support 70-95%. Tree is rooted on Euryarchaeota. Scale bar denotes the average expected substitutions per site. Uncollapsed phylogenies are available in Figshare.

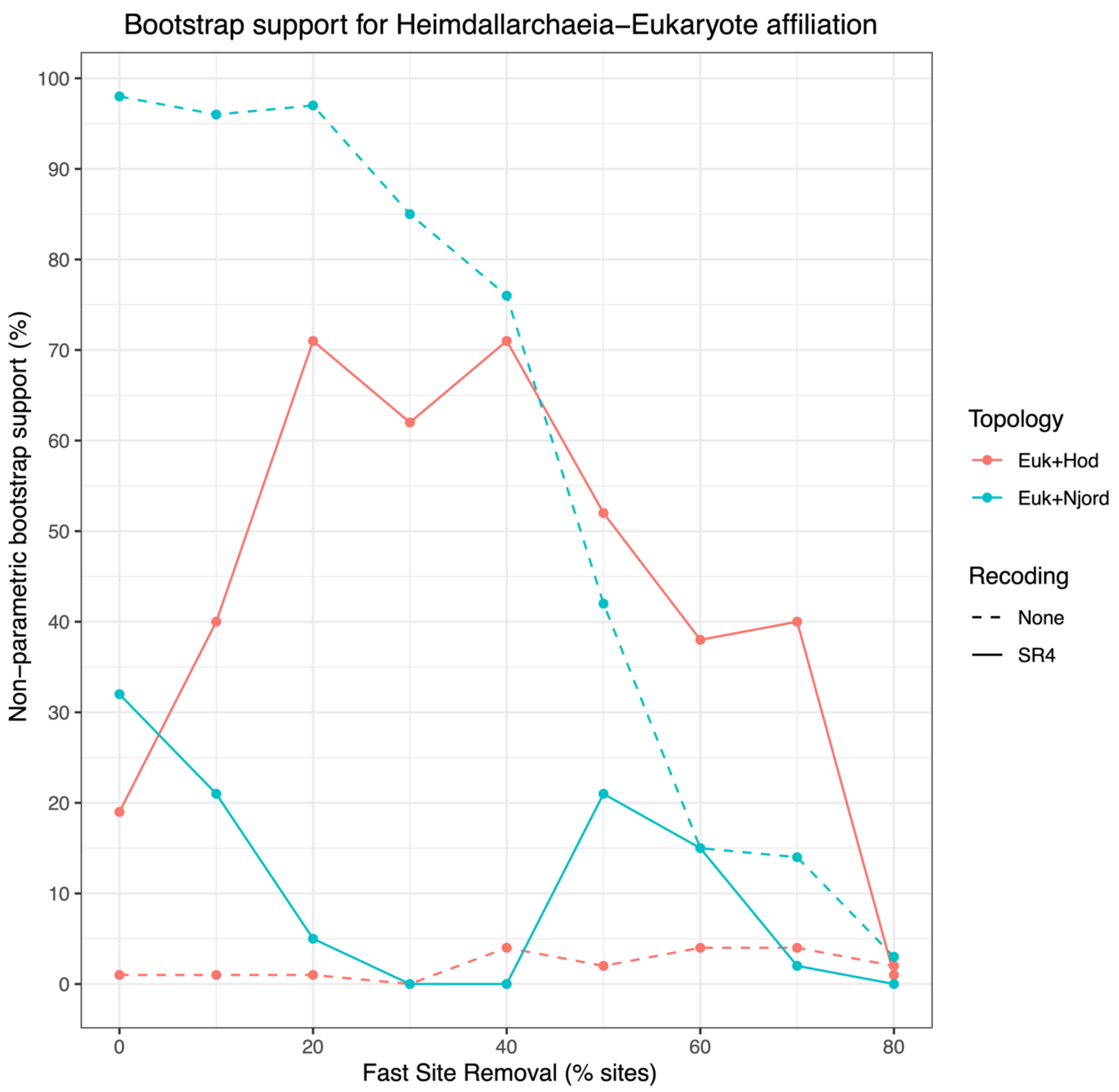


**Supplementary Figure 24.** Evolution of bootstrap support for the monophyly of eukaryotes and two distinct groups of Heimdallarchaeia: Hodarchaeales (red) and Njordarchaeales (blue), in phylogenies obtained from non-recoded (dashed) and SR4-recoded (full) alignments.

#### NM57-A175-nDK-SR4 (fastest 20% sites removed)

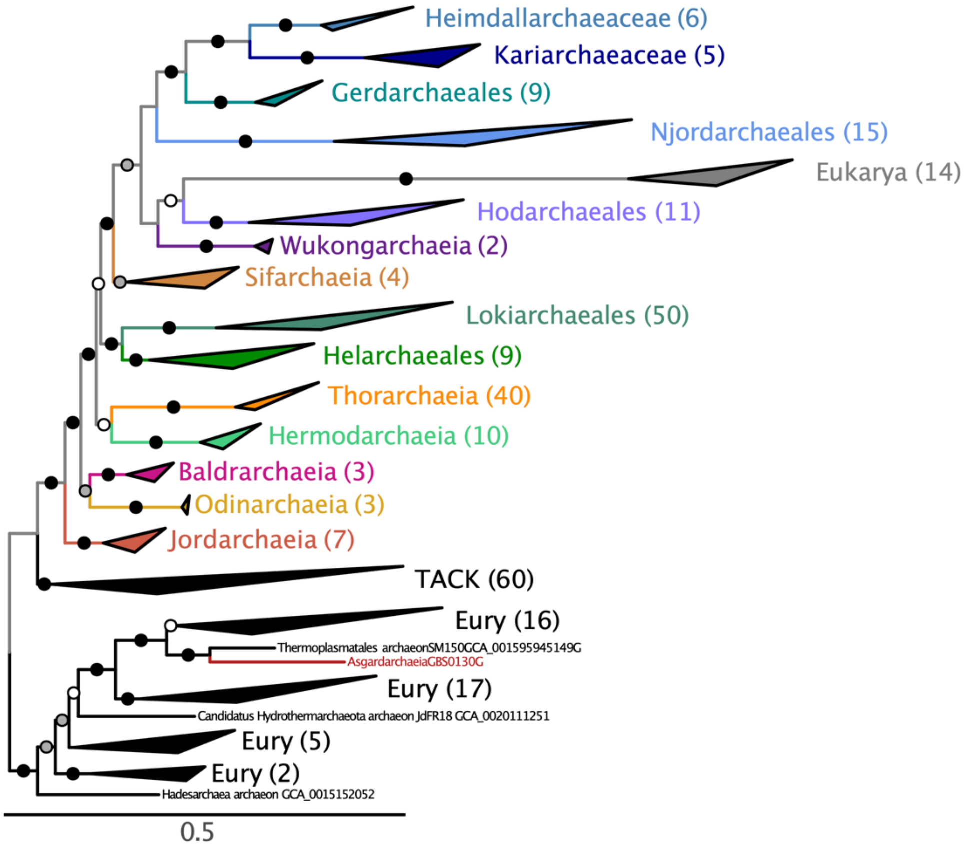


#### NM57-A175-nDK-SR4 (fastest 40% sites removed)

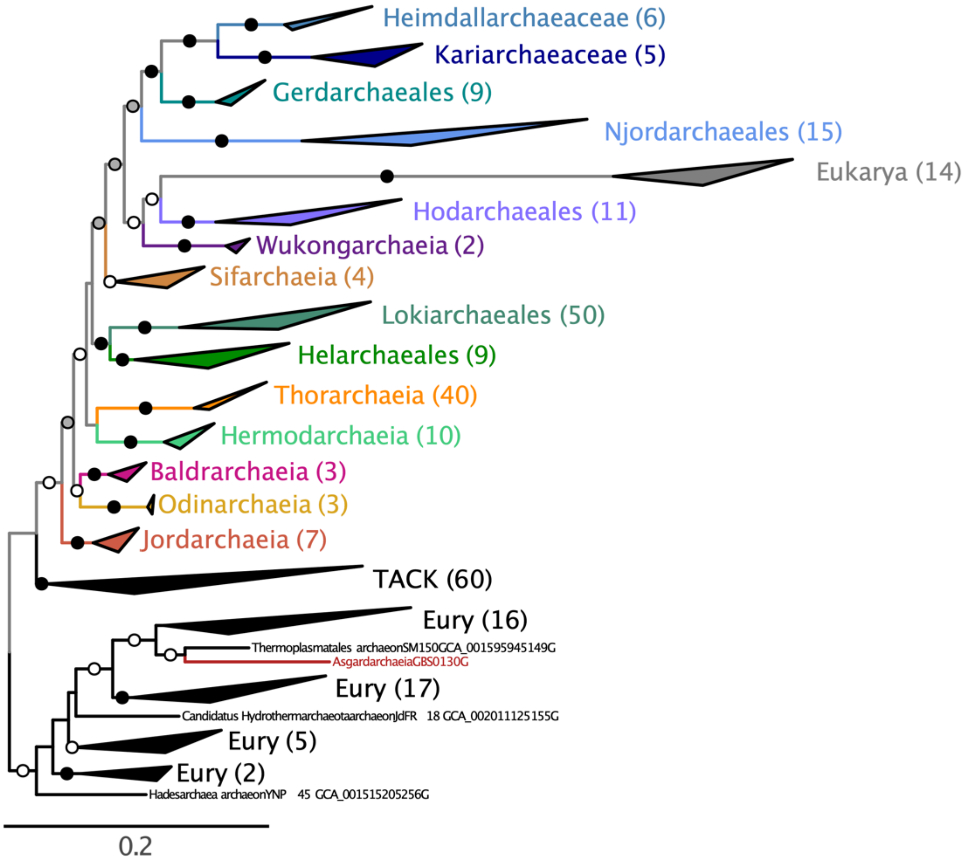

**Supplementary Figure 25.** Maximum-likelihood phylogenomic analysis based on the NM57-A175-nDK-SR4 with 20% and 40% of the fastest-evolving sites removed (292 taxa; 12584 and 9438 sites, respectively) using IQ-TREE under the under a user-defined ‘C60SR4’ model. DPANN and Korarchaeota have been discarded. Support at branches was estimated using the PMSF bootstrap approximation under the same model (100 pseudoreplicates). Black dots indicate maximum support values (100%), grey dots indicate bootstrap support 95-99% and white dots bootstrap support 70-95%. Tree is rooted on Euryarchaeota. Scale bar denotes the average expected substitutions per site. Uncollapsed phylogenies are available in Figshare.

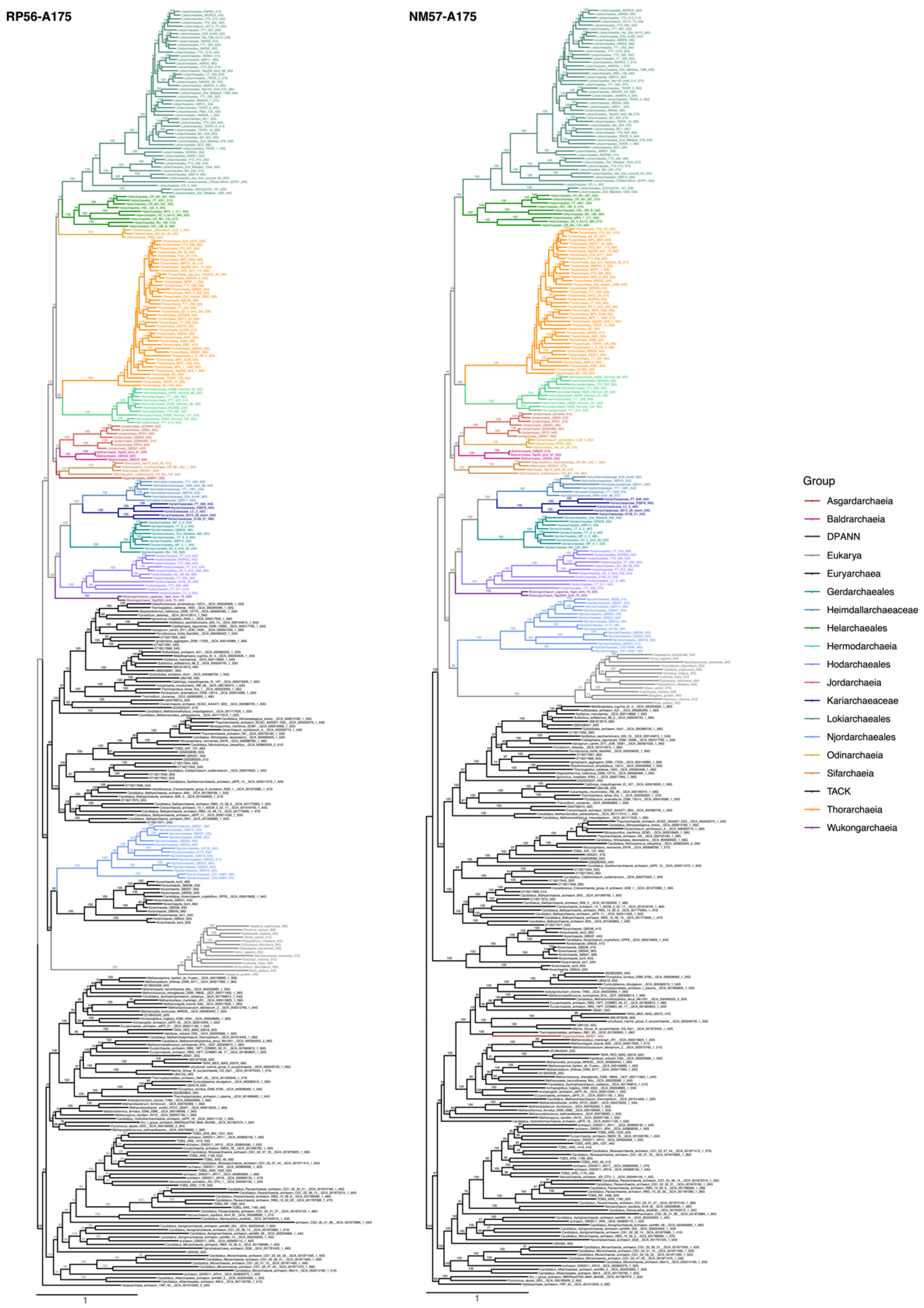


**Supplementary Figure 26.** Maximum likelihood phylogenomic analysis of the RP-A175 and NM-A175 dataset (7112 and 15733 sites after trimming, respectively; 345 taxa). Phylogeny was inferred using IQ-TREE under the LG+C60+F+Γ model. Support at branches was estimated using the PMSF bootstrap approximation under the same model (100 pseudoreplicates). Trees are midpoint rooted. Scale bar denotes the average expected substitutions per site. Uncollapsed trees are available in Figshare.

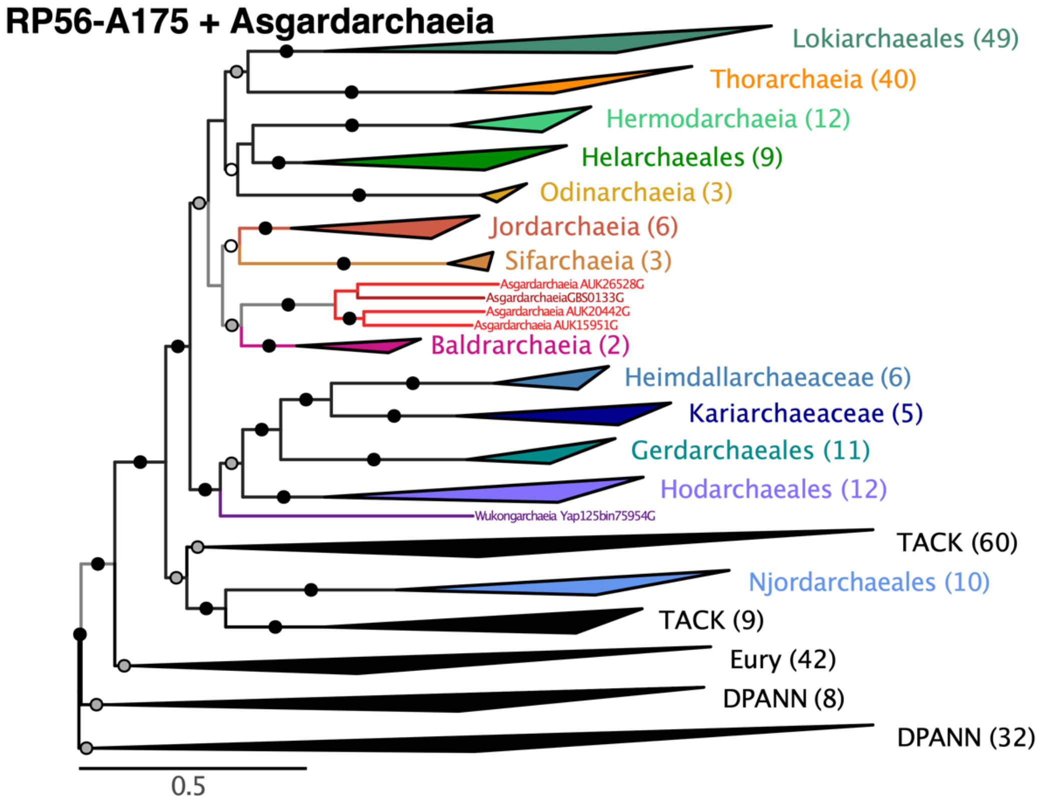
RP56-A175 + Asgardarchaeia
Lokiarchaeales (49)
Thorarchaeia (40)
Hermodarchaeia (12)
Helarchaeales (9)
Odinarchaeia (3)
Jordarchaeia (6)
Sifarchaeia (3)
Asgardarchaeia AUK26528G
AsgardarchaeiaGBS0133G
Asgardarchaeia AUK20442G
Asgardarchaeia AUK15951G
Baldrarchaeia (2)
Heimdallarchaeaceae (6)
Kariarchaeaceae (5)
Gerdarchaeales (11)
Hodarchaeales (12)
Wukongarchaeia Yap125bin75954G
TACK (60)
Njordarchaeales (10)
TACK (9)
Eury (42)
DPANN (8)
DPANN (32)
0.5

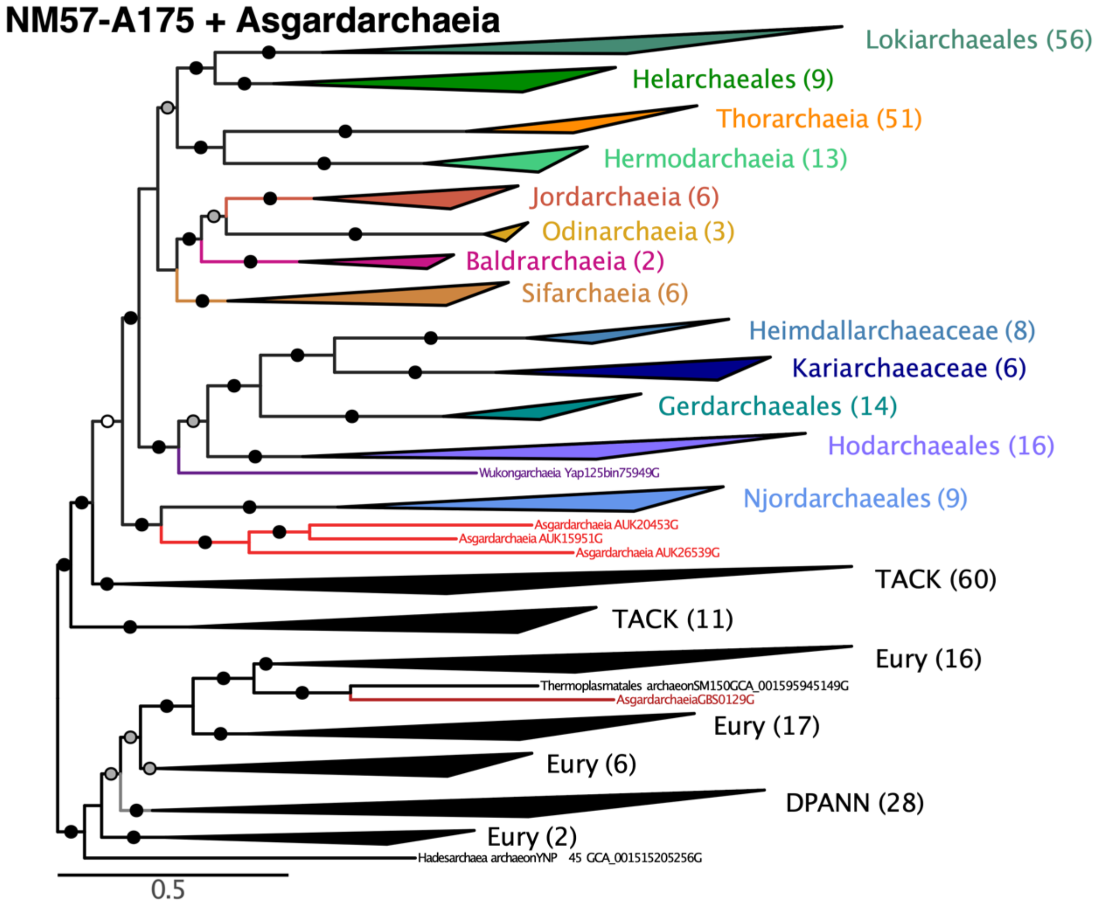
NM57-A175 + Asgardarchaeia
Lokiarchaeales (56)
Helarchaeales (9)
Thorarchaeia (51)
Hermodarchaeia (13)
Jordarchaeia (6)
Odinarchaeia (3)
Baldrarchaeia (2)
Sifarchaeia (6)
Heimdallarchaeaceae (8)
Kariarchaeaceae (6)
Gerdarchaeales (14)
Hodarchaeales (16)
Wukongarchaeia Yap125bin75949G
Njordarchaeales (9)
Asgardarchaeia AUK20453G
Asgardarchaeia AUK15951G
Asgardarchaeia AUK26539G
TACK (60)
TACK (11)
Eury (16)
Thermoplasmatales archaeonSM150GCA_001595945149G
AsgardarchaeiaGBS0129G
Eury (17)
Eury (6)
DPANN (28)
Eury (2)
Hadesarchaea archaeonYNP 45 GCA_001515205256G
0.5

**Supplementary Figure 27.** Maximum likelihood phylogenomic analysis of the RP56-A175 and NM57-A175 datasets, after including the protein sequences from three published strains (AUK159, AUK204 and AUK265), reducing the corresponding alignments by removing sequences 99% identical or more, and selecting taxa with 40 markers or more (8816 and 21393 sites after trimming, respectively; 469 and 451 taxa, respectively). Phylogeny was inferred using FastTree2 under the LG model with 20 rate categories. Trees are midpoint rooted. Scale bar denotes the average expected substitutions per site. Uncollapsed trees are available in Figshare.

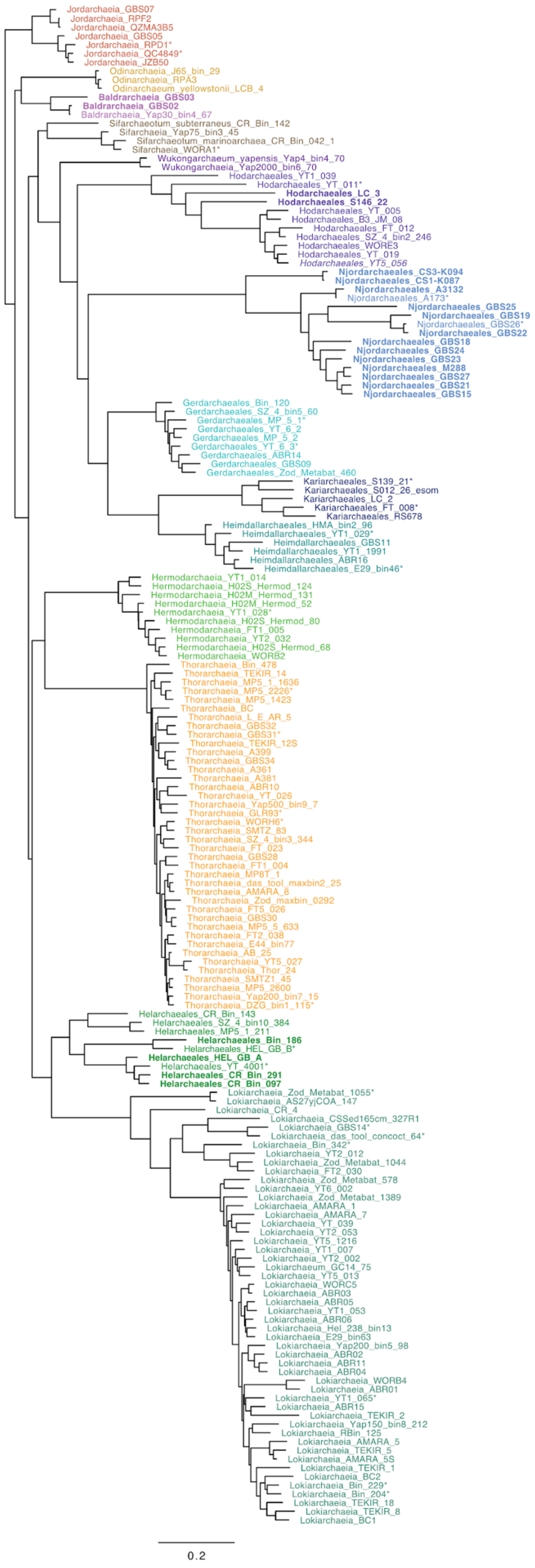
Jordarchaeia_GBS07
Jordarchaeia_RPF2
Jordarchaeia_QZMA3B5
Jordarchaeia_GBS05
Jordarchaeia_RPD1*
Jordarchaeia_QC4849*
Jordarchaeia_JZB50
Odinarchaeia_J65_bin_29
Odinarchaeia_RPA3
Odinarchaeum_yellowstonii_LCB_4
Baldrarchaeia_GBS03
Baldrarchaeia_GBS02
Baldrarchaeia_Yap30_bin4_67
Sifarchaeotum_subterraneus_CR_Bin_142
Sifarchaeia_Yap75_bin3_45
Sifarchaeotum_marinoarchaea_CR_Bin_042_1
Sifarchaeia_WORA1*
Wukongarchaeum_yapensis_Yap4_bin4_70
Wukongarchaeia_Yap2000_bin6_70
Hodarchaeales_YT1_039
Hodarchaeales_YT_011*
Hodarchaeales_LC_3
Hodarchaeales_S146_22
Hodarchaeales_YT_005
Hodarchaeales_B3_JM_08
Hodarchaeales_FT_012
Hodarchaeales_SZ_4_bin2_246
Hodarchaeales_WORE3
Hodarchaeales_YT_019
Hodarchaeales_YT5_056
Njordarchaeales_CS3-K094
Njordarchaeales_CS1-K087
Njordarchaeales_A3132
Njordarchaeales_A173*
Njordarchaeales_GBS25
Njordarchaeales_GBS19
Njordarchaeales_GBS26*
Njordarchaeales_GBS22
Njordarchaeales_GBS18
Njordarchaeales_GBS24
Njordarchaeales_GBS23
Njordarchaeales_M288
Njordarchaeales_GBS27
Njordarchaeales_GBS21
Njordarchaeales_GBS15
Gerdarchaeales_Bin_120
Gerdarchaeales_SZ_4_bin5_60
Gerdarchaeales_MP_5_1*
Gerdarchaeales_YT_6_2
Gerdarchaeales_MP_5_2
Gerdarchaeales_YT_6_3*
Gerdarchaeales_ABR14
Gerdarchaeales_GBS09
Gerdarchaeales_Zod_Metabat_460
Kariarchaeales_S139_21*
Kariarchaeales_S012_26_esom
Kariarchaeales_LC_2
Kariarchaeales_FT_008*
Kariarchaeales_RS678
Heimdallarchaeales_HMA_bin2_96
Heimdallarchaeales_YT1_029*
Heimdallarchaeales_GBS11
Heimdallarchaeales_YT1_1991
Heimdallarchaeales_ABR16
Heimdallarchaeales_E29_bin46*
Hermodarchaeia_YT1_014
Hermodarchaeia_H02S_Hermod_124
Hermodarchaeia_H02M_Hermod_131
Hermodarchaeia_H02M_Hermod_52
Hermodarchaeia_YT1_028*
Hermodarchaeia_H02S_Hermod_80
Hermodarchaeia_FT1_005
Hermodarchaeia_YT2_032
Hermodarchaeia_H02S_Hermod_68
Hermodarchaeia_WORB2
Thorarchaeia_Bin_478
Thorarchaeia_TEKIR_14
Thorarchaeia_MP5_1_1636
Thorarchaeia_MP5_2226*
Thorarchaeia_MP5_1423
Thorarchaeia_BC
Thorarchaeia_L_E_AR_5
Thorarchaeia_GBS32
Thorarchaeia_GBS31*
Thorarchaeia_TEKIR_12S
Thorarchaeia_A399
Thorarchaeia_GBS34
Thorarchaeia_A361
Thorarchaeia_A381
Thorarchaeia_ABR10
Thorarchaeia_YT_026
Thorarchaeia_Yap500_bin9_7
Thorarchaeia_GLR93*
Thorarchaeia_WORH6*
Thorarchaeia_SMTZ_83
Thorarchaeia_SZ_4_bin3_344
Thorarchaeia_FT_023
Thorarchaeia_GBS28
Thorarchaeia_FT1_004
Thorarchaeia_MP8T_1
Thorarchaeia_das_tool_maxbin2_25
Thorarchaeia_AMARA_8
Thorarchaeia_Zod_maxbin_0292
Thorarchaeia_FT5_026
Thorarchaeia_GBS30
Thorarchaeia_MP5_5_633
Thorarchaeia_FT2_038
Thorarchaeia_E44_bin77
Thorarchaeia_AB_25
Thorarchaeia_YT5_027
Thorarchaeia_Thor_24
Thorarchaeia_SMTZ1_45
Thorarchaeia_MP5_2600
Thorarchaeia_Yap200_bin7_15
Thorarchaeia_DZG_bin1_115*
Helarchaeales_CR_Bin_143
Helarchaeales_SZ_4_bin10_384
Helarchaeales_MP5_1_211
Helarchaeales_Bin_186
Helarchaeales_HEL_GB_B*
Helarchaeales_HEL_GB_A
Helarchaeales_YT_4001*
Helarchaeales_CR_Bin_291
Helarchaeales_CR_Bin_097
Lokiarchaeia_Zod_Metabat_1055*
Lokiarchaeia_AS27yjCOA_147
Lokiarchaeia_CR_4
Lokiarchaeia_CSSed165cm_327R1
Lokiarchaeia_GBS14*
Lokiarchaeia_das_tool_concoct_64*
Lokiarchaeia_Bin_342*
Lokiarchaeia_YT2_012
Lokiarchaeia_Zod_Metabat_1044
Lokiarchaeia_FT2_030
Lokiarchaeia_Zod_Metabat_578
Lokiarchaeia_YT6_002
Lokiarchaeia_Zod_Metabat_1389
Lokiarchaeia_AMARA_1
Lokiarchaeia_AMARA_7
Lokiarchaeia_YT_039
Lokiarchaeia_YT2_053
Lokiarchaeia_YT5_1216
Lokiarchaeia_YT1_007
Lokiarchaeia_YT2_002
Lokiarchaeum_GC14_75
Lokiarchaeia_YT5_013
Lokiarchaeia_WORC5
Lokiarchaeia_ABR03
Lokiarchaeia_ABR05
Lokiarchaeia_YT1_053
Lokiarchaeia_ABR06
Lokiarchaeia_Hel_238_bin13
Lokiarchaeia_E29_bin63
Lokiarchaeia_Yap200_bin5_98
Lokiarchaeia_ABR02
Lokiarchaeia_ABR11
Lokiarchaeia_ABR04
Lokiarchaeia_WORB4
Lokiarchaeia_ABR01
Lokiarchaeia_YT1_065*
Lokiarchaeia_ABR15
Lokiarchaeia_TEKIR_2
Lokiarchaeia_Yap150_bin8_212
Lokiarchaeia_RBin_125
Lokiarchaeia_AMARA_5
Lokiarchaeia_TEKIR_5
Lokiarchaeia_AMARA_5S
Lokiarchaeia_TEKIR_1
Lokiarchaeia_BC2
Lokiarchaeia_Bin_229*
Lokiarchaeia_Bin_204*
Lokiarchaeia_TEKIR_18
Lokiarchaeia_TEKIR_8
Lokiarchaeia_BC1
0.2

1984

**Supplementary Figure 28. Mapping of RNA-dependent DNA polymerase A (RpoA) gene structure onto the Asgard species tree.** Bold font indicates species which encode the fused version of RpoA. We could not identify both part of RpoA for the taxa marked with an asterisk, indicating that their RpoA gene structure is unknown. The species tree was inferred using Phylobayes on the NM-nDEK SR4-recoded dataset, and pruned from Euryarchaeota and TACK representatives for this figure.

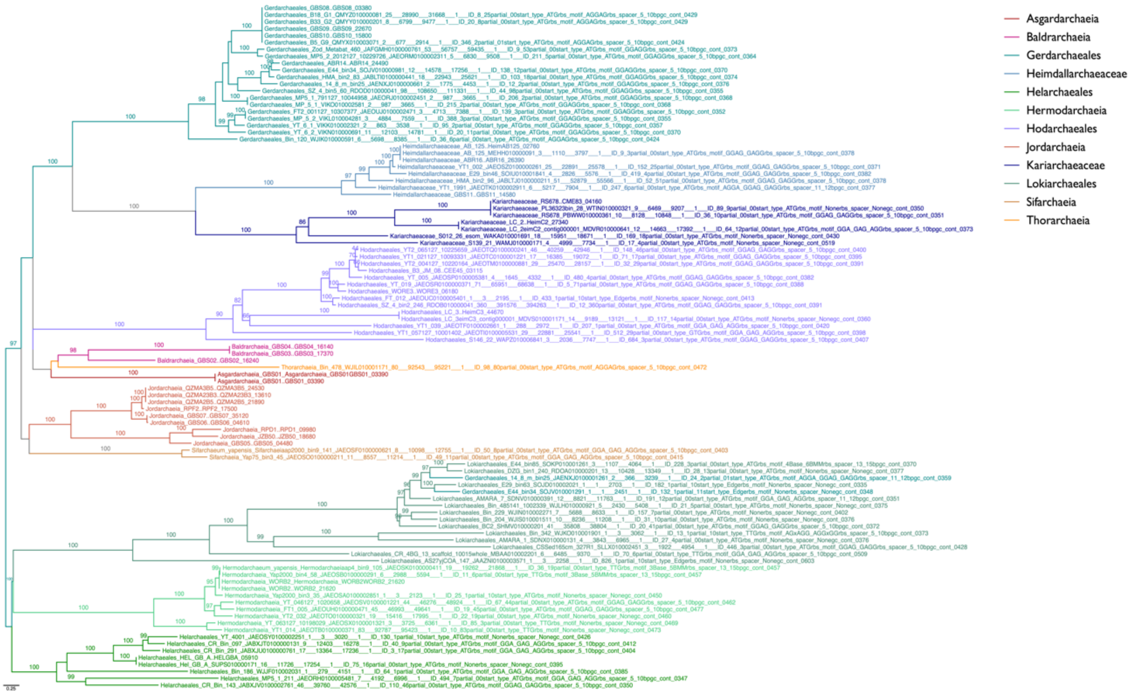


**Supplementary Figure 29.** Maximum likelihood phylogeny of the RNA polymerase A, subunit A’ in Asgard archaeal representatives. The tree was inferred from 99 sequences (849 sites) using IQ-TREE under the LG+C20+Γ model. Support at branches was estimated with 1000 ultrafast bootstrap replicates. Only support values >90% are displayed. Note the monophyly of all hodarchaeales homologs. Scale bar denotes the average expected substitutions per site.

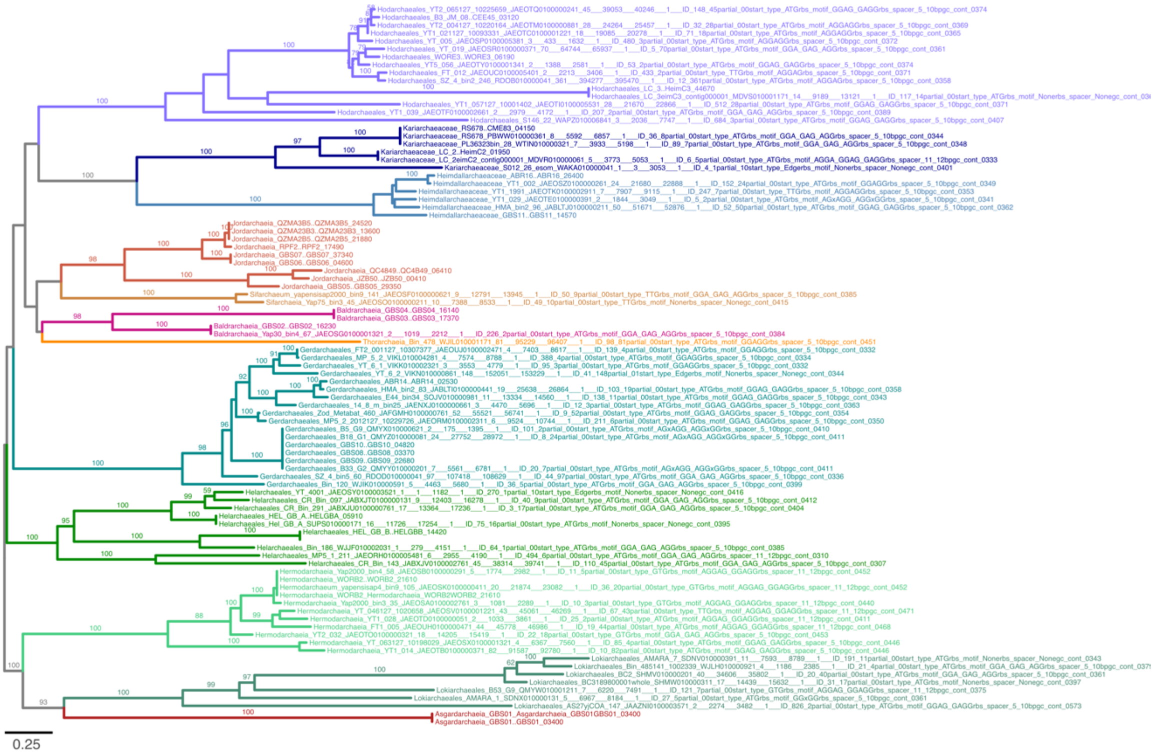


**Supplementary Figure 30.** Maximum likelihood phylogeny of the RNA polymerase A, subunit A’’ in Asgard archaeal representatives. The tree was inferred from 91 sequences (365 sites) using IQ-TREE under the LG+C20+Γ model. Support at branches was estimated with 1000 ultrafast bootstrap replicates. Only support values >90% are displayed. Note the monophyly of all hodarchaeales homologs. Scale bar denotes the average expected substitutions per site.

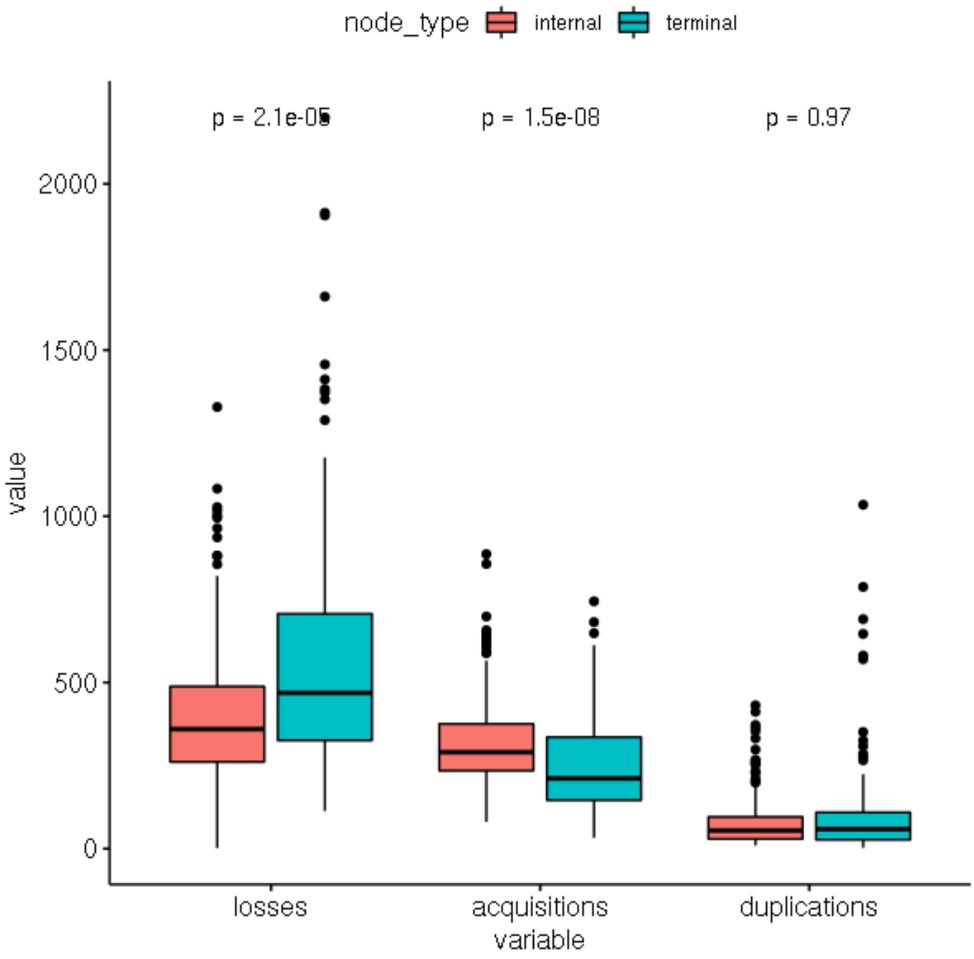


**Supplementary Figure 31. Total number of events inferred per Asgard node.** Boxplots show the distribution of the number of losses, duplications, and acquisitions (originations + transfers) inferred at internal (i.e., ancestors) and terminal (i.e., extant organisms) nodes. Wilcoxon test p-values for each pairwise comparison are shown on top. The higher number of inferred losses and lower number of inferred gains at terminal nodes compared to internal nodes likely result in part from MAG incompleteness. Indeed, because missing data is randomly distributed across MAGs, and because we are using a version of ALE taking into account the percentage of missing data in each genome by modeling a different loss rate in each terminal branch compared to the rest of the tree, inferences made at internal nodes are reliable.

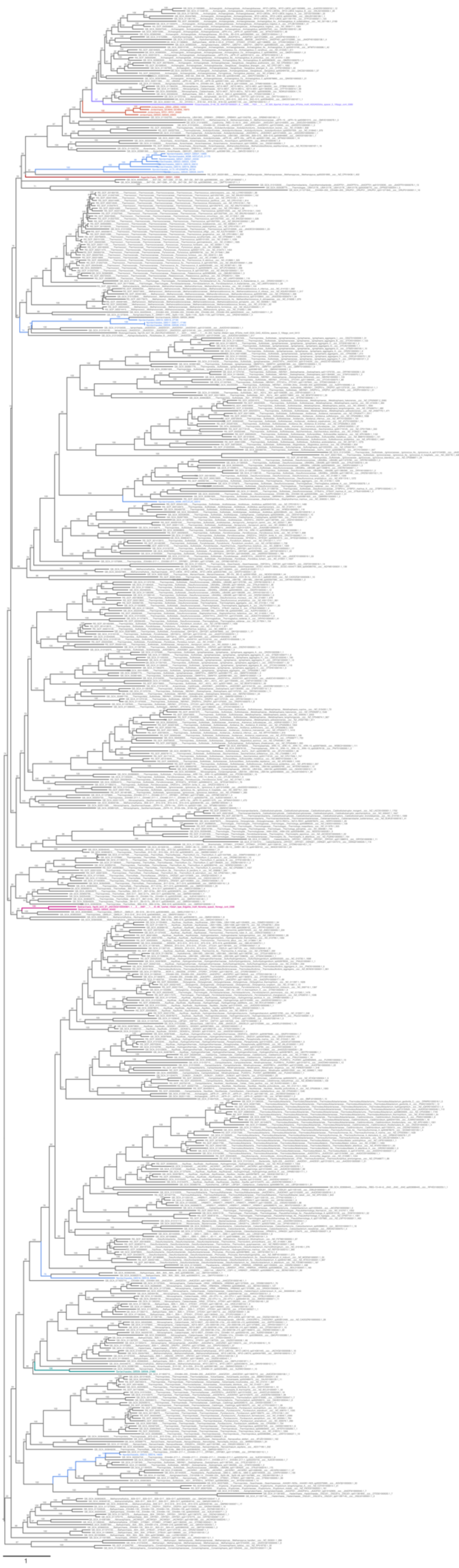

2028

**Supplementary Figure 32**. Maximum likelihood phylogeny of the toprim reverse gyrase homologs in archaeal representatives. The tree was inferred from 594 sequences (630 sites) using IQ-TREE under the LG+C60+Γ model. Support at branches was estimated with 1000 ultrafast bootstrap replicates. Tree is midpoint rooted. Scale bar denotes the average expected substitutions per site.

### Supplementary tables

**Supplementary Table 1. MAGs metadata.** The first and second tabs contain sampling sites metadata for all Asgard archaea MAGs included in this work: newly described and previously published, respectively. The third tab contains statistics related to all Asgard archaea MAGs included in this work evaluated with CheckM[39]. The fourth tab contains taxonomy information for each MAG based on phylogenomic analyses. Every MAG is classified at the phylum level, and Heimdallarchaeia MAGs are classified at the class level. Additionally, lower taxonomy ranks are included for *Candidatus t*ype species. A description of the taxonomy names is discussed in Supplementary discussion, section 1: "Description of new taxa". Bolded lines correspond to the MAGs described as part of this study. N/A: Not available.

**Supplementary Table 2. Phylogenomic analyses summary.** Statistical support for the monophyly of various clades is provided as PMSF bootstrap support values for ML analyses, and as percentage of sampled trees across each chain for the Phylobayes analyses (after burn-in). The "new markers" are identified and annotated as in Petitjean et al. (2015)[100]. Data treatment is indicated: SR4: SR4-recoding; FSR: Fast-site removal. The number listed after "FSR" indicates the percentage of sites removed (e.g., FSR10 means the 10% fastest evolving sites were removed). The monophyly tested is indicated by brackets around the relevant group(s). Groups are abbreviated as follows. H: Heimdallarchaeia; Ho: Hodarchaeales; Ge: Gerdarchaeales; Ka: Kariarchaeaceae; Hei: Heimdallarchaeaceae; Nj: Njordarchaeales; Asg: Asgard archaea; Asg_nH: All Asgard archaea minus Heimdallarchaeia; Asg_nN: All Asgard archaea minus Njordarchaeales; E: Eukaryotes ; K: Korarchaeota; TAC: Thaum-, Aig- and Crenarchaeota; L: Lokiarchaeia; Lo: Lokiarchaeales; Hel: Helarchaeales; T: Thorarchaeia; O: Odinarchaeia; J: Jordarchaeia; B: Baldrarchaeia; Her: Hermodarchaeia; A: Asgardarchaeia; S: Sifarchaeia; W: Wukongarchaeia. An "n" before a group name means that they are excluded from the monophyly tested. For example, the column "(Asg_nNj)" reports the support for the monophyly of all Asgard archaea to the exclusion of Njordarchaeales.

**Supplementary Table 3. Accession numbers of ESP homologs in Asgard archaea.** Each candidate is associated with the Interpro domains ID and description obtained with Interproscan, as well as its homologous cluster ID and associated arCOG annotation.

**Supplementary Table 4. Annotation and copy number of genes in selected Asgard ancestral nodes.** A gene was considered present if the inferred copy number in a given ancestral node was above 0.3 (filled circles). A gene was considered as possibly present (half-filled circle) if the copy number was between 0.1 and 0.3. The gene was considered absent (empty circle) when the copy number was below 0.1. Protein annotation of key enzymatic steps discussed throughout this manuscript was manually verified. Proteins potentially in central metabolic pathways are functionally grouped and numbered from 1-29. The description of those 29 pathways is shown in Sheet 1. Enzymes discussed in the text and/or with unusual distributions in the Asgard ancestors are highlighted in light yellow.

**Supplementary Table 5. Optimal Growth Temperature predictions.** OGT were predicted for the genomes presented here based on genomic and proteomic features[11]. Since nucleotide fractions of the ribosomal RNAs are used in this method, only those genomes with predicted rRNAs could be used.

**Supplementary Table 6. Statistical analyses for thermophily-related compositional bias.** t-test statistical analyses of thermophily-related amino acid compositional features between Korarchaeota, Njordarchaeales, Hodarchaeales, and the group formed by Gerdarchaeales, Heimdallarchaeaceae and Kariarchaeaceae. For each performed t-test, we Bonferroni-corrected p-values (p-values lower than 2.2e-16 are highlighted). Tests were performed using the t.test function in R.

| | Completeness | Contamination | Acquisitions | Duplications | Losses |
|---|---|---|---|---|---|
| **Completeness** | | -0.62*** | 0.25*** | 0.17* | -0.54*** |
| **Contamination** | | | -0.11 | 0.14 | 0.39*** |
| **Acquisitions** | | | | 0.62*** | 0.07 |
| **Duplications** | | | | | 0.15* |

**Supplementary Table 7. Correlations between the number of evolutionary events inferred per terminal node and the estimated completeness and contamination values.** Pairwise correlations between the inferred number of duplications, losses, acquisitions and the estimated contamination and completeness values according to CheckM. Excluding the highly redundant genome of Lokiarchaeum CG14_75 did not have a significant effect. Coefficients and p-values for each Spearman correlation are shown. P-values are represented by asterisks where *: p-value <= 0.05, **: p-value <=0.01, ***: p-value <=0.001.

**Supplementary Table 8. Inferred values for proteome size, and for gene transfer, origination, duplication, and loss events in ancestral reconstruction analysis.** Values are indicated per node number (or terminal node) as indicated in Supplementary Figure 12.

### Supplementary data

**Supplementary Data 1. Predicted proteomes for Asgard archaeal MAGs and protein clusters used as input for inferring ancestral genome content will be released on Figshare (https://figshare.com/account/home#/projects/111912) upon publication of the manuscript.**
